# *recount-brain*: a curated repository of human brain RNA-seq datasets metadata

**DOI:** 10.1101/618025

**Authors:** Ashkaun Razmara, Shannon E. Ellis, Dustin J. Sokolowski, Sean Davis, Michael D. Wilson, Jeffrey T. Leek, Andrew E. Jaffe, Leonardo Collado-Torres

## Abstract

The usability of publicly-available gene expression data is often limited by the availability of high-quality, standardized biological phenotype and experimental condition information (“metadata”). We released the *recount2* project, which involved re-processing ∼70,000 samples in the Sequencing Read Archive (SRA), Genotype-Tissue Expression (GTEx), and The Cancer Genome Atlas (TCGA) projects. While samples from the latter two projects are well-characterized with extensive metadata, the ∼50,000 RNA-seq samples from SRA in *recount2* are inconsistently annotated with metadata. Tissue type, sex, and library type can be estimated from the RNA sequencing (RNA-seq) data itself. However, more detailed and harder to predict metadata, like age and diagnosis, must ideally be provided by labs that deposit the data.

To facilitate more analyses within human brain tissue data, we have complemented phenotype predictions by manually constructing a uniformly-curated database of public RNA-seq samples present in SRA and *recount2*. We describe the reproducible curation process for constructing *recount-brain* that involves systematic review of the primary manuscript, which can serve as a guide to annotate other studies and tissues. We further expanded *recount-brain* by merging it with GTEx and TCGA brain samples as well as linking to controlled vocabulary terms for tissue, Brodmann area and disease. Furthermore, we illustrate how to integrate the sample metadata in *recount-brain* with the gene expression data in *recount2* to perform differential expression analysis. We then provide three analysis examples involving modeling postmortem interval, glioblastoma, and meta-analyses across GTEx and TCGA. Overall, *recount-brain* facilitates expression analyses and improves their reproducibility as individual researchers do not have to manually curate the sample metadata. *recount-brain* is available via the `add_metadata()` function from the *recount* Bioconductor package at <u>bioconductor.org/packages/recount</u>.

## Introduction

RNA sequencing (RNA-seq) is a valuable method for measuring gene expression across the transcriptome. The widespread availability and falling cost of high-throughput sequencing has lead to massive amounts of biological information being accumulated in public data repositories (Denk, 2017). The Sequence Read Archive (SRA) – which was established in 2007 by the National Center for Biotechnology Information (NCBI) and functions as the National Institute of Health’s (NIH) primary archive of high-throughput sequencing data – hosts raw sequencing data for >50,000 human RNA-seq samples (Leinonen et al., 2011) and is rapidly expanding (Kodama et al., 2012; Langmead and Nellore, 2018). This is a tremendous resource that allows genomic researchers to answer biological questions using already-sequenced reads from other laboratories. Deposition of data in the SRA is mandated by most funding agencies and open access journals, resulting in an expansive range of biological samples. However missing information on biological phenotype (sex, age, disease status and type) and experimental condition (library selection, brain bank), in short sample metadata, significantly reduces its utility to researchers (Langmead and Nellore, 2018). In fact, critical sample phenotype information is missing or incomplete for many samples (92.7%) within the SRA (Ellis et al., 2018), limiting their ability to answer biological questions using gene expression data.

Because the SRA is made up of individual submissions, data are not provided in a consistent format and annotations (such as methodology and technical sequencing details) are often unclear or missing (Bernstein et al., 2017). We previously developed *recount2* (Collado-Torres et al., 2017c, 2017a), a public resource with over 70,000 uniformly processed human RNA-seq samples, enabling comparisons across studies for human expression data in the public repository. However, inconsistent phenotype annotation still remains a barrier to taking advantage of public uniformly processed reads. Several efforts have been made to improve the sample metadata for SRA samples by predicting metadata from abstracts (Kingsford, 2016), automatically normalizing the available metadata with ontologies (Bernstein et al., 2017), and predicting sample metadata from expression values (Ellis et al., 2018). Here, we describe a reproducible curation process that complements phenotype predictions and automatic ontology inference. This curation process can be adapted and applied to other studies and tissues, thus expanding the use of the public RNA-seq data.

We applied our reproducible curation process to create *recount-brain*, a freely available human brain sample metadata database for SRA samples present in *recount2* with unified metadata variables for brain Genotype-Tissue Expression (GTEx) (GTEx Consortium, 2015) and The Cancer Genome Atlas (TCGA) samples (Cancer Genome Atlas Research Network, 2008; Hutter and Zenklusen, 2018). By accessing the sample metadata and expression data via the *recount* Bioconductor package at <u>bioconductor.org/packages/recount</u> (Collado-Torres et al., 2017b, 2017c) researchers may study transcriptomic changes in neurological diseases. Moreover, we offer a streamlined and efficient curation method for researchers and students to contribute critical sample metadata and enhance reproducibility in genomics. We outline our curation process here – the insights gained and lessons learned – so that others may reproduce similar results or analyze public human brain RNA-seq data in a fraction of the time.

## Results

### Ready to use human brain sample metadata

*recount-brain* hosts sample metadata for 4,431 human brain tissue samples from 62 projects from the SRA, out of which 3,214 (72.5%) samples have expression data available from *recount2* (Collado-Torres et al., 2017c). *recount-brain* supports powerful search and filter capabilities by tissue phenotype, including spanning 3,600 neurological controls with 2,900 from the SRP025982 study (SEQC/MAQC-III Consortium, 2014); 15 neurological diseases and information on brain tumor subtype, grade, and stage; 3 levels of detailed anatomic tissue site information; 5 developmental stages (Fetus, Infant, Child, Adolescent, Adult); demographic data (age, sex, race); technical sequencing information (RIN, PMI, sequencing layout, library source, etc.); and Brodmann area, tissue and disease ontologies (Methods: Reproducible curation process, Ontology mapping). The SRA samples in *recount-brain* are complemented by 1,409 GTEx (GTEx Consortium, 2015) and 707 TCGA (Brennan et al., 2013; Cancer Genome Atlas Research Network et al., 2015) samples covering 13 healthy regions of the brain and 2 tumor types, respectively (Methods: Merging recount-brain with GTEx and TCGA). In total, there are 6,547 samples with metadata in *recount-brain* with 5,330 (81.4%) present in *recount2* (Collado-Torres et al., 2017c) and the curation process is reproducible. Of the samples present in *recount-brain*, 58.7% are absent from MetaSRA (Bernstein et al., 2017) brain samples and conversely MetaSRA lists samples absent from *recount-brain* showcasing how these approaches complement each other (Methods: MetaSRA comparison). **Figure 1** outlines the variables that were used and a list of sample attributes found in the database. The complete list of variables and descriptions is available in **Table S1**.

**Figure 1.**
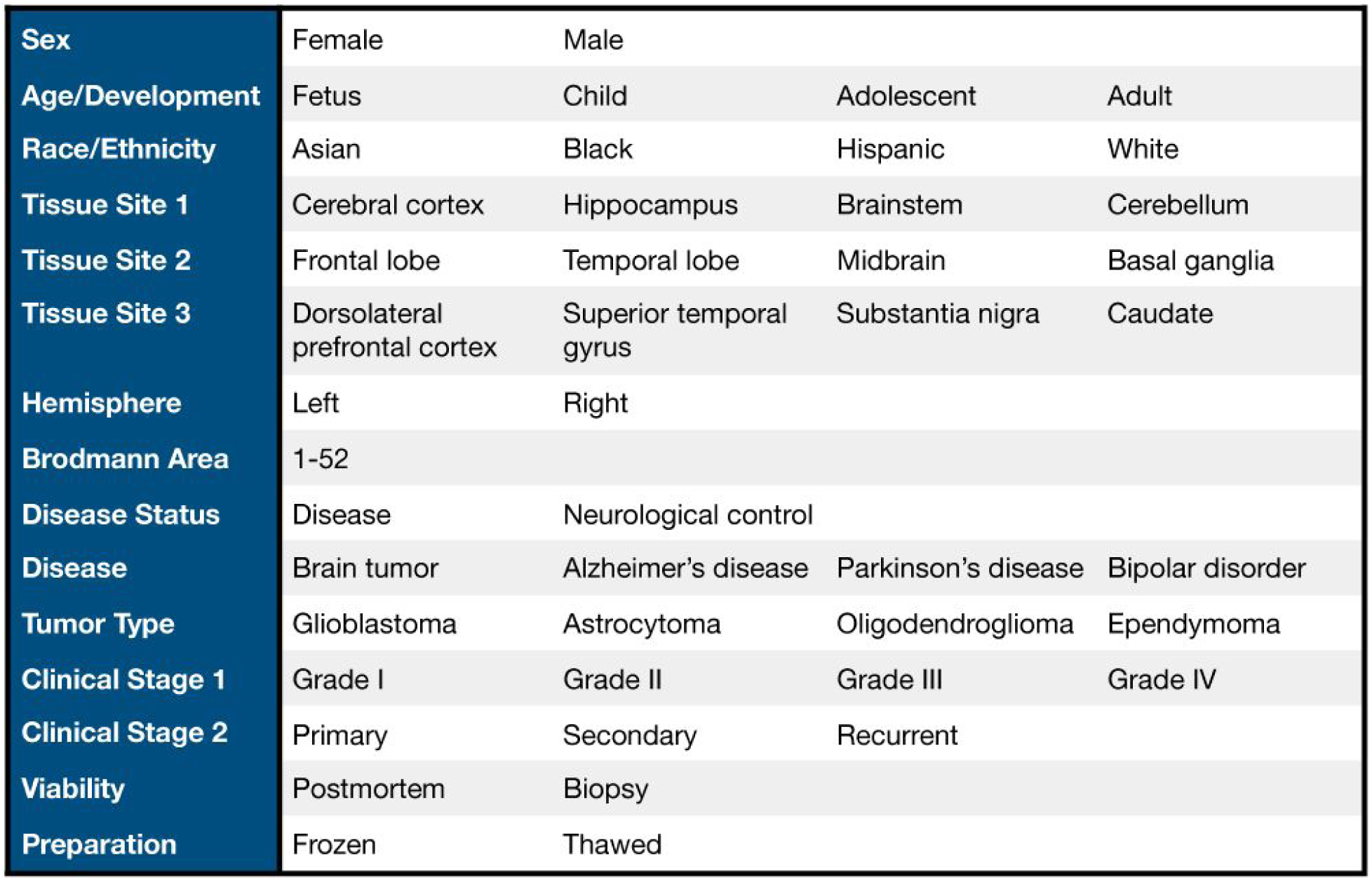
*recount-brain* example metadata. Example variables (blue column) and values of the data present in *recount-brain*. The full list of sample metadata variables and descriptions is available in **Table S1**.

The `add_metadata()`function in *recount* (Collado-Torres et al., 2017b) makes it easy to access the complete *recount-brain* metadata in RNA-seq analyses. This function can be used to access the full *recount-brain* metadata to find samples and studies of interest as illustrated in **Figure 2** (purple box). Alternatively, users can interactively explore *recount-brain* via https://jhubiostatistics.shinyapps.io/recount-brain/ to identify samples of interest (Methods: Interactive display). Once a study of interest has been identified, the user can download the gene expression data from *recount2* (Collado-Torres et al., 2017c) and append the *recount-brain* metadata as shown in **Figure 2**; this process is equivalent to appending custom metadata from Figure 2 of the *recount workflow* (Collado-Torres et al., 2017a). Once the expression data from *recount2* and the sample metadata from *recount-brain* have been combined, the user can proceed to perform analyses such as identification of differentially expressed genes and enriched gene ontologies, examples of them are illustrated in **Figure 2** with data from SRA study SRP027383 (Bao et al., 2014).

**Figure 2.**
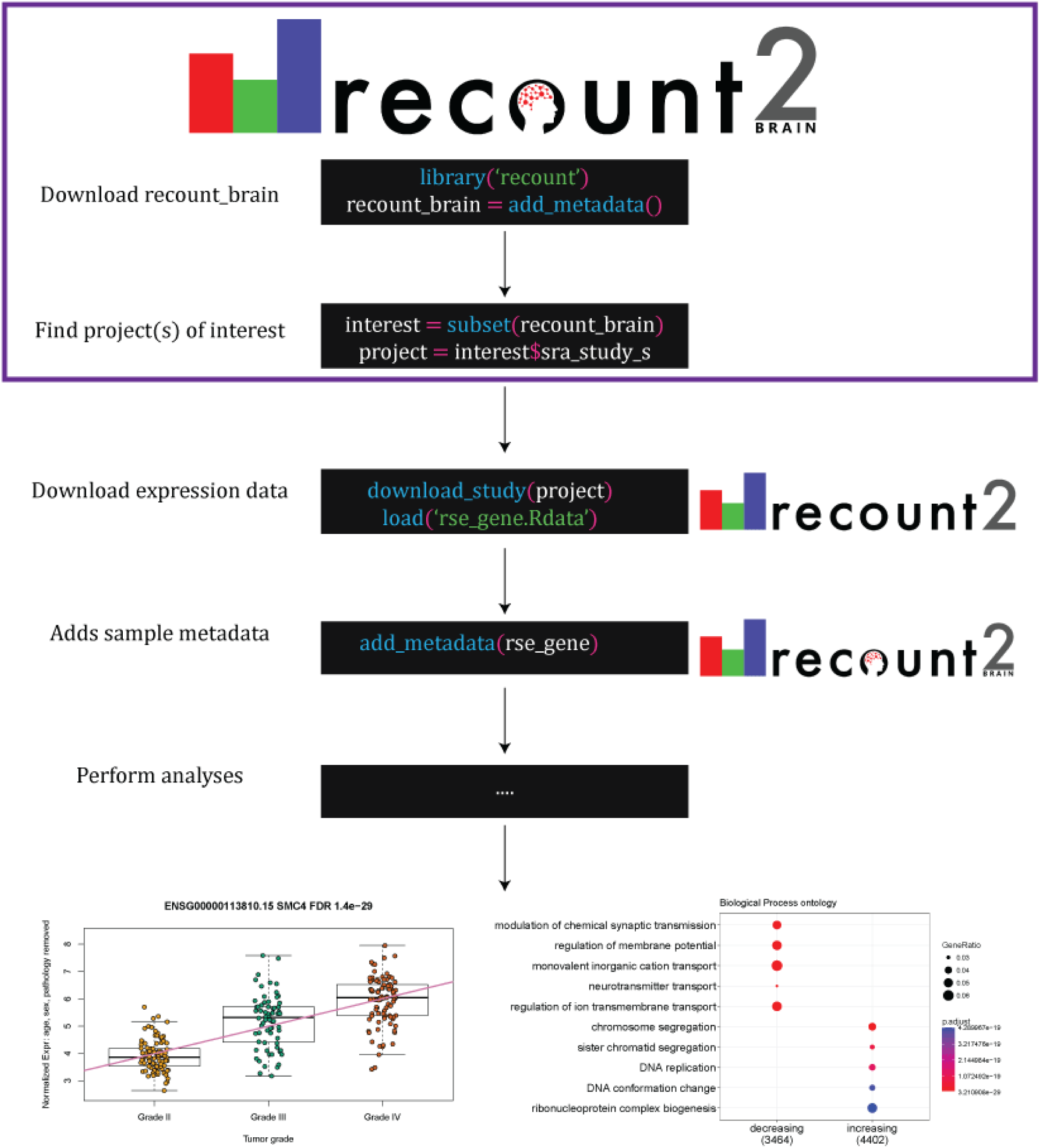
Uses of *recount-brain* and its relationship with *recount2*. *recount-brain* facilitates identifying project(s) of interest (purple box) programmatically or interactively through https://jhubiostatistics.shinyapps.io/recount-brain/. After downloading expression data from *recount2*, *recount-brain* can enrich the sample metadata for brain studies. This information can be used to perform analyses to find differentially expressed genes and enriched gene sets such as those exemplified with SRA Study SRP027383 (Bao et al., 2014), where the top differentially expressed gene among glioblastoma samples in *recount2* is *SCM4*. Black boxes represent R code with functions highlighted in blue, input arguments in green, and R objects in white.

### Differential gene expression analysis using *recount-brain*

To exemplify how *recount-brain* facilitates re-analysis of public human brain RNA-seq data, we selected an SRA study with almost no sample information available from the NCBI SRA Run Selector (Vera Alvarez et al., 2017) (Figure 1 of **Supplementary File 1**), studySRP027383 with data for 272 gliomas (Bao et al., 2014). Using the gene expression data from study SRP027383, we identified 6,116 and 6,438 genes with significant (FDR <1%) decreasing and increasing expression associations with linear tumor grade progression (II, III and IV), while adjusting for sex, age and pathology (IDH1 mutation status) using the 258 (94.9%) samples that had complete sample metadata (Methods: Differential expression by tumor grades with data from SRP027383). *SMC4*, the top differentially expressed gene (**Figure 2**) plays a role in the structural maintenance of chromosomes. The genes with an increase expression as tumor grade progresses are enriched for DNA replication and chromosome segregation biological processes (**Figure 2**). *SMC4* is a core component of the condensin complexes which has recently been associated with aggressive glioblastoma phenotypes (Jiang et al., 2017). Furthermore, *SMC4* mRNA and protein expression levels are associated with poor prognosis and could potentially be a therapeutic target in gliomas (Jiang et al., 2017).

The full code for reproducing this example analysis is available in **Supplementary File 1** and at http://LieberInstitute.github.io/recount-brain/. Subject-matter experts could further examine the results and guide analyses like the one we carried out. Without *recount-brain* and *recount2* (Collado-Torres et al., 2017c) for this analysis one would have had to process the raw expression data and obtain the relevant sample metadata, which would likely have taken a considerable amount of time. Furthermore, the processed RNA-seq data from *recount2*, the curated sample metadata from *recount-brain* and the analysis code provided in **Supplementary File 1** are all public resources that enable the full reproducibility of the analysis we described.

### Effects of post-mortem interval on transcription

Researchers with their own human brain datasets can use *recount-brain* to assess the replication of their results, regardless of the publication status of their projects. Recently Ferreira *et al*. investigated the impact of post-mortem interval (PMI) on gene expression using data from multiple tissues of post-mortem healthy donors obtained from the GTEx project (Ferreira et al., 2018). To exemplify how *recount-brain* can be used to replicate findings, we identified 10 SRA studies with variable PMI data for a total of 252 samples from 9 publications (Boudreau et al., 2014; Dumitriu et al., 2016; Khrameeva et al., 2014; Labadorf et al., 2015; Li et al., 2014; Magistri et al., 2015; Pardo et al., 2013; Voineagu et al., 2011; Wu et al., 2012). In their analysis of GTEx and PMI data, Ferreira *et al*. identified genes with significant temporal changes across tissues (Ferreira et al., 2018). Among them, Ferreira *et al*. (their Figure 2D) found no significant association between *RNASE2*, *EGR3*, *HBA1* and *CXCL2* expression and PMI interval in the brain cerebellum or cortex (Ferreira et al., 2018). We replicated their findings for *EGR3*, *HBA1* and *CXCL2* but found a significant decrease in *RNASE2* expression with PMI interval progression across the four intervals we had data for (**Figure 3**, Methods: Effects of post-mortem interval on transcription). This significant decrease in *RNASE2* expression as PMI increases is in line with Ferreira’s *et al*. findings for the 13 other tissues they examined (Ferreira et al., 2018). A sensitivity analysis using the 4 studies with samples spanning at least 3 PMI intervals (SRP019762, SRP048683, SRP051844, SRP058181) revealed increased cross-study variability in *RNASE2* expression through PMI intervals (Figures 9 to 12 from **Supplementary File 2**) which could be why Ferreira *et al*. observed no significant decrease in *RNASE2* expression in the human brain. Having multiple studies in *recount2* enables researchers to perform this type of sensitivity analyses.

**Figure 3.**
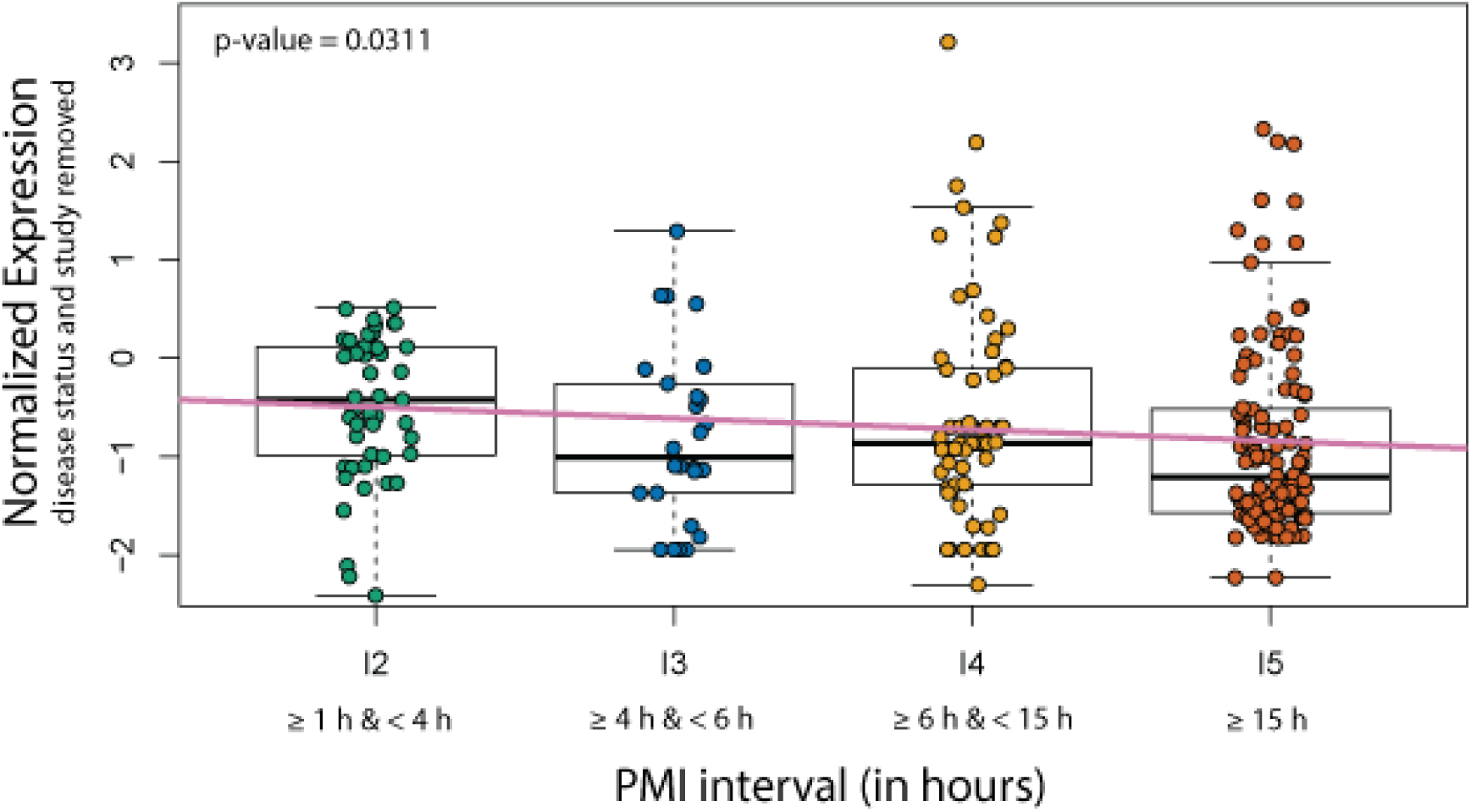
RNASE2 expression across post-mortem intervals. Normalized *RNASE2* expression, log2 (RPKM + 0.5), after controlling for disease status and study across 4 PMI intervals for 252 samples from 10 SRA studies. We had no observations for the first interval (< 1 h) that Ferreira *et al*. used in their Figure 2D (Ferreira et al., 2018).

Ferreira *et al*. assessed the impact of PMI in energy metabolism by investigating its relationship with mitochondrial expression levels (Figures 4B, 4C and Supplementary Figure 17) (Ferreira et al., 2018) and observed that certain tissues, such as brain cerebellum and cortex, exhibited a positive relationship between PMI and mitochondrial transcription, while other tissues demonstrated a negative relationship (Ferreira et al., 2018). We replicate this finding here by taking advantage of the fact that *recount2* provides base-pair coverage data in the form of BigWig files (Collado-Torres et al., 2017a, 2017c). We used this BigWig data to compute the percent of mitochondrial transcription and observed a positive association with PMI (in hours, **Figure 4A**). We also computed the percent of transcription that overlaps genes from Gencode v25 (Harrow et al., 2012) excluding the mitochondrial genes. We expected a negative association between mitochondrial and global gene transcription; however, we found instead that the 10 studies clustered into two distinct sets (**Figure 4B**). These sets could not be explained by disease status or PMI (Figures 14 and 15 from **Supplementary File 2**). The two sets of studies are set 1: SRP017933, SRP032539, SRP032540, SRP048683, SRP056604 (Boudreau et al., 2014; Li et al., 2014; Magistri et al., 2015; Pardo et al., 2013); set 2: ERP001304, SRP051844, SRP007483, SRP058181, SRP019762 (Dumitriu et al., 2016; Khrameeva et al., 2014; Labadorf et al., 2015; Voineagu et al., 2011; Wu et al., 2012). Once we considered the study set membership, we found no significant association between PMI and mitochondrial transcription as well as no significant difference in the change in mitochondrial transcription by a unit increase in PMI (the interaction effect): only a significant difference for the intercept (**Figure 4C**). We examined differences between the two sets of studies across disease status, demographic, biological, technical, quality covariates as well as SHARQ beta (Kingsford, 2016) and *phenopredict* sample metadata (Ellis et al., 2018) and found no clear cut differences. Most of the samples from the second study set are 100 base-pair paired-end reads and have a 0.63 higher mean RIN (0.35 to 0.92 95% CI, p-value 4.804e-05). However, these differences don’t predict the two type of relationships observed in **Figure 4B**. Overall, our replication analysis revealed that two types of associations between mitochondrial and genome transcription (excluding mitochondrial genes) that could change the interpretation of the implications of PMI in energy metabolism. The full code for reproducing this example analysis is available in **Supplementary File 2** and at http://LieberInstitute.github.io/recount-brain/.

**Figure 4.**
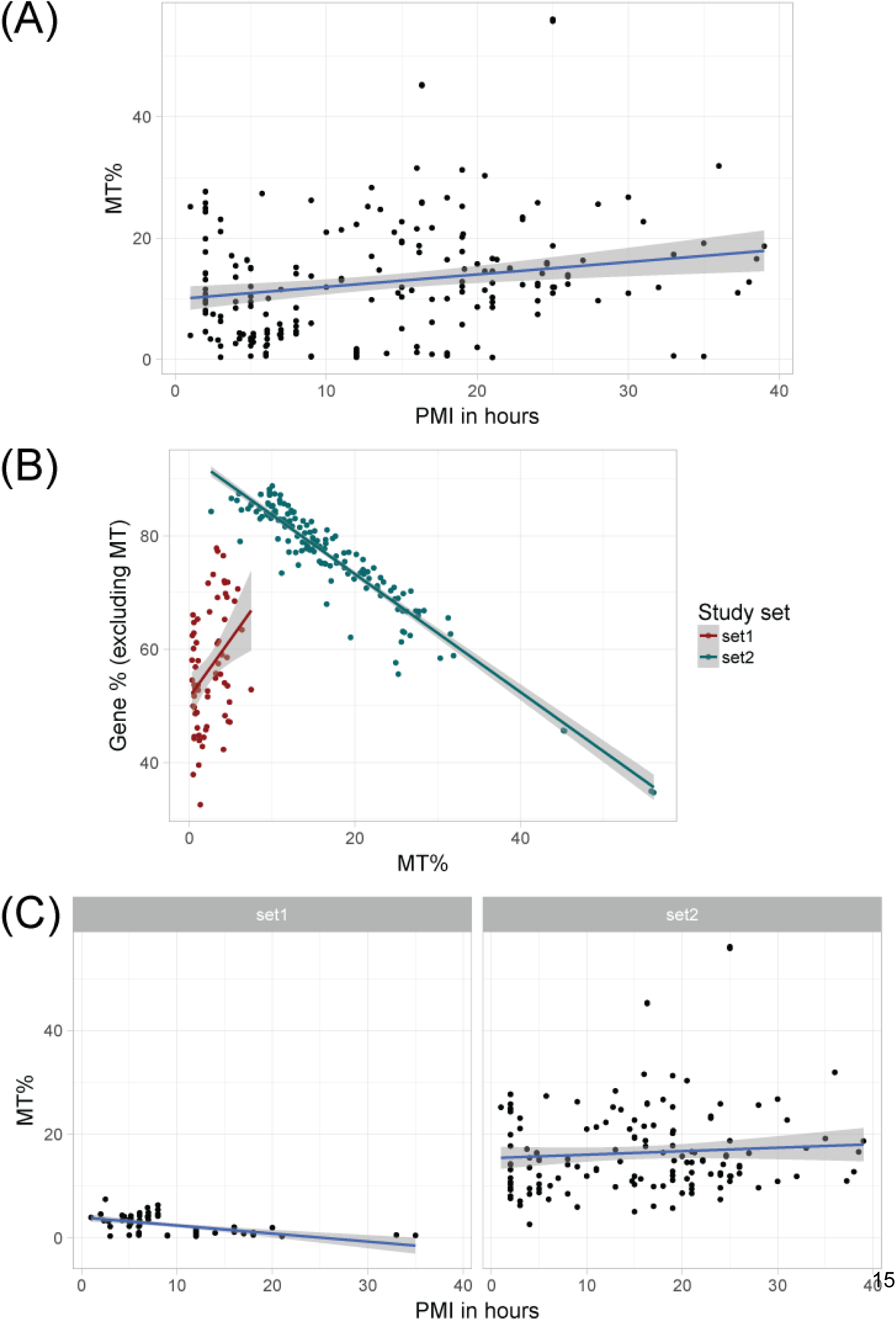
Association between mitochondrial transcription and post-mortem interval. (**A**) Percent of mitochondrial transcription is significantly associated (p < 0.05) with PMI in hours across 252 samples from 10 studies. (**B**) Percent of mitochondrial transcription and gene transcription (excluding mitochondrial genes) are negatively associated for samples from study set 2, but not for samples from study set 1. (**C**) PMI in hours does not have a significant association with mitochondrial transcription once samples are separated by study set and the trend is not significantly different between study sets.

### Variation in gene expression across multiple glioblastoma studies

Two large studies present in *recount2* (Collado-Torres et al., 2017c), GTEx (GTEx Consortium, 2015) and TCGA (Brennan et al., 2013; Hutter and Zenklusen, 2018), have more detailed sample metadata than the rest of the SRA samples. We thus expect that users will be interested in combining the human brain SRA samples from *recount-brain* with either GTEx or TCGA. To exemplify this process, we selected the two glioblastoma studies with at least 20 samples: SRP027383 (Bao et al., 2014) and SRP044668 (Gill et al., 2014). We then combined these SRA samples with those from TCGA listed as primary glioblastoma tumors (Brennan et al., 2013; Cancer Genome Atlas Research Network, 2008). Using only the tumor samples we normalized the studies and removed variation across them to make them comparable (Methods: Variation in gene expression across multiple glioblastoma studies). Once the studies were on a comparable scale (**Figure 5 A**) we computed the variance for each gene for each of these three studies. The 1,000 most variable genes in glioblastoma were highly concordant across the three studies (**Figure 5 B**). The tight concordance is more readily noticeable when comparing TCGA primary tumor data from the brain (Brennan et al., 2013) and kidney (Davis et al., 2014) (**Figure 5 B**).

**Figure 5.**
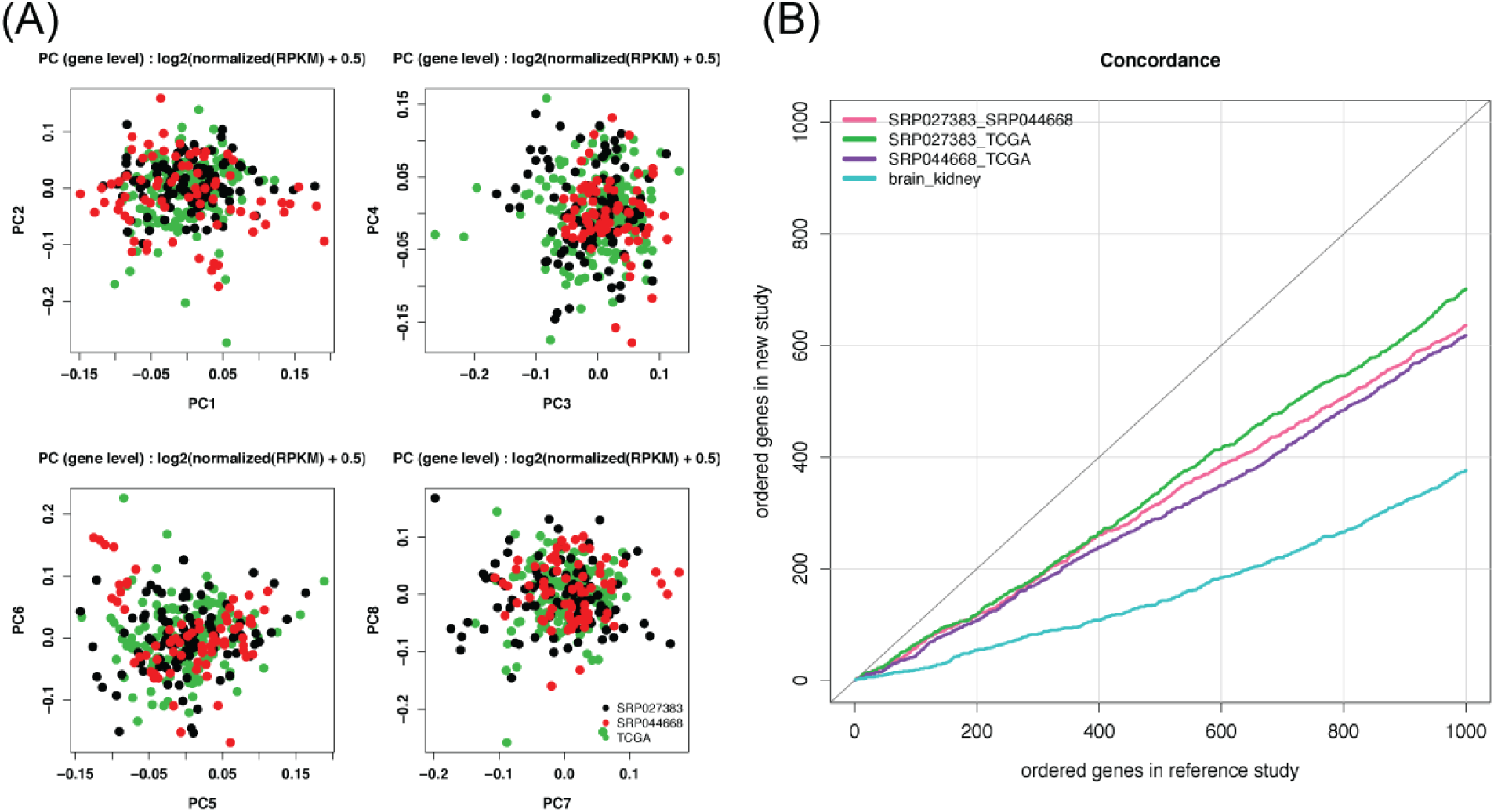
Gene ranking by variability consistency in glioblastoma studies. (**A**) Top 8 principal components (PCs) for the normalized RPKM in log2(x + 0.5) scale for the tumor samples from three glioblastoma studies: SRP027383 (black), SRP044668 (red) and TCGA (green). (**B**) Pairwise concordance comparisons for the top 1,000 genes ranked by their variability in each study. Pink: SRP027383 versus SRP044668, green: SRP027383 versus TCGA brain, purple: SRP044668 versus TCGA brain, blue: TCGA brain vs TCGA kidney.

This example analysis demonstrates how *recount-brain* can be used with TCGA data and facilitate meta-analyses. To further simplify the process of comparing SRA, GTEx and TCGA human brain data, we adapted the GTEx and TCGA human brain sample metadata and merged it into *recount-brain* (see Methods). The full code for reproducing this example analysis is available in **Supplementary File 3** and at http://LieberInstitute.github.io/recount-brain/.

## Discussion

Massive amounts of sequencing data are accumulating in public repositories, but unlabeled or unannotated variables limit the ability of researchers to analyze these data (Langmead and Nellore, 2018). We present *recount-brain*, a freely available human brain sample metadata database that pairs with the uniformly-processed RNA-seq data from *recount2* enabling researchers to study transcriptomic changes in neurological disease. Our metadata database and reproducible curation approach shows that freely-available data can be cleaned and curated to encourage and facilitate data reuse and increase the value of small studies, such as SRP017933 (Pardo et al., 2013), and large studies, such as GTEx and TCGA (Hutter and Zenklusen, 2018), alike.

In our methods, we described a semi-automated reproducible process through the pseudo-algorithm provided (**Figure 6**) that offers a streamlined, efficient method for researchers interested in genomics to contribute critical data and enhance reproducibility in the field. Prior experience and knowledge are useful but not necessary as the curation process has been distilled down to an easy to replicate step-by-step method. We envision future students, who may have a particular interest in a disease or organ system, collaborating with genomic data scientists to learn about the field in an interactive hands-on way while simultaneously contributing valuable and publishable work. Improving the metadata of public data will benefit everyone and facilitate the creation of major data search engines including the new public data Google website (Castelvecchi, 2018) described at https://www.blog.google/products/search/making-it-easier-discover-datasets/. The *recount-brain* metadata we curated is readily available via the `add_metadata()` functionality in the *recount* Bioconductor package (Collado-Torres et al., 2017b) available at <u>bioconductor.org/packages/recount</u>.

**Figure 6.**
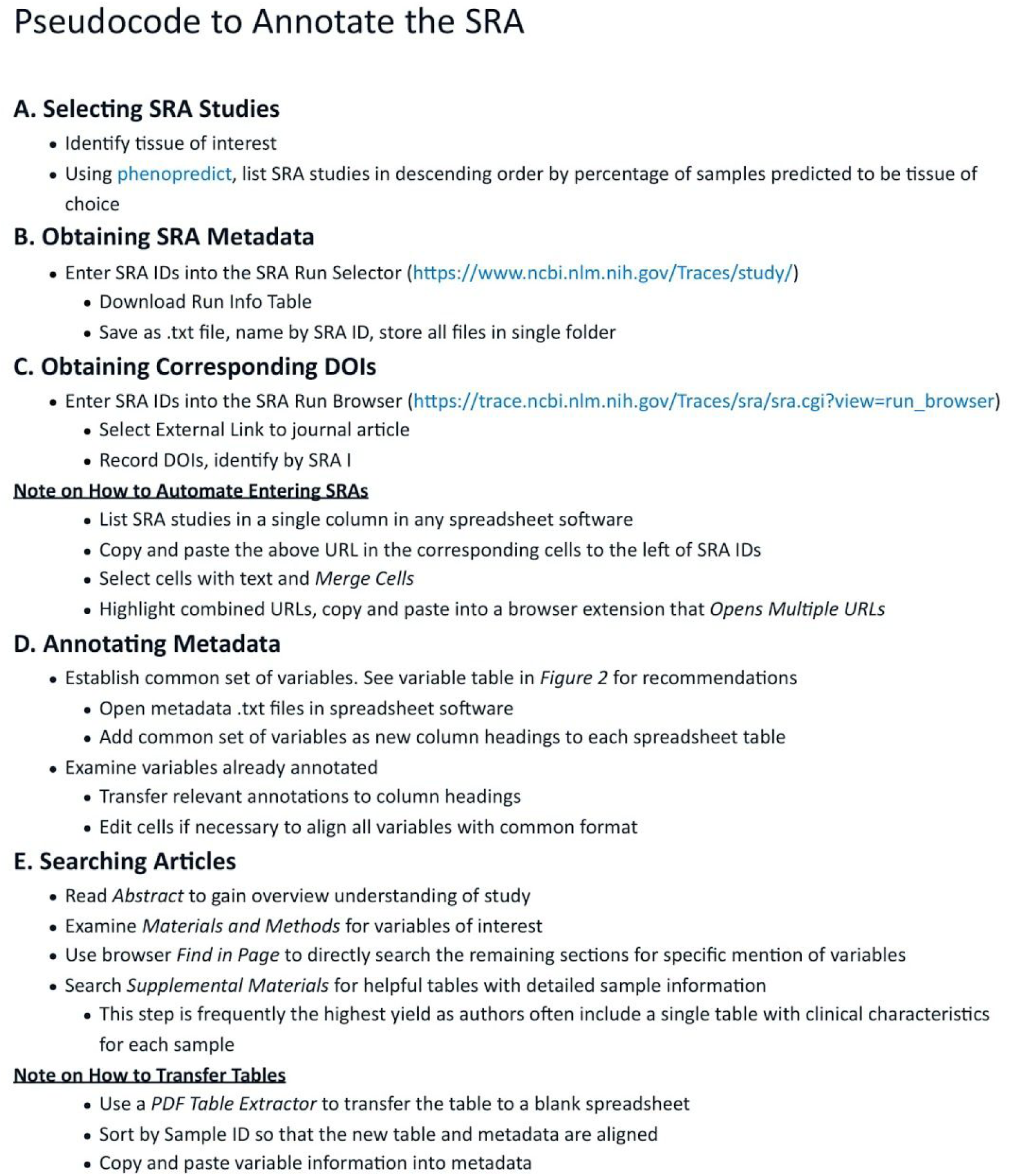
Pseudocode to annotate human samples from the Sequence Read Archive (SRA) present in *recount2*. This annotation workflow can be applied to other tissues by selecting a candidate projects using predictions by *phenopredict* (Ellis et al., 2018) or other efforts (Bernstein et al., 2017; Kingsford, 2016).

Enabling researchers to take advantage of deposited data will increase reproducibility in genomics research as “taking no notice of deposited data is similar to ignoring several independently published replication experiments” (Denk, 2017). Our refined database and reproducible model reuses public data, enhances reproducibility among genomic researchers, and enables translational discovery. *recount-brain* further improves the usability of the RNA-seq data present in *recount2* (Collado-Torres et al., 2017c). We showed how one can combine the two resources to perform analyses, explore different biological questions, replicate findings and expand analyses from other studies. *recount-brain* also facilitates meta-analyses across SRA, GTEx, and TCGA samples such as our example analysis on the expression variability in glioblastoma tumor samples.

The sample metadata in *recount-brain* can be combined with other projects that enhance *recount2* beyond gene expression. For example, Snaptron (Wilks et al., 2018) provides rapid querying of splice junctions and splicing patterns from samples in *recount2* and could be used together with *recount-brain* for studying human brain splicing. Furthermore, *recount-brain* metadata could be utilized together with the recently released transcript abundance estimates (Fu et al., 2018) using the same `add_metadata()` functionality that we demonstrated with our example use cases. The `add_metadata()` function can be easily expanded to cover other tissues if others follow our annotation workflow (**Figure 6**) to improve the sample metadata of other studies. Furthermore, curation efforts such as *recount-brain* will likely be useful in the creation of more refined sample metadata prediction algorithms (Ellis et al., 2018). Curation efforts complement prediction and automatic ontology inference approaches as there is always some uncertainty attached to predictions and inferences (Bernstein et al., 2017; Ellis et al., 2018).

We documented our curation process with detailed notes and code at http://LieberInstitute.github.io/recount-brain/. We believe that *recount-brain* will save time for other researchers since they can immediately bypass this time-consuming process: from extracting the information from the articles to merging the variables with GTEx and TCGA, as well as matching with ontology databases. By using the R package versioned framework and constructing a unified resource we are also promoting reproducibility of downstream analyses. Furthermore, we invite researchers to contribute to *recount-brain* and the `add_metadata()` framework in *recount* by curating more human datasets and submitting them via https://github.com/LieberInstitute/recount-brain/issues/new. We envision that researchers will follow similar curation processes to ours or compute sample metadata predictions and contribute them to the community via sample metadata unification projects.

## Methods

### Reproducible curation process

A summary of the curation process we followed is shown by the pseudo-algorithm in **Figure 6**. First, it was necessary to select the studies that would eventually compose *recount-brain*. For the 50,099 human RNA-seq SRA samples for which *recount2* has expression data, we used the v0.0.03 tissue predictions from the *phenopredict* R package (Ellis et al., 2018) to list all studies predicted to have brain tissue samples (**Figure 6A**). We then identified all SRA studies that were made up of >70% brain tissue samples and had at least 4 samples to increase our yield of total brain samples for our refined database. We downloaded study metadata from the NCBI SRA Run Selector (https://www.ncbi.nlm.nih.gov/Traces/study/), identified each study’s associated journal publications through the NCBI SRA Run Browser (https://trace.ncbi.nlm.nih.gov/Traces/sra/sra.cgi?view=run_browser) (Vera Alvarez et al., 2017), and obtained the corresponding articles and supplementary materials via NCBI PubMed (NCBI Resource Coordinators, 2018). Metadata for all studies were saved as tables and identified by SRA Study ID (**Figure 6B-C**).

Based on an exploratory analysis of the common annotated variables across included studies, we developed a common set of variables that we believed would be most useful to investigators for downstream analyses (**Figure 6D**). We then carefully and systematically searched article text and supplementary materials for specific information on the biological samples. Novel tissue attributes found in publication text but not included in the original metadata were added directly to the project-specific metadata tables. These included demographic data, technical sequencing information, clinical and pathological characteristics, and anatomical details. If downloaded metadata already included data for one or more of our uniform set of variables, then we aligned it with our uniform set of variables.

The most important aspect of curation is the search, identification, and transfer of tissue attributes not included in the original metadata. It is also, by far, the most time-consuming, which is why we have described our recommendations for the process (**Figure 6E**). The key was to develop a structured and systematic approach to each study, which allowed us to gather the most information possible while ensuring reproducibility. Our search process focused on *Materials and Methods* and *Supplementary Materials* sections in the articles we curated. Some of the time, we were able to locate a comprehensive table describing sample characteristics and clinical information about the donor. However, when no table existed, we relied on combing through the article text to find specific descriptions about tissue samples and how they were prepared. Our findings were then transferred the metadata spreadsheet one by one. We adhered strictly to a uniform method for annotation and documentation.

A thorough reproducibility document details the exact location (heading, section, text, table, and/or figure) where sample information was found. The full reproducibility document is available at http://LieberInstitute.github.io/recount-brain/ and is organized by SRA Study. Updated metadata tables were saved as CSV files and merged across projects into a single table *(recount-brain-v1*). We introduced the `add_metadata()` function to the *recount* (version >= 1.5.6) Bioconductor package (Collado-Torres et al., 2017b, 2017c) to facilitate merging the expression data from *recount2* with the sample metadata from *recount-brain*.

### Merging *recount-brain* with GTEx and TCGA

We combined the *recount-brain-v1* with the brain metadata from TCGA (Brennan et al., 2013; Cancer Genome Atlas Research Network, 2008; Cancer Genome Atlas Research Network et al., 2015) and GTEx (GTEx Consortium, 2015) and created *recount-brain-v2*. We found 707 and 1,409 brain samples in TCGA and GTEx, respectively. The metadata in TCGA and GTEx was relatively complete; however, the formats differed between them and from *recount-brain-v1*. We compiled the TCGA and GTEx metadata and converted them into the format used in *recount-brain-v1* when creating *recount-brain-v2*. Some variables, such as the “Brain tissue repository source” were directly combined between the three datasets; however, most involved some minor alterations or were not comparable. For example, the “Nature of Disease (Disease / Control)” variable informs if a sample is a “Solid Tissue Normal” or a tumour of any type (i.e primary or recurrent) in TCGA, and was adapted from the Death Classification: 4-point Hardy Scale in GTEx (GTEx Consortium, 2015). All alterations generated from TCGA and GTEx metadata can be found in the **Table S2**. Furthermore, study name, count-file identifier, and drug information pertaining to TCGA samples were added to *recount-brain-v2*. Additional information about these metadata variables can be found after the “Columns that are not from recount_brain” row of **Table S2**. If there are other metadata variables within TCGA or GTEx that are not part of *recount-brain-v2*, `all_metadata(subset = “gtex”)` or `all_metadata(subset = “tcga”)` can be used to download this metadata and then merged using the `count_file_identifier` variable from *recount-brain-v2*.

### Ontology mapping

To expand *recount-brain* and take advantage of curated ontologies, we matched *recount-brain* to ontologies available via the BioPortal project (Whetzel et al., 2011) such as UBERON (Mungall et al., 2012). For the Brodmann area, we used `brodmann_area` curated variable for matching to UBERON’s preferred label. The diseases were matched to ontologies manually. For tissues, we constructed a hierarchical tissue variable from `tissue_site_3` > `tissue_site_2` > `tissue_site_1` such that the more detailed information is used when available. We then searched UBERON’s preferred labels to identify the ontology entries that best matched the tissue descriptions. For Brodmann area and tissue ontology terms, we extracted their synonyms, parent term IDs and parent term labels to facilitate identifying samples of interest either through text based searches. The code for the ontology mapping is available at http://LieberInstitute.github.io/recount-brain/.

### MetaSRA comparison

We downloaded the MetaSRA (Bernstein et al., 2017) data for UBERON term 0000955 (brain) on April 15, 2019. The table we downloaded included information for 17,890 brain samples from 342 studies. Of these 342 studies, 100 included at least one sample present in *recount2* that are absent from *recount-brain*. Based on our selection criteria of at least 4 samples in *recount2* and 70% or brain samples in a study, 28 studies would match the criteria based on MetaSRA data. From these 28 studies, 5 of them are supported by the latest *phenopredict* predictions (version 0.0.06) and 7 by the SHARQ prototype tissue predictions (Kingsford, 2016). Conversely, 17 (26.6%) out of the 64 studies and 3,841 (58.6%) of the samples in *recount-brain* are absent from MetaSRA (including GTEx and TCGA). The code and full comparison results are available at http://LieberInstitute.github.io/recount-brain/. **Table S3** contains the list of the 100 studies with at least one brain sample according to MetaSRA that are present in *recount2* and absent from *recount-brain*.

### Interactive display

The interactive *recount-brain* display at https://jhubiostatistics.shinyapps.io/recount-brain/ was created using a custom version of the shinycsv project (Collado-Torres et al., 2018) from https://github.com/LieberInstitute/shinycsv using the shiny R package (Chang et al., 2019) available from https://cran.r-project.org/web/packages/shiny/index.html.

### Differential expression by tumor grades with data from SRP027383

We downloaded the gene expression data for SRA study SRP027383 (Bao et al., 2014) from *recount2* (Collado-Torres et al., 2017c) using *recount* v1.5.9, added the *recount-brain-v1* sample metadata and retained the 258 samples that have sex, age, pathology (IDH1 mutation either + or -) and tumor grade progression (II, III and IV) recorded. We then filtered the genes with a mean RPKM < 0.24 as suggested by the `expression_cutoff()` function from *jaffelab* v0.99.18 (Collado-Torres and Jaffe, 2018) to retain 25,649 genes. Next we computed library size adjustments with *edgeR* v3.21.9 (McCarthy et al., 2012; Robinson et al., 2010) and performed the differential gene expression using *limma*-voom 3.35.12 (Law et al., 2014; Ritchie et al., 2015). The model we used was `ordered(clinical_stage_1) + sex + age + pathology` such that we fitted a linear and a quadratic term for the tumor grade progression (stored in `clinical_stage_1`). We visualized the voom-normalized gene expression for the top differentially expressed genes (FDR 1%) after removing the effects of sex, age and pathology using the `cleaningY()` function from *jaffelab* v0.99.18 (Collado-Torres and Jaffe, 2018). Using the `compareCluster()` function from *clusterProfiler* v3.7.0 (Yu et al., 2012) with Ensembl (Zerbino et al., 2018) gene ids, p-value and q-value cutoffs of 0.05 we found enriched biological process, molecular function and cellular component ontologies for the genes with a significant linear association with tumor grade progression, separated by the trend direction. Code and full results are provided in **Supplementary File 1**.

### Effects of post-mortem interval on transcription

Using *recount-brain-v1* we identified 252 samples from 10 SRA studies described in nine publications (Boudreau et al., 2014; Dumitriu et al., 2016; Khrameeva et al., 2014; Labadorf et al., 2015; Li et al., 2014; Magistri et al., 2015; Pardo et al., 2013; Voineagu et al., 2011; Wu et al., 2012) with post-mortem interval (PMI) information that have variable measurements so that we could replicate some of the analyses performed by Ferreira *et al*. (Ferreira et al., 2018). We downloaded the gene expression data from *recount2* (Collado-Torres et al., 2017c) using *recount* v1.5.9 and assessed the changes in log(RPKM + 0.5) expression for genes *RNASE2*, *EGR3*, *HBA1* and *CXCL2* across the same PMI intervals (in hours) that Ferreira et al. used in their Figure 2D (Ferreira et al., 2018) after removing the study and disease status with `cleaningY()` from *jaffelab* v0.99.18 (Collado-Torres and Jaffe, 2018). Using the `coverage_matrix()` function from *recount* v1.5.9 (Collado-Torres et al., 2017b) we computed the percent of mitochondrial transcription as well as the global gene transcription for Gencode v25 (Harrow et al., 2012) (excluding mitochondrial genes) for these 252 samples. We compared the mitochondrial and gene transcription levels to each other and to PMI measured in hours similar to Ferreira’s *et al*. Figure 4B and 4C (Ferreira et al., 2018). Exploratory figures were made using *ggplot2* v2.2.1 (Wickham, 2009). Code and full results are provided in **Supplementary File 2**.

### Variation in gene expression across multiple glioblastoma studies

In order to illustrate how *recount-brain* metadata can be utilized with expression data from more than one study in an expression analysis, we used the two glioblastoma studies with at least 20 samples present in *recount-brain-v1*: SRP027383 (N=270) (Bao et al., 2014) and SRP044668 (N=93) (Gill et al., 2014). We then combined these SRA samples with those from TCGA listed as primary glioblastoma tumors (N=157) (Brennan et al., 2013; Cancer Genome Atlas Research Network, 2008) for a total of 520 samples. We used `expression_cutoff()` from *jaffelab* v0.99.21 (Collado-Torres and Jaffe, 2018) to filter the genes with mean RPKM < 0.21, retaining a total of 26,499 genes. As we are specifically interested in assessing variability across glioblastoma tumors in this particular analysis, control samples were dropped, retaining a total of 320 tumor samples: SRP027383 (N=99), SRP044668 (N=74), TCGA (N=157). We normalized for the data source effect by using a linear regression with an indicator variable differentiating the SRA and the TCGA samples and then regressing out this effect. We then removed the top 6 principal components (PCs) computed on the log2(RPKM+0.5) data to facilitate cross-study comparisons.

We then estimated the variance of each gene for each of the three datasets using `colVars()` from *matrixStats* v0.53.1 (Bengtsson, 2018) and compared the most variable gene ranking using concordance at the top plots. We also processed primary tumor kidney TCGA data (Davis et al., 2014) given its biological dissimilarity with human brain. We used a similar normalization procedure with the combined brain and kidney data (top 4 PCs removed) to produce a background pairwise comparison for the concordance at the top plots. Code and full results are provided in **Supplementary File 3**.

## Data Access

The *recount-brain* data (both version 1 and 2) is available via the `add_metadata()`*recount* (Collado-Torres et al., 2017b) Bioconductor package (version >= 1.5.6; see version >= 1.7.6 for updated examples).

There are four different ways to access the *recount-brain* dataset. Both an R version and a text version (csv) are available from the *recount-brain* GitHub repository at https://github.com/LieberInstitute/recount-brain. *recount-brain* can also be explored interactively from https://jhubiostatistics.shinyapps.io/recount-brain/ and subsets of interest can be downloaded to csv files from that website. However, we mainly recommend using the `add_metadata(source = “recount_brain_v2”)` function from the *recount* R package (Collado-Torres et al., 2017b) either with or without the `rse` argument in `add_metadata()` as shown in the help file for this function in *recount* (version >= 1.7.6) (Collado-Torres et al., 2017b). The code for creating *recount-brain* is available at the **Supplementary Website** http://LieberInstitute.github.io/recount-brain/.

## Author contributions

A.R. - Conceptualization, Data Curation, Methodology, Writing – Original Draft Preparation, Writing – Review & Editing.

S.E.E. - Conceptualization, Formal Analysis, Methodology, Writing – Review & Editing.

D.J.S. - Conceptualization, Formal Analysis, Writing – Original Draft Preparation.

S.D. - Conceptualization, Formal Analysis, Writing – Review & Editing.

M.D.W. - Conceptualization, Supervision, Writing – Review & Editing.

J.T.L. - Conceptualization, Funding Acquisition, Writing – Review & Editing.

A.E.J. - Conceptualization, Funding Acquisition, Writing – Review & Editing.

L.C.-T. - Conceptualization, Formal Analysis, Methodology, Software, Supervision, Writing – Original Draft Preparation, Writing – Review & Editing.

## Acknowledgements

We would like to thank Emily E. Burke (Lieber Institute for Brain Development, Baltimore, Maryland, USA) for designing the *recount-brain* logo. The results published here are in whole or part based upon data generated by the TCGA Research Network: http://cancergenome.nih.gov/. *recount2* is hosted on SciServer, a collaborative research environment for large-scale data-driven science. It is being developed at, and administered by, the Institute for Data Intensive Engineering and Science (IDIES) at Johns Hopkins University. SciServer is funded by the National Science Foundation Award ACI-1261715. For more information about SciServer, visit http://www.sciserver.org/. The interactive display for *recount-brain* at https://jhubiostatistics.shinyapps.io/recount-brain/ is hosted by the Johns Hopkins Bloomberg School of Public Health (JHBSPH) Biostatistics Department shinyapps account.

## Supplementary Material

**Supplementary Website**: http://LieberInstitute.github.io/recount-brain/ contains all the R code as well as the supplementary files that can be used to reproduce the entire *recount-brain* project.

**Table S1**: List of variables present in *recount-brain* and description of each variable saved in a CSV file. This file is also available at https://github.com/LieberInstitute/recount-brain/blob/master/SupplementaryTable1.csv. **Table S2**: Detailed notes on how the GTEx and TCGA variables were processed when creating *recount-brain-v2* in order to merge them with *recount-brain-v1*. This file is also available at https://github.com/LieberInstitute/recount-brain/blob/master/SupplementaryTable2.csv.

**Table S3**: List of studies present in MetaSRA and *recount2* with at least one brain sample that are absent in *recount-brain*. Includes the study abstract, whether the abstract mentions brain, number of brain samples listed, percent of brain samples for the project, and whether the study would pass the selection criteria used for this study. This file is also available at http://LieberInstitute.github.io/recount-brain/metasra_comp/discrepant_studies.csv.

**Supplementary File 1**: Full example analysis using data from SRP027383. This file is also available at https://github.com/LieberInstitute/recount-brain/blob/master/example_SRP027383/example_SRP027383.pdf.

**Supplementary File 2**: Full example analysis on the effects of post-mortem interval on transcription. This file is also available at https://github.com/LieberInstitute/recount-brain/blob/master/example_PMI/example_PMI.pdf.

**Supplementary File 3**: Full example meta-analysis on the expression variability in glioblastoma tumor samples. This file is also available at https://github.com/LieberInstitute/recount-brain/blob/master/example_multistudy/recount_brain_multistudy.pdf.

## References

Bao, Z.-S., Chen, H.-M., Yang, M.-Y., Zhang, C.-B., Yu, K., Ye, W.-L., Hu, B.-Q., Yan, W., Zhang, W., Akers, J., Ramakrishnan, V., Li, J., Carter, B., Liu, Y.-W., Hu, H.-M., Wang, Z., Li, M.-Y., Yao, K., Qiu, X.-G., Kang, C.-S., Jiang, T., 2014. RNA-seq of 272 gliomas revealed a novel, recurrent PTPRZ1-MET fusion transcript in secondary glioblastomas. Genome Res. 24, 1765–1773. doi:10.1101/gr.165126.113

Bengtsson, H., 2018. matrixStats: Functions that Apply to Rows and Columns of Matrices (and toVectors). CRAN.

Bernstein, M.N., Doan, A., Dewey, C.N., 2017. MetaSRA: normalized human sample-specific metadata for the Sequence Read Archive. Bioinformatics 33, 2914–2923. doi:10.1093/bioinformatics/btx334

Boudreau, R.L., Jiang, P., Gilmore, B.L., Spengler, R.M., Tirabassi, R., Nelson, J.A., Ross, C.A., Xing, Y., Davidson, B.L., 2014. Transcriptome-wide discovery of microRNA binding sites in human brain. Neuron 81, 294–305. doi:10.1016/j.neuron.2013.10.062

Brennan, C.W., Verhaak, R.G.W., McKenna, A., Campos, B., Noushmehr, H., Salama, S.R., Zheng, S., Chakravarty, D., Sanborn, J.Z., Berman, S.H., Beroukhim, R., Bernard, B., Wu, C.-J., Genovese, G., Shmulevich, I., Barnholtz-Sloan, J., Zou, L., Vegesna, R., Shukla, S.A., Ciriello, G., TCGA Research Network, 2013. The somatic genomic landscape of glioblastoma. Cell 155, 462–477. doi:10.1016/j.cell.2013.09.034

Cancer Genome Atlas Research Network, 2008. Comprehensive genomic characterization defines human glioblastoma genes and core pathways. Nature 455, 1061–1068. doi:10.1038/nature07385

Cancer Genome Atlas Research Network, Brat, D.J., Verhaak, R.G.W., Aldape, K.D., Yung, W.K.A., Salama, S.R., Cooper, L.A.D., Rheinbay, E., Miller, C.R., Vitucci, M., Morozova, O., Robertson, A.G., Noushmehr, H., Laird, P.W., Cherniack, A.D., Akbani, R., Huse, J.T., Ciriello, G., Poisson, L.M., Barnholtz-Sloan, J.S., et al., 2015. Comprehensive, Integrative Genomic Analysis of Diffuse Lower-Grade Gliomas. N. Engl. J. Med. 372, 2481–2498. doi:10.1056/NEJMoa1402121

Castelvecchi, D., 2018. Google unveils search engine for open data. Nature 561, 161–162. doi:10.1038/d41586-018-06201-x

Chang, W., Cheng, J., Allaire, J.J., Xie, Y., McPherson, J., 2019. shiny: Web Application Framework for R.

Collado-Torres, L., Jaffe, A.E., 2018. jaffelab: Commonly used functions by the Jaffe lab. GitHub, GitHub.

Collado-Torres, L., Nellore, A., Jaffe, A.E., 2017a. recount workflow: Accessing over 70,000 human RNA-seq samples with Bioconductor. [version 1; peer review: 1 approved, 2 approved with reservations]. F1000Res. 6, 1558. doi:10.12688/f1000research.12223.1

Collado-Torres, L., Nellore, A., Jaffe, A.E., Taub, M.A., Kammers, K., Ellis, S.E., Hansen, K.D., Langmead, B., Leek, J.T., 2017b. Explore and download data from the recount project. doi: 10.18129/B9.bioc.recount, Bioconductor.

Collado-Torres, L., Nellore, A., Kammers, K., Ellis, S.E., Taub, M.A., Hansen, K.D., Jaffe, A.E., Langmead, B., Leek, J.T., 2017c. Reproducible RNA-seq analysis using recount2. Nat. Biotechnol. 35, 319–321. doi:10.1038/nbt.3838

Collado-Torres, L., Semick, S., Jaffe, A.E., 2018. shinycsv: Explore a table interactively in a shiny application.

Davis, C.F., Ricketts, C.J., Wang, M., Yang, L., Cherniack, A.D., Shen, H., Buhay, C., Kang, H., Kim, S.C., Fahey, C.C., Hacker, K.E., Bhanot, G., Gordenin, D.A., Chu, A., Gunaratne, P.H., Biehl, M., Seth, S., Kaipparettu, B.A., Bristow, C.A., Donehower, L.A., Creighton, C.J., 2014. The somatic genomic landscape of chromophobe renal cell carcinoma. Cancer Cell 26, 319–330. doi:10.1016/j.ccr.2014.07.014

Denk, F., 2017. Don’t let useful data go to waste. Nature 543, 7. doi:10.1038/543007a

Dumitriu, A., Golji, J., Labadorf, A.T., Gao, B., Beach, T.G., Myers, R.H., Longo, K.A., Latourelle, J.C., 2016. Integrative analyses of proteomics and RNA transcriptomics implicate mitochondrial processes, protein folding pathways and GWAS loci in Parkinson disease. BMC Med. Genomics 9, 5. doi:10.1186/s12920-016-0164-y

Ellis, S.E., Collado-Torres, L., Jaffe, A., Leek, J.T., 2018. Improving the value of public RNA-seq expression data by phenotype prediction. Nucleic Acids Res. 46, e54. doi:10.1093/nar/gky102

Ferreira, P.G., Muñoz-Aguirre, M., Reverter, F., Sá Godinho, C.P., Sousa, A., Amadoz, A., Sodaei, R., Hidalgo, M.R., Pervouchine, D., Carbonell-Caballero, J., Nurtdinov, R., Breschi, A., Amador, R., Oliveira, P., Çubuk, C., Curado, J., Aguet, F., Oliveira, C., Dopazo, J., Sammeth, M., Guigó, R., 2018. The effects of death and post-mortem cold ischemia on human tissue transcriptomes. Nat. Commun. 9, 490. doi:10.1038/s41467-017-02772-x

Fu, J., Kammers, K., Nellore, A., Collado-Torres, L., Leek, J.T., Taub, M.A., 2018. RNA-seq transcript quantification from reduced-representation data in recount2. BioRxiv. doi:10.1101/247346

Gill, B.J., Pisapia, D.J., Malone, H.R., Goldstein, H., Lei, L., Sonabend, A., Yun, J., Samanamud, J., Sims, J.S., Banu, M., Dovas, A., Teich, A.F., Sheth, S.A., McKhann, G.M., Sisti, M.B., Bruce, J.N., Sims, P.A., Canoll, P., 2014. MRI-localized biopsies reveal subtype-specific differences in molecular and cellular composition at the margins of glioblastoma. Proc Natl Acad Sci USA 111, 12550–12555. doi:10.1073/pnas.1405839111

GTEx Consortium, 2015. Human genomics. The Genotype-Tissue Expression (GTEx) pilot analysis: multitissue gene regulation in humans. Science 348, 648–660. doi:10.1126/science.1262110

Harrow, J., Frankish, A., Gonzalez, J.M., Tapanari, E., Diekhans, M., Kokocinski, F., Aken, B.L., Barrell, D., Zadissa, A., Searle, S., Barnes, I., Bignell, A., Boychenko, V., Hunt, T., Kay, M., Mukherjee, G., Rajan, J., Despacio-Reyes, G., Saunders, G., Steward, C., Hubbard, T.J., 2012. GENCODE: the reference human genome annotation for The ENCODE Project. Genome Res. 22, 1760–1774. doi:10.1101/gr.135350.111

Hutter, C., Zenklusen, J.C., 2018. The Cancer Genome Atlas: Creating Lasting Value beyond Its Data. Cell 173, 283–285. doi:10.1016/j.cell.2018.03.042

Jiang, L., Zhou, J., Zhong, D., Zhou, Y., Zhang, W., Wu, W., Zhao, Z., Wang, W., Xu, W., He, L., Ma, Y., Hu, Y., Zhang, W., Li, J., 2017. Overexpression of SMC4 activates TGFβ/Smad signaling and promotes aggressive phenotype in glioma cells. Oncogenesis 6, e301. doi:10.1038/oncsis.2017.8

Khrameeva, E.E., Bozek, K., He, L., Yan, Z., Jiang, X., Wei, Y., Tang, K., Gelfand, M.S., Prufer, K., Kelso, J., Paabo, S., Giavalisco, P., Lachmann, M., Khaitovich, P., 2014. Neanderthal ancestry drives evolution of lipid catabolism in contemporary Europeans. Nat. Commun. 5, 3584. doi:10.1038/ncomms4584

Kingsford, C., 2016. SHARQ PROTOTYPE [WWW Document]. Search public, human, RNA-seq experiments by cell, tissue type, and other features | Indexing files. URL http://www.cs.cmu.edu/~ckingsf/sharq/ (accessed 11.13.17).

Kodama, Y., Shumway, M., Leinonen, R., International Nucleotide Sequence Database Collaboration, 2012. The Sequence Read Archive: explosive growth of sequencing data. Nucleic Acids Res. 40, D54–6. doi:10.1093/nar/gkr854

Labadorf, A., Hoss, A.G., Lagomarsino, V., Latourelle, J.C., Hadzi, T.C., Bregu, J., MacDonald, M.E., Gusella, J.F., Chen, J.-F., Akbarian, S., Weng, Z., Myers, R.H., 2015. RNA sequence analysis of human huntington disease brain reveals an extensive increase in inflammatory and developmental gene expression. PLoS ONE 10, e0143563. doi:10.1371/journal.pone.0143563

Langmead, B., Nellore, A., 2018. Cloud computing for genomic data analysis and collaboration. Nat. Rev. Genet. 19, 208–219. doi:10.1038/nrg.2017.113

Law, C.W., Chen, Y., Shi, W., Smyth, G.K., 2014. voom: Precision weights unlock linear model analysis tools for RNA-seq read counts. Genome Biol. 15, R29. doi:10.1186/gb-2014-15-2-r29

Leinonen, R., Sugawara, H., Shumway, M., International Nucleotide Sequence Database Collaboration, 2011. The sequence read archive. Nucleic Acids Res. 39, D19–21. doi:10.1093/nar/gkq1019

Li, J., Shi, M., Ma, Z., Zhao, S., Euskirchen, G., Ziskin, J., Urban, A., Hallmayer, J., Snyder, M., 2014. Integrated systems analysis reveals a molecular network underlying autism spectrum disorders. Mol. Syst. Biol. 10, 774. doi:10.15252/msb.20145487

Magistri, M., Velmeshev, D., Makhmutova, M., Faghihi, M.A., 2015. Transcriptomics Profiling of Alzheimer’s Disease Reveal Neurovascular Defects, Altered Amyloid-β Homeostasis, and Deregulated Expression of Long Noncoding RNAs. J Alzheimers Dis 48, 647–665. doi:10.3233/JAD-150398

McCarthy, D.J., Chen, Y., Smyth, G.K., 2012. Differential expression analysis of multifactor RNA-Seq experiments with respect to biological variation. Nucleic Acids Res. 40, 4288–4297. doi:10.1093/nar/gks042

Mungall, C.J., Torniai, C., Gkoutos, G.V., Lewis, S.E., Haendel, M.A., 2012. Uberon, an integrative multi-species anatomy ontology. Genome Biol. 13, R5. doi:10.1186/gb-2012-13-1-r5

NCBI Resource Coordinators, 2018. Database resources of the National Center for Biotechnology Information. Nucleic Acids Res. 46, D8–D13. doi:10.1093/nar/gkx1095

Pardo, L.M., Rizzu, P., Francescatto, M., Vitezic, M., Leday, G.G.R., Sanchez, J.S., Khamis, A., Takahashi, H., van de Berg, W.D.J., Medvedeva, Y.A., van de Wiel, M.A., Daub, C.O., Carninci, P., Heutink, P., 2013. Regional differences in gene expression and promoter usage in aged human brains. Neurobiol. Aging 34, 1825–1836. doi:10.1016/j.neurobiolaging.2013.01.005

Ritchie, M.E., Phipson, B., Wu, D., Hu, Y., Law, C.W., Shi, W., Smyth, G.K., 2015. limma powers differential expression analyses for RNA-sequencing and microarray studies. Nucleic Acids Res. 43, e47. doi:10.1093/nar/gkv007

Robinson, M.D., McCarthy, D.J., Smyth, G.K., 2010. edgeR: a Bioconductor package for differential expression analysis of digital gene expression data. Bioinformatics 26, 139–140. doi:10.1093/bioinformatics/btp616

SEQC/MAQC-III Consortium, 2014. A comprehensive assessment of RNA-seq accuracy, reproducibility and information content by the Sequencing Quality Control Consortium. Nat. Biotechnol. 32, 903–914. doi:10.1038/nbt.2957

Vera Alvarez, R., Medeiros Vidal, N., Garzón-Martínez, G.A., Barrero, L.S., Landsman, D., Mariño-Ramírez, L., 2017. Workflow and web application for annotating NCBI BioProject transcriptome data. Database (Oxford) 2017. doi:10.1093/database/bax008

Voineagu, I., Wang, X., Johnston, P., Lowe, J.K., Tian, Y., Horvath, S., Mill, J., Cantor, R.M., Blencowe, B.J., Geschwind, D.H., 2011. Transcriptomic analysis of autistic brain reveals convergent molecular pathology. Nature 474, 380–384. doi:10.1038/nature10110

Whetzel, P.L., Noy, N.F., Shah, N.H., Alexander, P.R., Nyulas, C., Tudorache, T., Musen, M.A., 2011. BioPortal: enhanced functionality via new Web services from the National Center for Biomedical Ontology to access and use ontologies in software applications. Nucleic Acids Res. 39, W541–5. doi:10.1093/nar/gkr469

Wickham, H., 2009. ggplot2: Elegant graphics for data analysis. Springer, New York.

Wilks, C., Gaddipati, P., Nellore, A., Langmead, B., 2018. Snaptron: querying splicing patterns across tens of thousands of RNA-seq samples. Bioinformatics 34, 114–116. doi:10.1093/bioinformatics/btx547

Wu, J.Q., Wang, X., Beveridge, N.J., Tooney, P.A., Scott, R.J., Carr, V.J., Cairns, M.J., 2012. Transcriptome sequencing revealed significant alteration of cortical promoter usage and splicing in schizophrenia. PLoS ONE 7, e36351. doi:10.1371/journal.pone.0036351

Yu, G., Wang, L.-G., Han, Y., He, Q.-Y., 2012. clusterProfiler: an R package for comparing biological themes among gene clusters. OMICS 16, 284–287. doi:10.1089/omi.2011.0118

Zerbino, D.R., Achuthan, P., Akanni, W., Amode, M.R., Barrell, D., Bhai, J., Billis, K., Cummins, C., Gall, A., Girón, C.G., Gil, L., Gordon, L., Haggerty, L., Haskell, E., Hourlier, T., Izuogu, O.G., Janacek, S.H., Juettemann, T., To, J.K., Laird, M.R., Flicek, P., 2018. Ensembl 2018. Nucleic Acids Res. 46, D754–D761. doi:10.1093/nar/gkx1098

