## Supplementary File 1 for "*recount-brain*: a curated repository of human brain RNA-seq datasets metadata"

### recount_brain example with data from SRP027383

**3 March 2018**

**Abstract**

This is an example on how to use recount_brain applied to the SRP027383 study. We show how to download data from recount2, add the sample metadata from recount_brain, explore the sample metadata and the gene expression data, and perform a gene expression analysis.

### 1 Introduction

This document is an example of how you can use `recount_brain`. We will use the data from the SRA study SRP027383 which is described in "RNA-seq of 272 gliomas revealed a novel, recurrent PTPRZ1-MET fusion transcript in secondary glioblastomas" (Bao, Chen, Yang, Zhang, et al., 2014). As you can see in Figure 1 a lot of the metadata for these samples is missing from the SRA Run Selector which makes it a great case for using `recount_brain`. We will show how to add the `recount_brain` metadata and perform a gene differential expression analysis using this information.

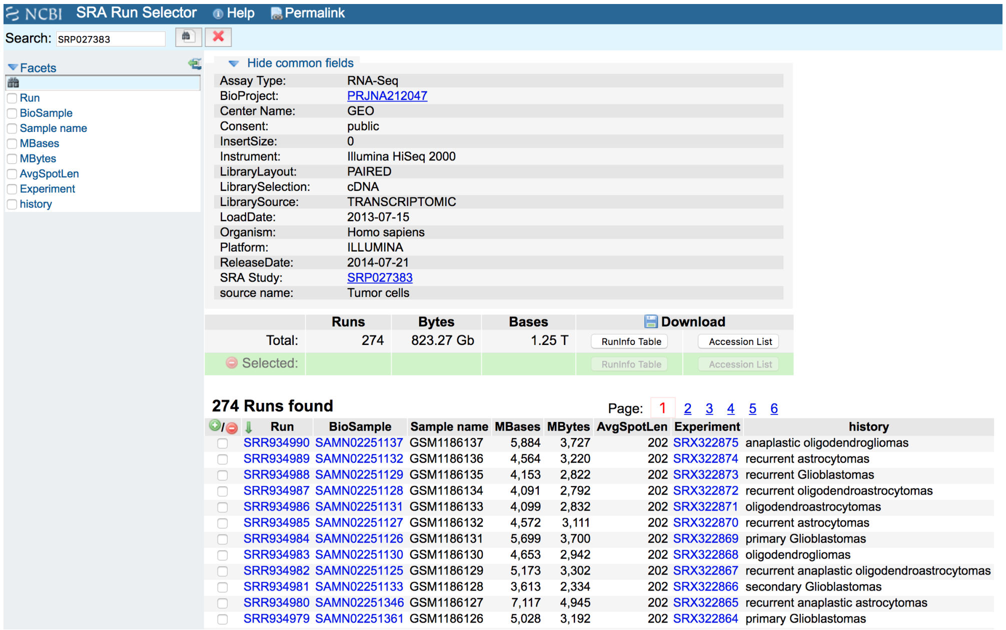

**Figure 1: SRA Run Selector information for study SRP027383**
Screenshot from 2018-02-26.

### 2 Sample metadata

Just like any study in `recount2` (Collado-Torres, Nellore, Kammers, Ellis, et al., 2017), we first need to download the gene count data using `recount::download_study()`. Since we will be using many functions from the `recount` package, lets load it first[1].

```
## Load the package
library('recount')
```

[1] If you are a first time `recount` user, we recommend first reading the package vignette at bioconductor.org/packages/recount.

#### 2.1 Download gene data

Having loaded the package, we next download the gene-level data.

```
if(!file.exists(file.path('SRP027383', 'rse_gene.Rdata'))) {
    download_study('SRP027383')
}
load(file.path('SRP027383', 'rse_gene.Rdata'), verbose = TRUE)
## Loading objects:
##   rse_gene
```

#### 2.2 Sample metadata included in `recount`

We can next explore the sample metadata that is included by default using `SummarizedExperiment::colData()`. These variables are explained in more detail in the supplementary material of the `recount2` paper (Collado-Torres, Nellore, Kammers, Ellis, et al., 2017) and in the `recount workflow` paper (Collado-Torres, Nellore, and Jaffe, 2017).

```
colData(rse_gene)
## DataFrame with 270 rows and 21 columns
##                 project      sample  experiment         run
##             <character> <character> <character> <character>
## SRR934717   SRP027383   SRS457680   SRX322602   SRR934717
## SRR934718   SRP027383   SRS457681   SRX322603   SRR934718
## SRR934719   SRP027383   SRS457682   SRX322604   SRR934719
## SRR934720   SRP027383   SRS457683   SRX322605   SRR934720
## SRR934721   SRP027383   SRS457684   SRX322606   SRR934721
## ...               ...         ...         ...         ...
## SRR934986   SRP027383   SRS457949   SRX322871   SRR934986
## SRR934987   SRP027383   SRS457950   SRX322872   SRR934987
## SRR934988   SRP027383   SRS457951   SRX322873   SRR934988
## SRR934989   SRP027383   SRS457952   SRX322874   SRR934989
## SRR934990   SRP027383   SRS457953   SRX322875   SRR934990
##             read_count_as_reported_by_sra reads_downloaded
##                                 <integer>        <integer>
## SRR934717                        56887576         56887576
## SRR934718                        39683692         39683692
## SRR934719                        39392540         39392540
## SRR934720                        60287388         60287388
## SRR934721                        31089346         31089346
## ...                                   ...              ...
## SRR934986                        42563170         42563170
## SRR934987                        42481802         42481802
## SRR934988                        43121132         43121132
## SRR934989                        47384314         47384314
## SRR934990                        61093682         61093682
##             proportion_of_reads_reported_by_sra_downloaded paired_end
##                                                  <numeric>  <logical>
## SRR934717                                                1       TRUE
## SRR934718                                                1       TRUE
## SRR934719                                                1       TRUE
## SRR934720                                                1       TRUE
## SRR934721                                                1       TRUE
```

```
## ...                                                  ...        ...
## SRR934986                                              1       TRUE
## SRR934987                                              1       TRUE
## SRR934988                                              1       TRUE
## SRR934989                                              1       TRUE
## SRR934990                                              1       TRUE
##           sra_misreported_paired_end mapped_read_count        auc
##                            <logical>         <integer>  <numeric>
## SRR934717                      FALSE          56189295 5628071616
## SRR934718                      FALSE          39636163 3950872208
## SRR934719                      FALSE          39373323 3958083805
## SRR934720                      FALSE          60261401 6047049537
## SRR934721                      FALSE          30964054 3072882301
## ...                              ...               ...        ...
## SRR934986                      FALSE          42449491 4259218453
## SRR934987                      FALSE          42358446 4245759225
## SRR934988                      FALSE          42997366 4309934199
## SRR934989                      FALSE          47223491 4739386115
## SRR934990                      FALSE          60917502 6110940825
##           sharq_beta_tissue sharq_beta_cell_type biosample_submission_date
##                 <character>          <character>               <character>
## SRR934717    umbilical cord                  esc   2013-07-15T11:26:36.860
## SRR934718    umbilical cord                  esc   2013-07-15T11:28:33.710
## SRR934719    umbilical cord                  esc   2013-07-15T11:26:47.540
## SRR934720    umbilical cord                  esc   2013-07-15T11:26:44.253
## SRR934721    umbilical cord                  esc   2013-07-15T11:28:18.330
## ...                     ...                  ...                       ...
## SRR934986    umbilical cord                  esc   2013-07-15T11:22:27.600
## SRR934987    umbilical cord                  esc   2013-07-15T11:22:07.083
## SRR934988    umbilical cord                  esc   2013-07-15T11:22:10.270
## SRR934989    umbilical cord                  esc   2013-07-15T11:22:37.680
## SRR934990    umbilical cord                  esc   2013-07-15T11:23:19.253
##           biosample_publication_date   biosample_update_date avg_read_length
##                          <character>             <character>       <integer>
## SRR934717    2014-07-20T00:44:13.497 2014-07-20T01:22:14.790             202
## SRR934718    2014-07-20T00:44:16.773 2014-07-20T01:22:14.977             200
## SRR934719    2014-07-20T00:44:13.637 2014-07-20T01:22:15.377             202
## SRR934720    2014-07-20T00:44:13.573 2014-07-20T01:22:15.650             202
## SRR934721    2014-07-20T00:44:16.493 2014-07-20T01:22:16.003             200
## ...                              ...                     ...             ...
## SRR934986    2014-07-20T00:44:09.693 2014-07-20T01:15:29.503             202
## SRR934987    2014-07-20T00:44:09.567 2014-07-20T01:18:22.877             202
## SRR934988    2014-07-20T00:44:09.610 2014-07-20T01:18:23.733             202
## SRR934989    2014-07-20T00:44:09.730 2014-07-20T01:18:24.270             202
## SRR934990    2014-07-20T00:44:09.930 2014-07-20T01:18:25.100             202
##           geo_accession  bigwig_file       title
##             <character>  <character> <character>
## SRR934717    GSM1185864 SRR934717.bw    CGGA_171
## SRR934718    GSM1185865 SRR934718.bw    CGGA_235
## SRR934719    GSM1185866 SRR934719.bw    CGGA_236
## SRR934720    GSM1185867 SRR934720.bw    CGGA_241
```

```
## SRR934721    GSM1185868 SRR934721.bw    CGGA_243
## ...                 ...          ...         ...
## SRR934986    GSM1186133 SRR934986.bw   CGGA_J030
## SRR934987    GSM1186134 SRR934987.bw   CGGA_J042
## SRR934988    GSM1186135 SRR934988.bw   CGGA_J100
## SRR934989    GSM1186136 SRR934989.bw   CGGA_J130
## SRR934990    GSM1186137 SRR934990.bw   CGGA_J023
##                                    characteristics
##                                    <CharacterList>
## SRR934717          history: oligodendroastrocytomas
## SRR934718          history: oligodendroastrocytomas
## SRR934719               history: oligodendrogliomas
## SRR934720          history: oligodendroastrocytomas
## SRR934721          history: oligodendroastrocytomas
## ...                                             ...
## SRR934986          history: oligodendroastrocytomas
## SRR934987 history: recurrent oligodendroastrocytomas
## SRR934988          history: recurrent Glioblastomas
## SRR934989           history: recurrent astrocytomas
## SRR934990     history: anaplastic oligodendrogliomas
```

Note how the `characteristics` column matches the information from the SRA Run Selector in Figure 1. Still not very useful.

```
colData(rse_gene)$characteristics
## CharacterList of length 270
## [[1]] history: oligodendroastrocytomas
## [[2]] history: oligodendroastrocytomas
## [[3]] history: oligodendrogliomas
## [[4]] history: oligodendroastrocytomas
## [[5]] history: oligodendroastrocytomas
## [[6]] history: recurrent astrocytomas
## [[7]] history: oligodendroastrocytomas
## [[8]] history: astrocytomas
## [[9]] history: oligodendroastrocytomas
## [[10]] history: astrocytomas
## ...
## <260 more elements>
```

#### 2.3 Add `recount_brain` sample metadata

So lets add the available sample metadata from `recount_brain` using the `recount::add_metadata()` function.

```
rse_gene <- add_metadata(rse = rse_gene, source = 'recount_brain_v1')
## 2018-03-03 11:21:16 downloading the recount_brain metadata to /var/folders/cx/n9s558kx6fb7jf5z_pgszgb8000
## Loading objects:
##   recount_brain
## 2018-03-03 11:21:17 found 270 out of 270 samples in the recount_brain metadata
```

#### 2.4 Explore `recount_brain` metadata

We can now explore the available metadata from `recount_brain` for the SRP027383 study.

```
## Find which new columns have observations
new_non_NA <- sapply(22:ncol(colData(rse_gene)),
    function(i) any(!is.na(colData(rse_gene)[, i])) )
## Display the observations
colData(rse_gene)[, (22:ncol(colData(rse_gene)))[new_non_NA]]
## DataFrame with 270 rows and 33 columns
##           assay_type_s avgspotlen_l bioproject_s  biosample_s center_name_s
##            <character>    <integer>  <character>  <character>   <character>
## SRR934717      RNA-Seq          202  PRJNA212047 SAMN02251223           GEO
## SRR934718      RNA-Seq          200  PRJNA212047 SAMN02251267           GEO
## SRR934719      RNA-Seq          202  PRJNA212047 SAMN02251226           GEO
## SRR934720      RNA-Seq          202  PRJNA212047 SAMN02251225           GEO
## SRR934721      RNA-Seq          200  PRJNA212047 SAMN02251260           GEO
## ...                ...          ...          ...          ...           ...
## SRR934986      RNA-Seq          202  PRJNA212047 SAMN02251131           GEO
## SRR934987      RNA-Seq          202  PRJNA212047 SAMN02251128           GEO
## SRR934988      RNA-Seq          202  PRJNA212047 SAMN02251129           GEO
## SRR934989      RNA-Seq          202  PRJNA212047 SAMN02251132           GEO
## SRR934990      RNA-Seq          202  PRJNA212047 SAMN02251137           GEO
##             consent_s disease_status experiment_s insertsize_l
##           <character>    <character>  <character>    <integer>
## SRR934717      public        Disease    SRX322602            0
## SRR934718      public        Disease    SRX322603            0
## SRR934719      public        Disease    SRX322604            0
## SRR934720      public        Disease    SRX322605            0
## SRR934721      public        Disease    SRX322606            0
## ...               ...            ...          ...          ...
## SRR934986      public        Disease    SRX322871            0
## SRR934987      public        Disease    SRX322872            0
## SRR934988      public        Disease    SRX322873            0
## SRR934989      public        Disease    SRX322874            0
## SRR934990      public        Disease    SRX322875            0
##                  instrument_s librarylayout_s libraryselection_s
##                   <character>     <character>        <character>
## SRR934717 Illumina HiSeq 2000          PAIRED               cDNA
## SRR934718 Illumina HiSeq 2000          PAIRED               cDNA
## SRR934719 Illumina HiSeq 2000          PAIRED               cDNA
## SRR934720 Illumina HiSeq 2000          PAIRED               cDNA
## SRR934721 Illumina HiSeq 2000          PAIRED               cDNA
## ...                       ...             ...                ...
## SRR934986 Illumina HiSeq 2000          PAIRED               cDNA
## SRR934987 Illumina HiSeq 2000          PAIRED               cDNA
## SRR934988 Illumina HiSeq 2000          PAIRED               cDNA
## SRR934989 Illumina HiSeq 2000          PAIRED               cDNA
## SRR934990 Illumina HiSeq 2000          PAIRED               cDNA
##           librarysource_s  loaddate_s  mbases_l  mbytes_l   organism_s
##               <character> <character> <integer> <integer>  <character>
```

```
## SRR934717  TRANSCRIPTOMIC  2013-07-15      5479      3584 Homo sapiens
## SRR934718  TRANSCRIPTOMIC  2013-07-15      3784      2853 Homo sapiens
## SRR934719  TRANSCRIPTOMIC  2013-07-15      3794      2650 Homo sapiens
## SRR934720  TRANSCRIPTOMIC  2013-07-15      5806      3829 Homo sapiens
## SRR934721  TRANSCRIPTOMIC  2013-07-15      2964      2267 Homo sapiens
## ...                   ...         ...       ...       ...          ...
## SRR934986  TRANSCRIPTOMIC  2013-07-15      4099      2832 Homo sapiens
## SRR934987  TRANSCRIPTOMIC  2013-07-15      4091      2792 Homo sapiens
## SRR934988  TRANSCRIPTOMIC  2013-07-15      4153      2822 Homo sapiens
## SRR934989  TRANSCRIPTOMIC  2013-07-15      4564      3220 Homo sapiens
## SRR934990  TRANSCRIPTOMIC  2013-07-15      5884      3727 Homo sapiens
##            platform_s releasedate_s sample_name_s sra_sample_s sra_study_s
##           <character>   <character>   <character>  <character> <character>
## SRR934717    ILLUMINA    2014-07-21    GSM1185864    SRS457680   SRP027383
## SRR934718    ILLUMINA    2014-07-21    GSM1185865    SRS457681   SRP027383
## SRR934719    ILLUMINA    2014-07-21    GSM1185866    SRS457682   SRP027383
## SRR934720    ILLUMINA    2014-07-21    GSM1185867    SRS457683   SRP027383
## SRR934721    ILLUMINA    2014-07-21    GSM1185868    SRS457684   SRP027383
## ...               ...           ...           ...          ...         ...
## SRR934986    ILLUMINA    2014-07-21    GSM1186133    SRS457949   SRP027383
## SRR934987    ILLUMINA    2014-07-21    GSM1186134    SRS457950   SRP027383
## SRR934988    ILLUMINA    2014-07-21    GSM1186135    SRS457951   SRP027383
## SRR934989    ILLUMINA    2014-07-21    GSM1186136    SRS457952   SRP027383
## SRR934990    ILLUMINA    2014-07-21    GSM1186137    SRS457953   SRP027383
##           sample_origin development         sex   age_units       age
##             <character> <character> <character> <character> <numeric>
## SRR934717         Brain       Adult      female       Years        37
## SRR934718         Brain       Adult        male       Years        25
## SRR934719         Brain       Adult        male       Years        47
## SRR934720         Brain       Adult        male       Years        34
## SRR934721         Brain       Adult      female       Years        31
## ...                 ...         ...         ...         ...       ...
## SRR934986         Brain       Adult        male       Years        38
## SRR934987         Brain       Adult        male       Years        38
## SRR934988         Brain       Adult        male       Years        55
## SRR934989         Brain       Adult        male       Years        40
## SRR934990         Brain       Adult        male       Years        36
##               disease clinical_stage_1                    tumor_type
##           <character>      <character>                   <character>
## SRR934717       Tumor         Grade II       Oligodendroastrocytoma
## SRR934718       Tumor         Grade II       Oligodendroastrocytoma
## SRR934719       Tumor         Grade II            Oligodendroglioma
## SRR934720       Tumor         Grade II       Oligodendroastrocytoma
## SRR934721       Tumor         Grade II       Oligodendroastrocytoma
## ...               ...              ...                          ...
## SRR934986       Tumor         Grade II       Oligodendroastrocytoma
## SRR934987       Tumor         Grade II       Oligodendroastrocytoma
## SRR934988       Tumor         Grade IV                  Glioblastoma
## SRR934989       Tumor         Grade II                   Astrocytoma
## SRR934990       Tumor        Grade III Anaplastic Oligodendrogliomas
##                 pathology clinical_stage_2 present_in_recount
```

```
##                <character>      <character>          <logical>
## SRR934717 + IDH1 Mutation               NA               TRUE
## SRR934718 - IDH1 Mutation               NA               TRUE
## SRR934719 + IDH1 Mutation               NA               TRUE
## SRR934720 + IDH1 Mutation               NA               TRUE
## SRR934721                NA             NA               TRUE
## ...                     ...            ...                ...
## SRR934986 - IDH1 Mutation               NA               TRUE
## SRR934987 + IDH1 Mutation        Recurrent               TRUE
## SRR934988 + IDH1 Mutation        Recurrent               TRUE
## SRR934989 - IDH1 Mutation        Recurrent               TRUE
## SRR934990 + IDH1 Mutation               NA               TRUE
```

Several of these variables are technical and may be duplicated with data already present, such as the SRA Experiment ids. We can still use them to verify that entries are correctly matched. Other variables might not be of huge relevance for this study such as `disease_status` since all samples in this study are from diseased tissue. However, they might be useful when working with other studies or doing meta-analyses.

```
## Check experiment ids
identical(rse_gene$experiment, rse_gene$experiment_s)
## [1] TRUE

## No healthy controls in this study
table(rse_gene$disease_status)
##
## Disease
##     270

## All ages reported in the same unit
table(rse_gene$age_units)
##
## Years
##   270
```

In this study there are several variables of biological interest that we can use for different analyses. We have information about `sex`, `age`, `tumor_type`, `pathology`, `clinical_stage_1` and `clinical_stage_2`. These variables are described in more detail in the original study (Bao, Chen, Yang, Zhang, et al., 2014). Below we explore each variable at a time, to get an idea on how diverse the data is.

```
## Univariate exploration of the biological variables for SRP027383
table(rse_gene$sex)
##
## female   male
##    102    166
summary(rse_gene$age)
##    Min. 1st Qu.  Median    Mean 3rd Qu.    Max.    NA's
##   18.00   36.00   42.00   43.12   51.00   81.00       2
table(rse_gene$clinical_stage_1)
##
##  Grade II Grade III  Grade IV
```

```
##        98        72        98
table(rse_gene$tumor_type)
##
##            Anaplastic Astrocytomas Anaplastic Oligodendroastrocytomas
##                                 24                                 35
##      Anaplastic Oligodendrogliomas                        Astrocytoma
##                                 13                                 41
##                        Glioblastoma              Oligodendroastrocytoma
##                                 99                                 37
##                   Oligodendroglioma
##                                 21
table(rse_gene$pathology, useNA = 'ifany')
##
## - IDH1 Mutation + IDH1 Mutation            <NA>
##             121             137              12
table(rse_gene$clinical_stage_2, useNA = 'ifany')
##
##   Primary Recurrent Secondary      <NA>
##        59        59        20       132
```

We can ask some questions such as is there a difference in the mean age by sex or if the tumor grade (`clinical_stage_1`), the tumor type or the pathology is associated with sex. The answer is no for these questions so we can infer that the study design is well balanced so far.

```
## Age mean difference by sex? No
with(colData(rse_gene), t.test(age ~ sex))
##
##  Welch Two Sample t-test
##
## data:  age by sex
## t = 0.52713, df = 201.03, p-value = 0.5987
## alternative hypothesis: true difference in means is not equal to 0
## 95 percent confidence interval:
##  -2.101339  3.634767
## sample estimates:
## mean in group female   mean in group male
##             43.59804             42.83133

## Tumor grade and sex association? No
with(colData(rse_gene), addmargins(table(sex, clinical_stage_1)))
##         clinical_stage_1
## sex      Grade II Grade III Grade IV Sum
##   female       41        27       34 102
##   male         57        45       64 166
##   Sum          98        72       98 268
with(colData(rse_gene), chisq.test(table(sex, clinical_stage_1)))
##
##  Pearson's Chi-squared test
##
## data:  table(sex, clinical_stage_1)
```

```
## X-squared = 1.0736, df = 2, p-value = 0.5846

## Tumor type and sex association? No
with(colData(rse_gene), addmargins(table(sex, tumor_type)))
##         tumor_type
## sex      Anaplastic Astrocytomas Anaplastic Oligodendroastrocytomas
##   female                       7                                 18
##   male                        17                                 17
##   Sum                         24                                 35
##         tumor_type
## sex      Anaplastic Oligodendrogliomas Astrocytoma Glioblastoma
##   female                             2          18           34
##   male                              11          23           64
##   Sum                               13          41           98
##         tumor_type
## sex      Oligodendroastrocytoma Oligodendroglioma Sum
##   female                     16                 7 102
##   male                       20                14 166
##   Sum                        36                21 268
with(colData(rse_gene), chisq.test(table(sex, tumor_type)))
## Warning in chisq.test(table(sex, tumor_type)): Chi-squared approximation may
## be incorrect
##
##  Pearson's Chi-squared test
##
## data:  table(sex, tumor_type)
## X-squared = 8.1801, df = 6, p-value = 0.2252

## Sex and pathology association? No
with(colData(rse_gene), addmargins(table(sex, pathology)))
##         pathology
## sex      - IDH1 Mutation + IDH1 Mutation Sum
##   female              39              59  98
##   male                82              78 160
##   Sum                121             137 258
with(colData(rse_gene), chisq.test(table(sex, pathology)))
##
##  Pearson's Chi-squared test with Yates' continuity correction
##
## data:  table(sex, pathology)
## X-squared = 2.7583, df = 1, p-value = 0.09675
```

### 3 Gene differential expression analysis

#### 3.1 Gene DE setup

Now that we have sample metadata to work with we can proceed to perform a differential expression analysis at the gene level. To get started we need to load some packages.

```
## Load required packages for DE analysis
library('limma')
library('edgeR')
library('jaffelab')
## You can install it with
# devtools::install_github('LieberInstitute/jaffelab')
```

From our earlier exploration, we noticed that not all samples have pathology information, so we will drop those that are missing this information.

```
## Keep only the samples that have pathology reported
has_patho <- rse_gene[, !is.na(rse_gene$pathology)]
```

Next we will compute RPKM values and use `expression_cutoff()` from *jaffelab* to get a suggested RPKM cutoff for dropping genes with low expression levels. Note that you can also use *genefilter* or other packages for computing a low expression cutoff. Figure 2 shows the relationship between the mean RPKM cutoff and the number of features above the given cutoff. Figure 3 is the same information but in percent. Figure 4 is a tad more complicated as it explore the relationship between the cutoff and the distribution of the number of non-zero samples. All three figures show estimated points where the curves bend and simply provide a guide for choosing a cutoff.

```
## Compute RPKM and mean RPKM
rpkm <- getRPKM(scale_counts(has_patho))
rpkm_mean <- rowMeans(rpkm)
## Esmate a mean RPKM cutoff
expr_cuts <- expression_cutoff(rpkm)
```

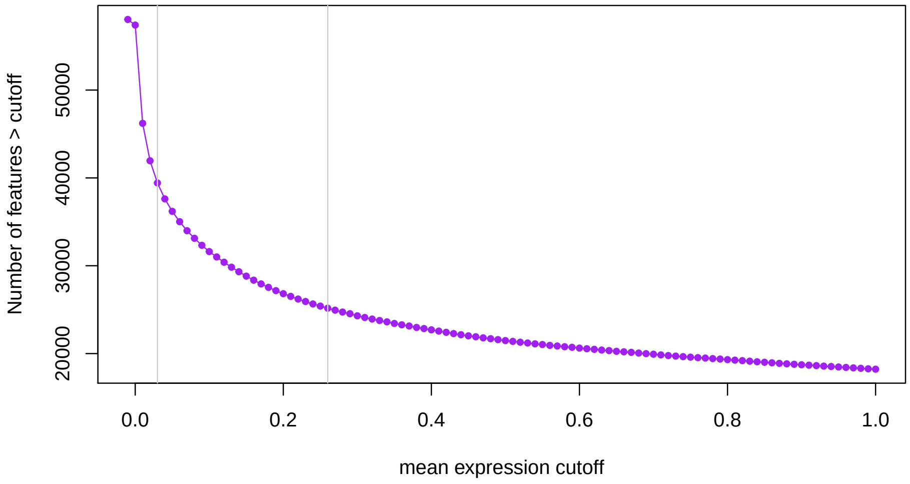

**Figure 2: Number of genes expressed at given mean RPKM cutoff**

```
## 2018-03-03 11:22:19 the suggested expression cutoff is 0.24
```

```
round(mean(expr_cuts), 2)
## [1] 0.24
```

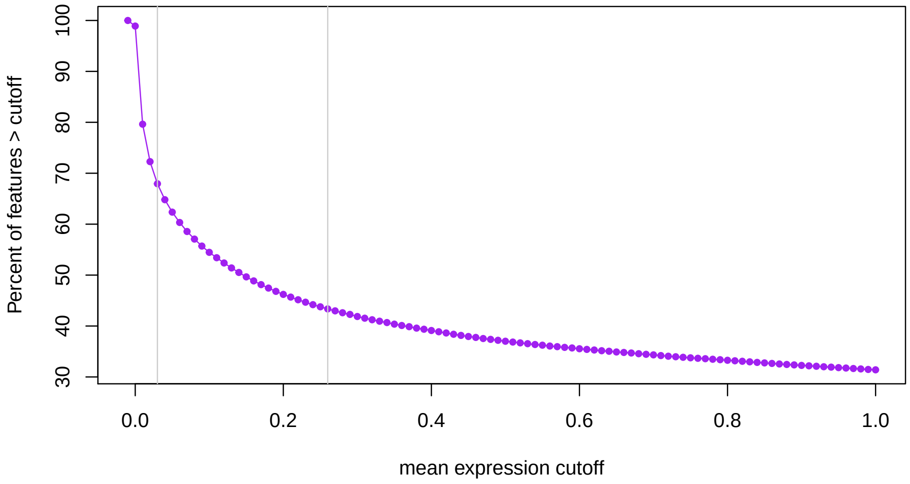

**Figure 3:** **Percent of genes epxressed at a given mean RPKM cutoff**

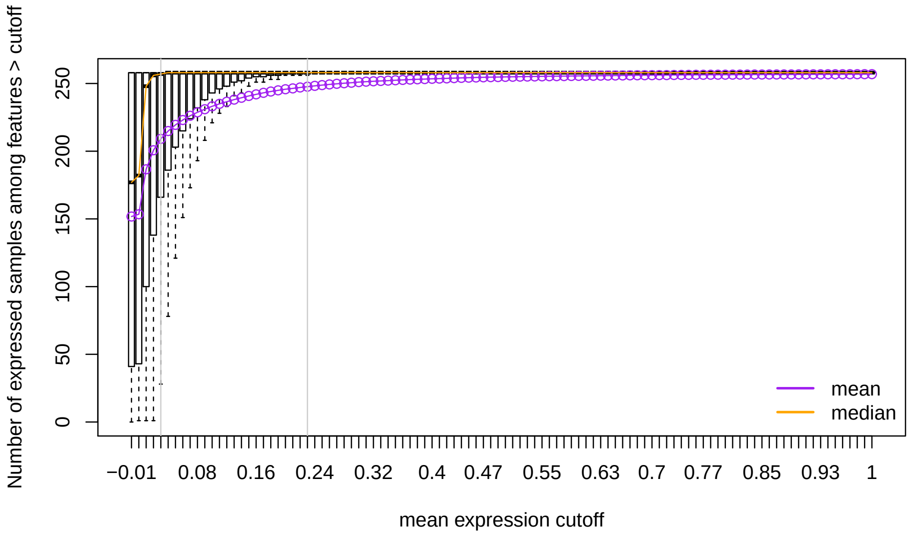

**Figure 4:** **Distribution of number of expressed samples across all genes at a given mean RPKM cutoff**

```
## Filter genes with low levels of expression
has_patho <- has_patho[rpkm_mean > round(mean(expr_cuts), 2), ]
```

Having filtered the genes with low levels of expression, we can now normalize the read counts and identify genes that either have a linear trend or quadratic trend in expression levels between tumor grades II, III and IV while adjusting for age, sex and pathology. Note that this is just an example and you are welcome to try other models. We will use functions from *edgeR* and *limma*.

```
## Get read counts and normalize
dge <- DGEList(counts = assays(scale_counts(has_patho))$counts,
    genes = rowRanges(has_patho))
## Warning in as.data.frame(mcols(x), ...): Arguments in '...' ignored
```

```
dge <- calcNormFactors(dge)

## Build the DE model
## See https://support.bioconductor.org/p/54707/ for details
mod <- with(colData(has_patho),
    model.matrix(~ ordered(clinical_stage_1) + sex + age + pathology))

## Terms of the DE model
colnames(mod)
## [1] "(Intercept)"                  "ordered(clinical_stage_1).L"
## [3] "ordered(clinical_stage_1).Q" "sexmale"
## [5] "age"                          "pathology+ IDH1 Mutation"

## Check that the dimensions match
stopifnot(ncol(dge) == nrow(mod))

## Run voom then run limma model
gene_voom <- voom(dge, mod)
gene_fit <- eBayes(lmFit(gene_voom, mod))
```

Now that we have fitted our differential expression model we can find which genes have a linear or a quadratic change in expression along tumor grade progression. At a false discovery rate (FDR) of 1% none of the genes have a quadratic effect.

```
## Extract the stats for both coefficients
stats_linear <- topTable(gene_fit, coef = 2, p.value = 1,
    number = nrow(has_patho), sort.by = 'none')
stats_quad <- topTable(gene_fit, coef = 3, p.value = 1,
    number = nrow(has_patho), sort.by = 'none')

## How many genes are DE for the linear and the quadratic terms at FDR 1%?
addmargins(table('FDR 1% DE linear' = stats_linear$adj.P.Val < 0.01,
    'FDR 1% DE quadractic' = stats_quad$adj.P.Val < 0.01))
##                  FDR 1% DE quadractic
## FDR 1% DE linear FALSE   Sum
##            FALSE 13095 13095
##            TRUE  12554 12554
##            Sum   25649 25649
```

The fold changes are not necessarily going in the same directions for the differentially expressed genes in the linear term. From the Chi-squared test we can see that the signs are not independent. We could use this information to further explore the gene subsets.

```
## Are the fold changes on the same direction?
addmargins(table(
    'logFC sign linear' = sign(stats_linear$logFC[
        stats_linear$adj.P.Val < 0.01]),
    'logFC sign quadratic' = sign(stats_quad$logFC[
        stats_linear$adj.P.Val < 0.01]))
)
##                    logFC sign quadratic
## logFC sign linear    -1     1   Sum
```

```
##               -1   2626  3490  6116
##               1    4066  2372  6438
##               Sum  6692  5862 12554
chisq.test(table(
    'logFC sign linear' = sign(stats_linear$logFC[
        stats_linear$adj.P.Val < 0.01]),
    'logFC sign quadratic' = sign(stats_quad$logFC[
        stats_linear$adj.P.Val < 0.01]))
)
##
##  Pearson's Chi-squared test with Yates' continuity correction
##
## data:  table(`logFC sign linear` = sign(stats_linear$logFC[stats_linear$adj.P.Val <     0.01]), `logFC sig
## X-squared = 514.36, df = 1, p-value < 2.2e-16
```

#### 3.2 Visualize DE genes

There are thousands of genes that have are differentially expressed in a linear progression of tumor grades. As always, it's always good to visually check some of these genes. For example, we could plot the top 100 DE genes, the 1000 to 1100 top DE genes, etc. The expression can be visualized at different points. We could visualize the raw expression counts (Figure 5), the voom-normalized expression (Figure 6) (Law, Chen, Shi, and Smyth, 2014), or the *cleaned* voom-normalized expression (Figure 7). The last one is the normalized expression where we regress out the effects of the adjustment covariates. This can be done using the `cleaningY()` function from *jaffelab*.

In the following code, we first computed the *cleaned* normalized expression protecting the intercept term as well as the linear and quadratic trend terms. We also write a function that we can use to select which genes to plot as well as actually make the visualization with some nice features (colors, jitter points, linear trend line).

```
## Regress out sex, age and pathology from the gene expression
cleaned_expr <- cleaningY(gene_voom$E, mod, P = 3)

## gene plotting function
plot_gene <- function(ii, type = 'cleaned', sign = 'any') {
    ## Keep the jitter reproducible
    set.seed(20180203)

    ## Order by FDR and subset by logFC sign if necessary
    if(sign == 'any') {
        fdr_sorted <- with(stats_linear, gene_id[order(adj.P.Val)])
    } else {
        fdr_sorted <- with(stats_linear[sign(stats_linear$logFC) == sign, ],
            gene_id[order(adj.P.Val)])
    }

    ## Get the actual gene it matches originally
    i <- match(fdr_sorted[ii], names(rowRanges(has_patho)))
```

```
    ## Define what type of expression we are looking at
    if(type == 'cleaned') {
        y <- cleaned_expr[i, ]
        ylab <- 'Normalized Expr: age, sex, pathology removed'
    } else if (type == 'norm') {
        y <- gene_voom$E[i, ]
        ylab <- 'Normalized Expr'
    } else if (type == 'raw') {
        y <- dge$counts[i, ]
        ylab <- 'Raw Expr'
    }
    ylim <- abs(range(y)) * c(0.95, 1.05) * sign(range(y))

    ## Plot components
    x <- ordered(has_patho$clinical_stage_1)
    title <- with(stats_linear, paste(gene_id[i], symbol[i], 'FDR',
        signif(adj.P.Val[i], 3)))

    ## Make the plot ^^
    plot(y ~ x, xlab = 'Tumor grade', ylab = ylab, outline = FALSE,
        ylim = ylim, main = title)
    points(y ~ jitter(as.integer(x), 0.6),
        bg = c("#E69F00", "#009E73", "#D55E00")[as.integer(x)], pch = 21)
    abline(lm(y ~ as.integer(x)), lwd = 3, col = "#CC79A7")
}
```

Having built our plotting function, we can now visualize the top gene as shown in Figures 5, 6 and 7. In this case, there's not a large difference between the cleaned expression in Figure 7 and the normalized expression in Figure 6. From GeneCards we can see that the *SMC4* gene plays a role in the structural maintenance of chromosomes, which make sense in our context. Figure 8 shows the top DE gene with a decreasing expression trend across tumor grade progression. *CCNI2* is a paralog of *CCNI* which has been implicated in mitosis.

```
## Visualize the top gene
plot_gene(1, 'raw')
```

```
plot_gene(1, 'norm')
```

```
plot_gene(1)
```

```
## Visualize top gene with a downward trend
plot_gene(1, sign = '-1')
```

We are not experts in gliomas, but maybe your colleagues are and might recognize important genes. You can use the following code to make plots of some of the top DE genes in both directions and share the images with them to get feedback. Check the top50_increasing and top50_decreasing genes in the linked PDF files.

```
## Plot the top 50 increasing and decreasing genes
pdf('top50_increasing.pdf')
for(i in seq_len(50)) plot_gene(i, sign = '1')
```

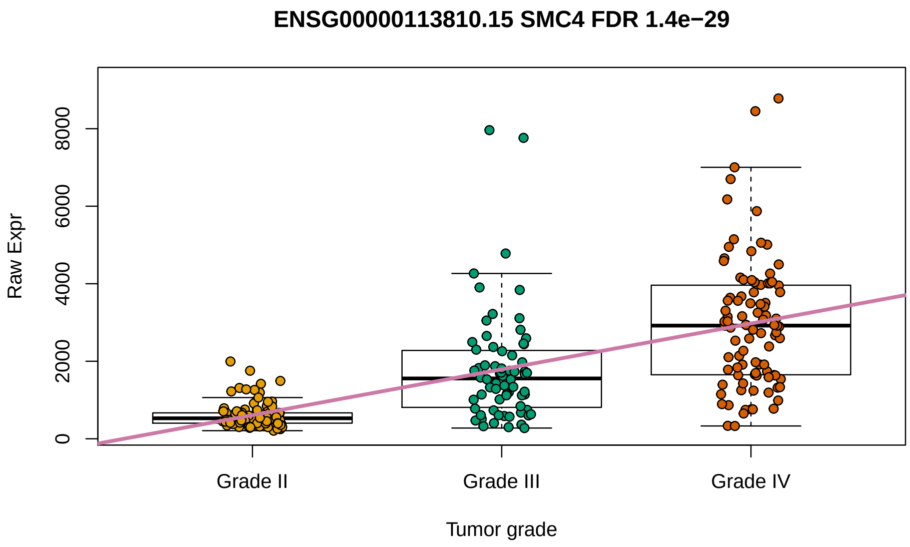

**Figure 5:** Raw expression for the top DE gene

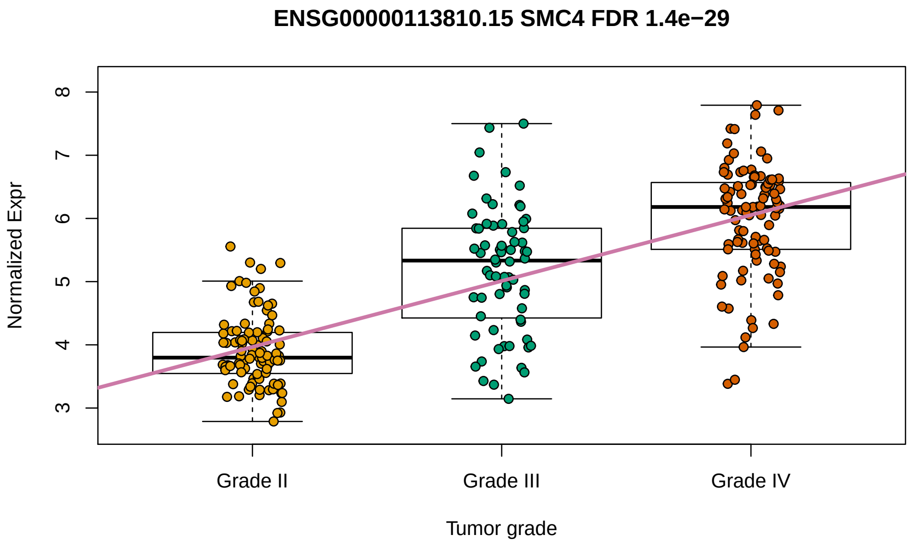

**Figure 6:** Voom-normalized expression for the top DE gene

```
dev.off()
## pdf
##   2

pdf('top50_decreasing.pdf')
for(i in seq_len(50)) plot_gene(i, sign = '-1')
dev.off()
## pdf
##   2
```

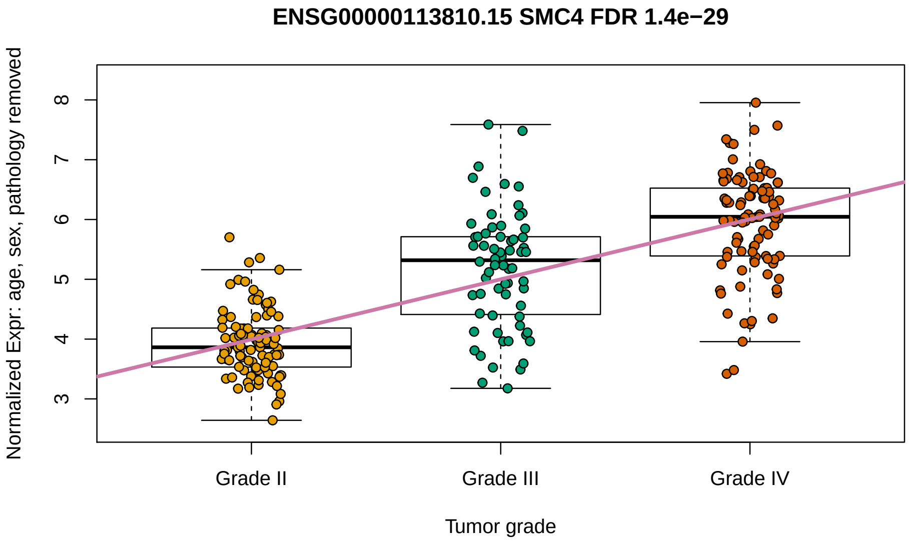

**Figure 7:** **Cleaned voom-normalized expression for the top DE gene**

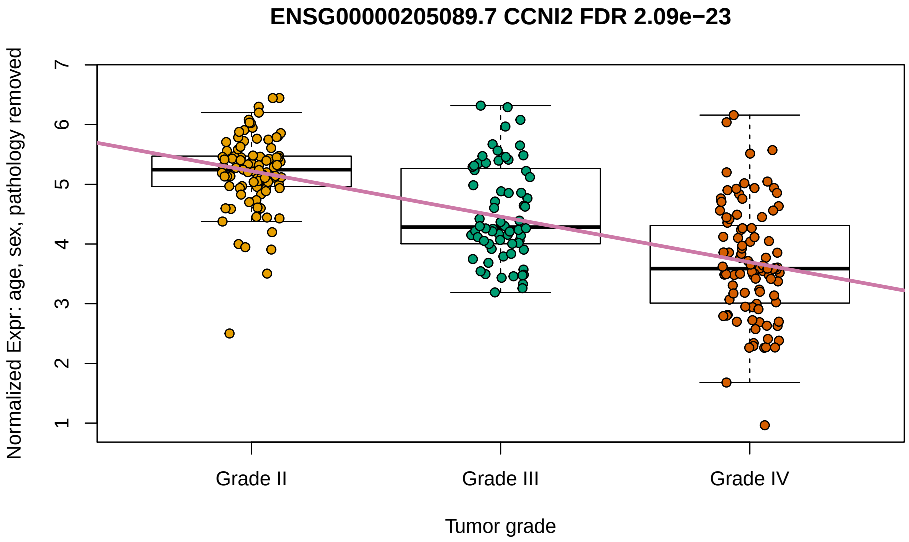

**Figure 8:** **Cleaned voom-normalized expression for the top DE gene with a decreasing trend**

#### 3.3 Gene ontology

Rather than look at the GeneCards for each gene, we can explore which gene ontologies are enriched in the DE genes that have a decreasing and an increasing trend with tumor grade progression. We can use *clusterProfiler* for this exploratory task[2].

```
library('clusterProfiler')
```

We need to extract the gene ids for our sets of genes of interest. Lets explore again the contents of the `stats_linear` object we created earlier. In the `gene_id` column we have the Gencode ids, which can be converted to ENSEMBL gene ids that *clusterProfiler* can then use.

[2] If you haven't done gene ontology enrichment analyses before check the vignette at bioconductor.org/packages/clusterProfiler.

```
head(stats_linear)
##                      seqnames     start       end  width strand
## ENSG00000000003.14     chrX 100627109 100639991  12883      -
## ENSG00000000005.5      chrX 100584802 100599885  15084      +
## ENSG00000000419.12    chr20  50934867  50958555  23689      -
## ENSG00000000457.13     chr1 169849631 169894267  44637      -
## ENSG00000000460.16     chr1 169662007 169854080 192074      +
## ENSG00000000938.12     chr1  27612064  27635277  23214      -
##                               gene_id bp_length   symbol       logFC
## ENSG00000000003.14 ENSG00000000003.14      4535   TSPAN6  0.24561059
## ENSG00000000005.5   ENSG00000000005.5      1610     TNMD -0.03870644
## ENSG00000000419.12 ENSG00000000419.12      1207     DPM1  0.37619802
## ENSG00000000457.13 ENSG00000000457.13      6883    SCYL3  0.10701765
## ENSG00000000460.16 ENSG00000000460.16      5967 C1orf112  0.36815450
## ENSG00000000938.12 ENSG00000000938.12      3474      FGR  0.29567134
##                      AveExpr          t      P.Value    adj.P.Val         B
## ENSG00000000003.14  5.402896  2.9838609 3.122016e-03 6.662499e-03 -2.879919
## ENSG00000000005.5  -2.971041 -0.1571861 8.752223e-01 9.044552e-01 -6.242625
## ENSG00000000419.12  4.238931  7.7909740 1.673109e-13 3.263390e-12 20.009519
## ENSG00000000457.13  4.018716  3.2001746 1.547119e-03 3.519785e-03 -2.103600
## ENSG00000000460.16  3.428671  7.2552308 4.792806e-12 6.609176e-11 16.798772
## ENSG00000000938.12  2.899634  2.6296133 9.066692e-03 1.745415e-02 -3.540602
```

With the following code we extract all the DE genes at a FDR of 1% that have an increasing or a decreasing trend. The code comments include a way you could further subset these genes to look at say the top 200 DE genes in each direction. We will use as our *universe* of genes all the genes that passed our low expression filter.

```
## Get ENSEMBL gene ids for all the DE genes with a decreasing and an
## increasing trend with tumor grade progression
de_genes <- lapply(c('-1', '1'), function(s) {
    ens <- with(stats_linear, gene_id[sign(logFC) == s & adj.P.Val < 0.01])
    ## Code if you wanted the top 200 instead
    #ens <- with(stats_linear[sign(stats_linear$logFC) == s, ],
    #    head(gene_id[order(adj.P.Val)], 200))
    ens <- gsub('\\..*', '', ens)
    return(ens)
})
names(de_genes) <- c('decreasing', 'increasing')
uni <- with(stats_linear, gsub('\\..*', '', gene_id))
```

Now that we have our `list` object with the set of genes with a decreasing or an increasing trend as well as our set of universe genes, we can compare the sets using `compareCluster()`. We will check the biological process, molecular function and cellular component ontologies.

```
## Which GO terms are enriched?
go_comp <- lapply(c('BP', 'MF', 'CC'), function(bp) {
    message(paste(Sys.time(), 'processing', bp))
    compareCluster(de_genes, fun = "enrichGO",
        universe = uni, OrgDb = 'org.Hs.eg.db',
        ont = bp, pAdjustMethod = "BH",
        pvalueCutoff  = 0.05, qvalueCutoff  = 0.05,
```

```
        readable= TRUE, keyType = 'ENSEMBL')
})
## 2018-03-03 10:54:07 processing BP
## 2018-03-03 10:56:32 processing MF
## 2018-03-03 10:58:00 processing CC
names(go_comp) <- c('Biological Process', 'Molecular Function',
    'Cellular Component')
```

Now that we have the data for each of the ontologies we can visualize the results using `clusterProfiler::plot()`. Figure 9 shows the enriched biological process terms where we see terms enriched for DNA replication and chromosome segregation in the genes with an increasing expression relationship with grade tumor progression. Intuitively this makes sense since gliomas are a type of cancer. The enriched molecular function ontology terms show in Figure 10 reflect the same picture with transmembrane transporters enriched in the genes with a decreasing expression association with grade tumor progression. Figure 11 shows the enriched cellular components with chromosome-releated terms related with the genes that have a higher expression as tumor progression advances. This is related to the findings in the original study where they focused in gene fusions (Bao, Chen, Yang, Zhang, et al., 2014).

```
## Visualize enriched GO terms
xx <- lapply(names(go_comp), function(bp) {
    print(plot(go_comp[[bp]], title = paste(bp, 'ontology'), font.size = 15))
    return(NULL)
})
```

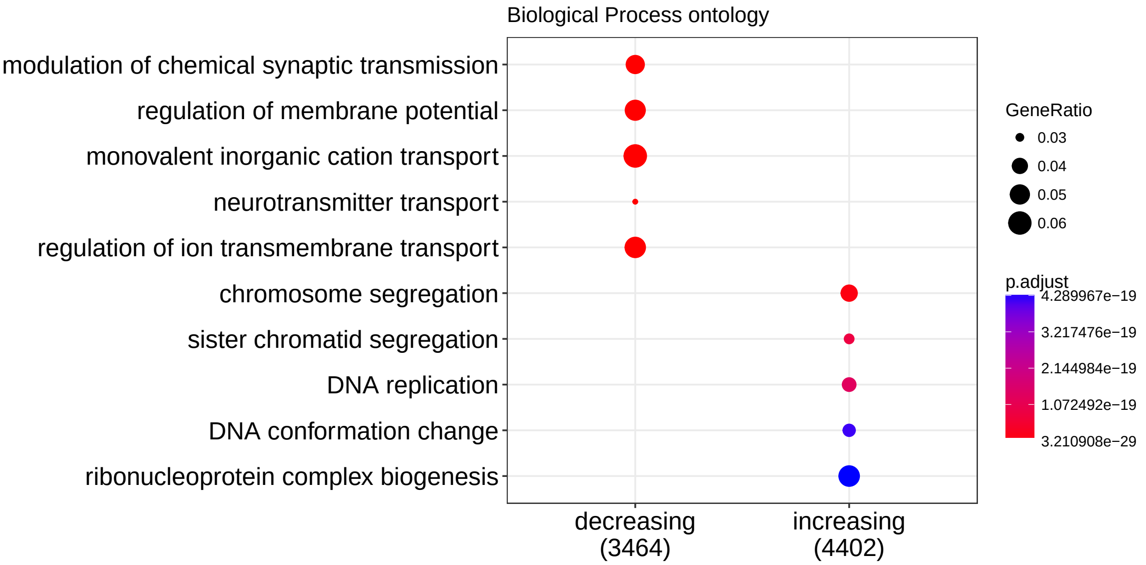

**Figure 9:** **Enriched biological process ontology terms**

We can finally save our exploratory results in case we want to carry out more analyses with them later on.

```
## Save results
save(stats_linear, stats_quad, go_comp, file = 'example_results.Rdata')
```

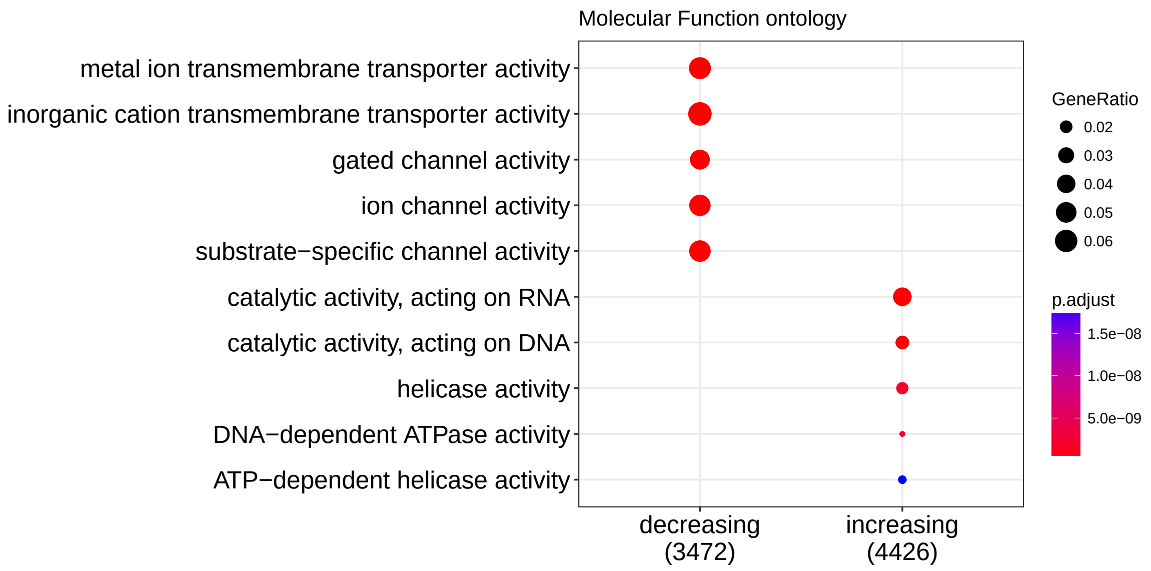

**Figure 10:** **Enriched molecular function ontology terms**

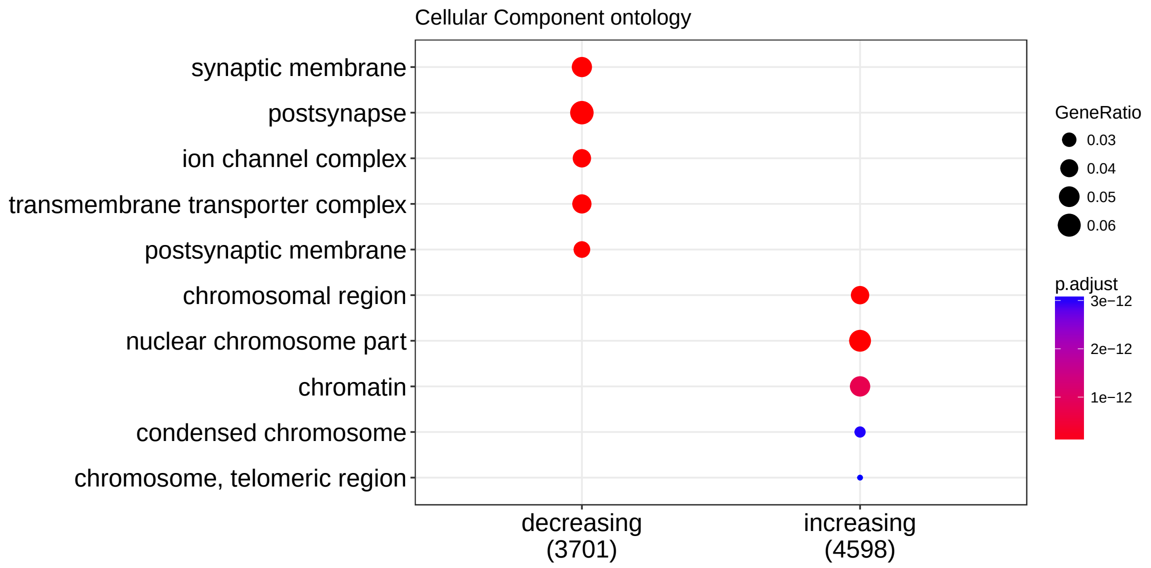

**Figure 11:** **Enriched cellular component ontology terms**

### 4 Conclusions

In this document we showed how you can download expression data from `recount2` using the *recount* package and add the sample metadata from `recount_brain`. We then illustrated how both the sample metadata and expression data can be used to explore a biological question of interest. We identified 6116 and 6438 differentially expressed genes at a FDR of 1% with decreasing and increasing linear trends in expression as tumor grade progresses while adjusting for age (in years), sex, and pathology (IDH1 mutation presence/absence).

### Reproducibility

```
## Reproducibility information
Sys.time()
## [1] "2018-03-03 11:22:53 EST"
proc.time()
##    user  system elapsed
##  97.124   8.554 113.460
options(width = 120)
devtools::session_info()
## Session info -------------------------------------------------------------------------------------------------
##  setting  value
##  version  R Under development (unstable) (2017-11-29 r73789)
##  system   x86_64, darwin15.6.0
##  ui       X11
##  language (EN)
##  collate  en_US.UTF-8
##  tz       America/New_York
##  date     2018-03-03
## Packages -----------------------------------------------------------------------------------------------------
##  package              * version   date       source
##  acepack                1.4.1     2016-10-29 CRAN (R 3.5.0)
##  AnnotationDbi          1.41.4    2017-12-11 Bioconductor
##  assertthat             0.2.0     2017-04-11 CRAN (R 3.5.0)
##  backports              1.1.2     2017-12-13 CRAN (R 3.5.0)
##  base                 * 3.5.0     2017-11-29 local
##  base64enc              0.1-3     2015-07-28 CRAN (R 3.5.0)
##  bibtex                 0.4.2     2017-06-30 CRAN (R 3.5.0)
##  bindr                  0.1       2016-11-13 CRAN (R 3.5.0)
##  bindrcpp               0.2       2017-06-17 CRAN (R 3.5.0)
##  Biobase              * 2.39.2    2018-01-25 Bioconductor
##  BiocGenerics         * 0.25.3    2018-02-09 Bioconductor
##  BiocParallel         * 1.13.1    2017-12-31 Bioconductor
##  BiocStyle            * 2.7.8     2018-01-20 Bioconductor
##  biomaRt                2.35.12   2018-03-03 Bioconductor
##  Biostrings             2.47.9    2018-02-10 Bioconductor
##  bit                    1.1-12    2014-04-09 CRAN (R 3.5.0)
##  bit64                  0.9-7     2017-05-08 CRAN (R 3.5.0)
##  bitops                 1.0-6     2013-08-17 CRAN (R 3.5.0)
##  blob                   1.1.0     2017-06-17 CRAN (R 3.5.0)
##  bookdown               0.7       2018-02-18 CRAN (R 3.5.0)
##  BSgenome               1.47.5    2018-02-13 Bioconductor
##  bumphunter             1.21.0    2017-10-31 Bioconductor
##  checkmate              1.8.5     2017-10-24 CRAN (R 3.5.0)
##  cluster                2.0.6     2017-03-10 CRAN (R 3.5.0)
##  clusterProfiler      * 3.7.0     2017-10-31 Bioconductor
##  codetools              0.2-15    2016-10-05 CRAN (R 3.5.0)
##  colorout             * 1.1-3     2017-11-29 Github (jalvesaq/colorout@e2a175c)
##  colorspace             1.3-2     2016-12-14 CRAN (R 3.5.0)
##  compiler               3.5.0     2017-11-29 local
```

```
##  curl                     3.1       2017-12-12 CRAN (R 3.5.0)
##  data.table               1.10.4-3  2017-10-27 CRAN (R 3.5.0)
##  datasets               * 3.5.0     2017-11-29 local
##  DBI                      0.8       2018-03-02 CRAN (R 3.5.0)
##  DelayedArray           * 0.5.22    2018-03-02 Bioconductor
##  derfinder                1.13.8    2018-02-24 cran (@1.13.8)
##  derfinderHelper          1.13.0    2017-10-31 Bioconductor
##  devtools                 1.13.5    2018-02-18 CRAN (R 3.5.0)
##  digest                   0.6.15    2018-01-28 CRAN (R 3.5.0)
##  DO.db                    2.9       2017-11-30 Bioconductor
##  doRNG                    1.6.6     2017-04-10 CRAN (R 3.5.0)
##  DOSE                   * 3.5.0     2017-10-31 Bioconductor
##  downloader               0.4       2015-07-09 CRAN (R 3.5.0)
##  dplyr                    0.7.4     2017-09-28 CRAN (R 3.5.0)
##  edgeR                  * 3.21.9    2018-02-27 Bioconductor
##  evaluate                 0.10.1    2017-06-24 CRAN (R 3.5.0)
##  fastmatch                1.1-0     2017-01-28 CRAN (R 3.5.0)
##  fgsea                    1.5.2     2018-02-24 Bioconductor
##  foreach                  1.4.4     2017-12-12 CRAN (R 3.5.0)
##  foreign                  0.8-70    2017-11-28 CRAN (R 3.5.0)
##  Formula                  1.2-2     2017-07-10 CRAN (R 3.5.0)
##  GenomeInfoDb           * 1.15.5    2018-02-04 Bioconductor
##  GenomeInfoDbData         1.1.0     2017-12-15 Bioconductor
##  GenomicAlignments        1.15.12   2018-02-11 Bioconductor
##  GenomicFeatures          1.31.10   2018-02-10 Bioconductor
##  GenomicFiles             1.15.2    2018-02-09 Bioconductor
##  GenomicRanges          * 1.31.22   2018-02-16 Bioconductor
##  GEOquery                 2.47.18   2018-03-02 Bioconductor
##  ggplot2                  2.2.1     2016-12-30 CRAN (R 3.5.0)
##  glue                     1.2.0     2017-10-29 CRAN (R 3.5.0)
##  GO.db                    3.5.0     2017-11-30 Bioconductor
##  GOSemSim                 2.5.1     2018-02-10 Bioconductor
##  graphics               * 3.5.0     2017-11-29 local
##  grDevices              * 3.5.0     2017-11-29 local
##  grid                     3.5.0     2017-11-29 local
##  gridExtra                2.3       2017-09-09 CRAN (R 3.5.0)
##  gtable                   0.2.0     2016-02-26 CRAN (R 3.5.0)
##  Hmisc                    4.1-1     2018-01-03 CRAN (R 3.5.0)
##  hms                      0.4.1     2018-01-24 CRAN (R 3.5.0)
##  htmlTable                1.11.2    2018-01-20 CRAN (R 3.5.0)
##  htmltools                0.3.6     2017-04-28 CRAN (R 3.5.0)
##  htmlwidgets              1.0       2018-01-20 CRAN (R 3.5.0)
##  httr                     1.3.1     2017-08-20 CRAN (R 3.5.0)
##  igraph                   1.1.2     2017-07-21 CRAN (R 3.5.0)
##  IRanges                * 2.13.28   2018-02-24 cran (@2.13.28)
##  iterators                1.0.9     2017-12-12 CRAN (R 3.5.0)
##  jaffelab               * 0.99.18   2018-02-27 Github (LieberInstitute/jaffelab@a8e6430)
##  jsonlite                 1.5       2017-06-01 CRAN (R 3.5.0)
##  knitcitations          * 1.0.8     2017-07-04 CRAN (R 3.5.0)
##  knitr                    1.20      2018-02-20 CRAN (R 3.5.0)
##  labeling                 0.3       2014-08-23 CRAN (R 3.5.0)
```

```
##  lattice               0.20-35   2017-03-25 CRAN (R 3.5.0)
##  latticeExtra          0.6-28    2016-02-09 CRAN (R 3.5.0)
##  lazyeval              0.2.1     2017-10-29 CRAN (R 3.5.0)
##  limma               * 3.35.12   2018-02-22 Bioconductor
##  locfit                1.5-9.1   2013-04-20 CRAN (R 3.5.0)
##  lubridate             1.7.3     2018-02-27 CRAN (R 3.5.0)
##  magrittr              1.5       2014-11-22 CRAN (R 3.5.0)
##  Matrix                1.2-12    2017-11-20 CRAN (R 3.5.0)
##  matrixStats         * 0.53.1    2018-02-11 CRAN (R 3.5.0)
##  memoise               1.1.0     2017-04-21 CRAN (R 3.5.0)
##  methods             * 3.5.0     2017-11-29 local
##  munsell               0.4.3     2016-02-13 CRAN (R 3.5.0)
##  nnet                  7.3-12    2016-02-02 CRAN (R 3.5.0)
##  parallel            * 3.5.0     2017-11-29 local
##  pillar                1.2.1     2018-02-27 CRAN (R 3.5.0)
##  pkgconfig             2.0.1     2017-03-21 CRAN (R 3.5.0)
##  pkgmaker              0.22      2014-05-14 CRAN (R 3.5.0)
##  plyr                  1.8.4     2016-06-08 CRAN (R 3.5.0)
##  prettyunits           1.0.2     2015-07-13 CRAN (R 3.5.0)
##  progress              1.1.2     2016-12-14 CRAN (R 3.5.0)
##  purrr                 0.2.4     2017-10-18 CRAN (R 3.5.0)
##  qvalue                2.11.0    2017-10-31 Bioconductor
##  R6                    2.2.2     2017-06-17 CRAN (R 3.5.0)
##  rafalib             * 1.0.0     2015-08-09 CRAN (R 3.5.0)
##  RColorBrewer          1.1-2     2014-12-07 CRAN (R 3.5.0)
##  Rcpp                  0.12.15   2018-01-20 CRAN (R 3.5.0)
##  RCurl                 1.95-4.10 2018-01-04 CRAN (R 3.5.0)
##  readr                 1.1.1     2017-05-16 CRAN (R 3.5.0)
##  recount             * 1.5.9     2018-03-01 Github (leekgroup/recount@458d4f2)
##  RefManageR            0.14.20   2017-08-17 CRAN (R 3.5.0)
##  registry              0.5       2017-12-03 CRAN (R 3.5.0)
##  rentrez               1.2.0     2018-02-12 CRAN (R 3.5.0)
##  reshape2              1.4.3     2017-12-11 CRAN (R 3.5.0)
##  rlang                 0.2.0     2018-02-20 CRAN (R 3.5.0)
##  rmarkdown             1.9       2018-03-01 CRAN (R 3.5.0)
##  RMySQL                0.10.14   2018-02-26 CRAN (R 3.5.0)
##  rngtools              1.2.4     2014-03-06 CRAN (R 3.5.0)
##  rpart                 4.1-13    2018-02-23 CRAN (R 3.5.0)
##  rprojroot             1.3-2     2018-01-03 CRAN (R 3.5.0)
##  Rsamtools             1.31.3    2018-02-02 Bioconductor
##  RSQLite               2.0       2017-06-19 CRAN (R 3.5.0)
##  rstudioapi            0.7       2017-09-07 CRAN (R 3.5.0)
##  rtracklayer           1.39.9    2018-02-11 Bioconductor
##  rvcheck               0.0.9     2017-07-10 CRAN (R 3.5.0)
##  S4Vectors           * 0.17.36   2018-03-03 Bioconductor
##  scales                0.5.0     2017-08-24 CRAN (R 3.5.0)
##  segmented             0.5-3.0   2017-11-30 CRAN (R 3.5.0)
##  splines               3.5.0     2017-11-29 local
##  stats               * 3.5.0     2017-11-29 local
##  stats4              * 3.5.0     2017-11-29 local
##  stringi               1.1.6     2017-11-17 CRAN (R 3.5.0)
```

```
##  stringr                1.3.0     2018-02-19 CRAN (R 3.5.0)
##  SummarizedExperiment * 1.9.15    2018-02-24 cran (@1.9.15)
##  survival               2.41-3    2017-04-04 CRAN (R 3.5.0)
##  tibble                 1.4.2     2018-01-22 CRAN (R 3.5.0)
##  tidyr                  0.8.0     2018-01-29 CRAN (R 3.5.0)
##  tools                  3.5.0     2017-11-29 local
##  utils                * 3.5.0     2017-11-29 local
##  VariantAnnotation      1.25.12   2018-01-25 Bioconductor
##  withr                  2.1.1     2017-12-19 CRAN (R 3.5.0)
##  xfun                   0.1       2018-01-22 CRAN (R 3.5.0)
##  XML                    3.98-1.10 2018-02-19 CRAN (R 3.5.0)
##  xml2                   1.2.0     2018-01-24 CRAN (R 3.5.0)
##  xtable                 1.8-2     2016-02-05 CRAN (R 3.5.0)
##  XVector                0.19.9    2018-02-28 Bioconductor
##  yaml                   2.1.17    2018-02-27 CRAN (R 3.5.0)
##  zlibbioc               1.25.0    2017-10-31 Bioconductor
```

### References

The analyses were made possible thanks to:

- R (R Core Team, 2017)
- *BiocStyle* (Oleś, Morgan, and Huber, 2018)
- *clusterProfiler* (Yu, Wang, Han, and He, 2012)
- *devtools* (Wickham, Hester, and Chang, 2018)
- *edgeR* (Robinson, McCarthy, and Smyth, 2010; McCarthy, J., Chen, Yunshun, et al., 2012)
- *jaffelab* (Collado-Torres and Jaffe, 2018)
- *knitcitations* (Boettiger, 2017)
- *knitr* (Xie, 2014)
- *limma* (Ritchie, Phipson, Wu, Hu, et al., 2015; Law, Chen, Shi, and Smyth, 2014)
- *recount* (Collado-Torres, Nellore, Kammers, Ellis, et al., 2017; Collado-Torres, Nellore, and Jaffe, 2017)
- *rmarkdown* (Allaire, Xie, McPherson, Luraschi, et al., 2018)

Full bibliography file.

[1] J. Allaire, Y. Xie, J. McPherson, J. Luraschi, et al. rmarkdown: Dynamic Documents for R. R package version 1.9. 2018. URL: https://CRAN.R-project.org/package=rmarkdown.

[2] Z. Bao, H. Chen, M. Yang, C. Zhang, et al. “RNA-seq of 272 gliomas revealed a novel, recurrentPTPRZ1-METfusion transcript in secondary glioblastomas”. In: Genome Research 24.11 (Aug. 2014), pp. 1765–1773. DOI: 10.1101/gr.165126.113. URL: https://doi.org/10.1101/gr.165126.113.

[3] C. Boettiger. knitcitations: Citations for ‘Knitr’ Markdown Files. R package version 1.0.8. 2017. URL: https://CRAN.R-project.org/package=knitcitations.

[4] L. Collado-Torres and A. E. Jaffe. jaffelab: Commonly used functions by the Jaffe lab. R package version 0.99.18. 2018. URL: https://github.com/LieberInstitute/jaffelab.

[5] L. Collado-Torres, A. Nellore and A. E. Jaffe. "recount workflow: Accessing over 70,000 human RNA-seq samples with Bioconductor [version 1; referees: 1 approved, 2 approved with reservations]". In: F1000Research (2017). DOI: 10.12688/f1000research.12223.1. URL: https://f1000research.com/articles/6-1558/v1.

[6] L. Collado-Torres, A. Nellore, K. Kammers, S. E. Ellis, et al. "Reproducible RNA-seq analysis using recount2". In: Nature Biotechnology (2017). DOI: 10.1038/nbt.3838. URL: http://www.nature.com/nbt/journal/v35/n4/full/nbt.3838.html.

[7] C. Law, Y. Chen, W. Shi and G. Smyth. "Voom: precision weights unlock linear model analysis tools for RNA-seq read counts". In: Genome Biology 15 (2014), p. R29.

[8] McCarthy, D. J., Chen, Yunshun, et al. "Differential expression analysis of multifactor RNA-Seq experiments with respect to biological variation". In: Nucleic Acids Research 40.10 (2012), pp. 4288-4297.

[9] A. Oleś, M. Morgan and W. Huber. BiocStyle: Standard styles for vignettes and other Bioconductor documents. R package version 2.7.8. 2018. URL: https://github.com/Bioconductor/BiocStyle.

[10] R Core Team. R: A Language and Environment for Statistical Computing. R Foundation for Statistical Computing. Vienna, Austria, 2017. URL: https://www.R-project.org/.

[11] M. E. Ritchie, B. Phipson, D. Wu, Y. Hu, et al. "limma powers differential expression analyses for RNA-sequencing and microarray studies". In: Nucleic Acids Research 43.7 (2015), p. e47.

[12] M. D. Robinson, D. J. McCarthy and G. K. Smyth. "edgeR: a Bioconductor package for differential expression analysis of digital gene expression data". In: Bioinformatics 26.1 (2010), pp. 139-140.

[13] H. Wickham, J. Hester and W. Chang. devtools: Tools to Make Developing R Packages Easier. R package version 1.13.5. 2018. URL: https://CRAN.R-project.org/package=devtools.

[14] Y. Xie. "knitr: A Comprehensive Tool for Reproducible Research in R". In: Implementing Reproducible Computational Research. Ed. by V. Stodden, F. Leisch and R. D. Peng. ISBN 978-1466561595. Chapman and Hall/CRC, 2014. URL: http://www.crcpress.com/product/isbn/9781466561595.

[15] G. Yu, L. Wang, Y. Han and Q. He. "clusterProfiler: an R package for comparing biological themes among gene clusters". In: OMICS: A Journal of Integrative Biology 16.5 (2012), pp. 284-287. DOI: 10.1089/omi.2011.0118.
