## Supplementary File 2 for "*recount-brain*: a curated repository of human brain RNA-seq datasets metadata"

### Using recount_brain to explore relationships with post-mortem interval

**3 March 2018**

**Abstract**

This is an example of how you can use recount_brain to check if some results from another study are replicated in brain samples from the SRA present in recount2. This example also shows how you can quantify the percent of mitochondrial transcription and options for comparing two sets of SRA studies.

### 1 Introduction

This document is an example of how you can use `recount_brain`. In a recent paper by Ferreira *et al.* called "The effects of death and post-mortem cold ischemia on human tissue transcriptomes" the authors explore associations between expression and post-mortem interval (PMI) (Ferreira, Muñoz-Aguirre, Reverter, Godinho, et al., 2018). We asked ourselves if PMI was recorded in any brain samples in `recount2` (Collado-Torres, Nellore, Kammers, Ellis, et al., 2017) and got started using `recount_brain`. Here we will check the expression trends of some of the genes highlighted by Ferreira *et al.* in Figure 2d and the association between PMI and percent of reads aligning to the mitochondrial chromosome from Figure 4b and 4c. We will use the gene count data as well as the BigWig files provided by `recount2` (Collado-Torres, Nellore, Kammers, Ellis, et al., 2017) and combine this information with the sample metadata from `recount_brain`.

This example shows how you can use the full `recount_brain` table to identify samples/studies of interest and the diversity of expression data available in `recount2`. This can be useful prior to publication if you want to check if your results are replicated in other data sets, for post-publication meta-analyses and for education purposes.

### 2 Identify samples and studies of interest

Just like in the example with data from SRP027373, we will e will be using many functions from the *recount* package, lets load it first[1].

[1] If you are a first time `recount` user, we recommend first reading the package vignette at bioconductor.org/packages/recount.

```
## Load the package
library('recount')
```

#### 2.1 Download `recount_brain`

Now that we have loaded *recount*, lets obtain the full `recount_brain` table. This can be done via the `add_metadata()` function when we don't specify a `rse` argument.

```
recount_brain <- add_metadata(source = 'recount_brain_v1')
## 2018-03-03 11:38:15 downloading the recount_brain metadata to /var/folders/cx/n9s558kx6fb7jf5z_pgszgb8000
## Loading objects:
##   recount_brain
dim(recount_brain)
## [1] 4431   48
```

#### 2.2 Explore `recount_brain` data

The full table includes samples not present in `recount2`, so lets drop those.

```
## Some studies included non-human Illumina RNA-seq data,
## which by design are absent in recount2
recount_brain <- recount_brain[recount_brain$present_in_recount, ]
```

##### Using recount_brain to explore relationships with post-mortem interval

```
dim(recount_brain)
## [1] 3214   48
```

Next, we can find which variables have information about post-mortem interval (PMI).

```
colnames(recount_brain)[grep('pmi', colnames(recount_brain))]
## [1] "pmi_units" "pmi"
```

We can see that information about PMI is stored in two variables: the PMI value and the PMI unit. Lets keep only the samples where both of them are observed: most the samples with PMI values have a PMI unit.

```
with(recount_brain, addmargins(table(
    'PMI observed' = !is.na(pmi),
    'Has PMI units' = !is.na(pmi_units)
    ))
)
##              Has PMI units
## PMI observed FALSE TRUE  Sum
##        FALSE  2931    1 2932
##        TRUE      0  282  282
##        Sum    2931  283 3214
recount_brain <- recount_brain[with(recount_brain,
    !is.na(pmi) & !is.na(pmi_units)), ]
dim(recount_brain)
## [1] 282  48
```

Now that we have selected our samples of interest, lets check how many different studies these samples come from. As we can see below it's 12 studies with a range of 8 to 73 samples in each study.

```
length(unique(recount_brain$sra_study_s))
## [1] 12
sort(table(recount_brain$sra_study_s))
##
## SRP056604 SRP055440 SRP032539 SRP032540 SRP007483 SRP048683 SRP019762
##         8         9        11        11        12        12        14
## ERP001304 SRP067645 SRP017933 SRP051844 SRP058181
##        18        21        25        68        73
```

We can also explore the range of PMI values and units. Actually, all of the PMI values are measured in hours as we can see below.

```
table(recount_brain$pmi_units)
##
## Hours
##   282
summary(recount_brain$pmi)
##    Min. 1st Qu.  Median    Mean 3rd Qu.    Max.
##    1.00    5.00   11.50   12.67   19.00   39.00
```

#### Using recount_brain to explore relationships with post-mortem interval

In Figure 1 we plot the distribution of PMI in hours across all the samples of interest from the 12 studies. There are a few peaks which could be due to some tissues having less diverse PMI values.

```
library('ggplot2')

## Used code from
## http://www.cookbook-r.com/Graphs/Plotting_distributions_(ggplot2)/
ggplot(recount_brain, aes(x = pmi)) +
    geom_histogram(aes(y=..density..),
        colour="black", fill="white") +
    geom_density(alpha=.2, fill="#FF6666") + xlab('PMI in hours') +
    theme_grey(base_size = 18)
```

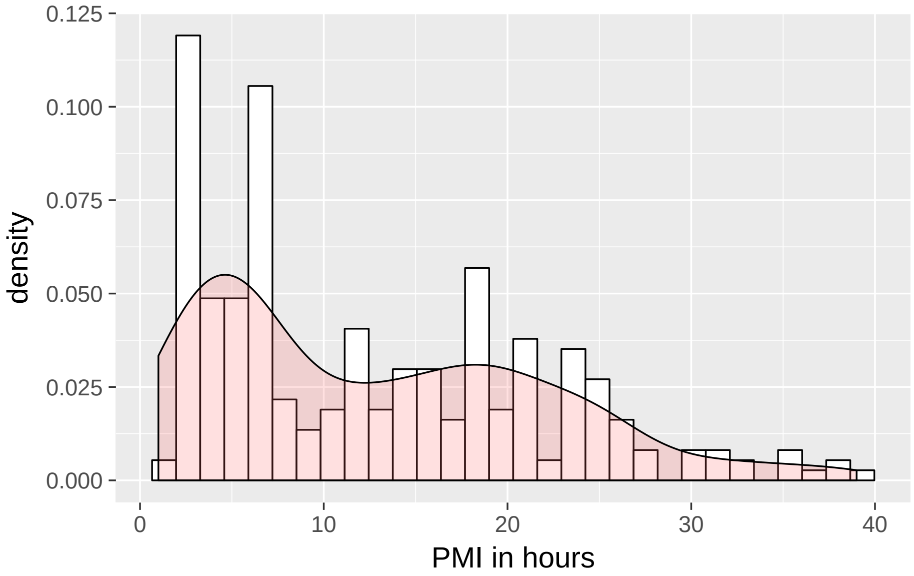

**Figure 1: Distribution of PMI in hours for all SRA studies with PMI information present in recount_brain**

Figure 2 shows the distribution of PMI in hours for each of the 12. Two of them stand out as they have no variation in PMI: studies SRP067645 and SRP055440.

```
ggplot(recount_brain, aes(y = pmi, x = sra_study_s)) +
    geom_boxplot() + coord_flip() + theme_bw(base_size = 18) +
    ylab('PMI in hours') + xlab('SRA Study ID')
```

```
recount_brain <- subset(recount_brain,
    !sra_study_s %in% c('SRP067645', 'SRP055440'))
## Final dimentions
dim(recount_brain)
## [1] 252  48
```

We can now remake Figure 1 having dropped the two studies with no PMI variability as shown in Figure 3. There's still a peak at low PMI values, most likely from the 8 samples from study SRP056604.

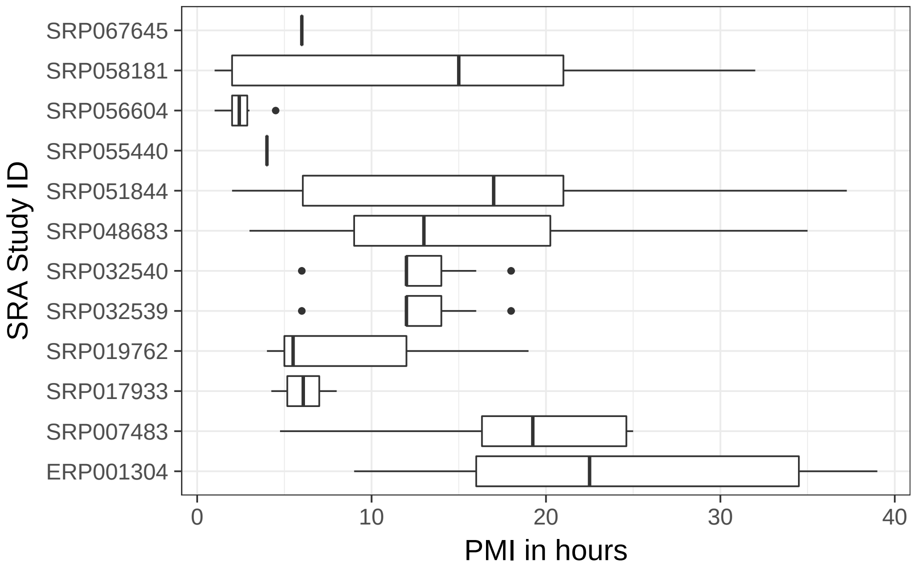

**Figure 2:** **Distribution of PMI in hours by study**

```
ggplot(recount_brain, aes(x = pmi)) +
    geom_histogram(aes(y=..density..),
        colour="black", fill="white") +
    geom_density(alpha=.2, fill="#FF6666") + xlab('PMI in hours') +
    theme_grey(base_size = 18)
```

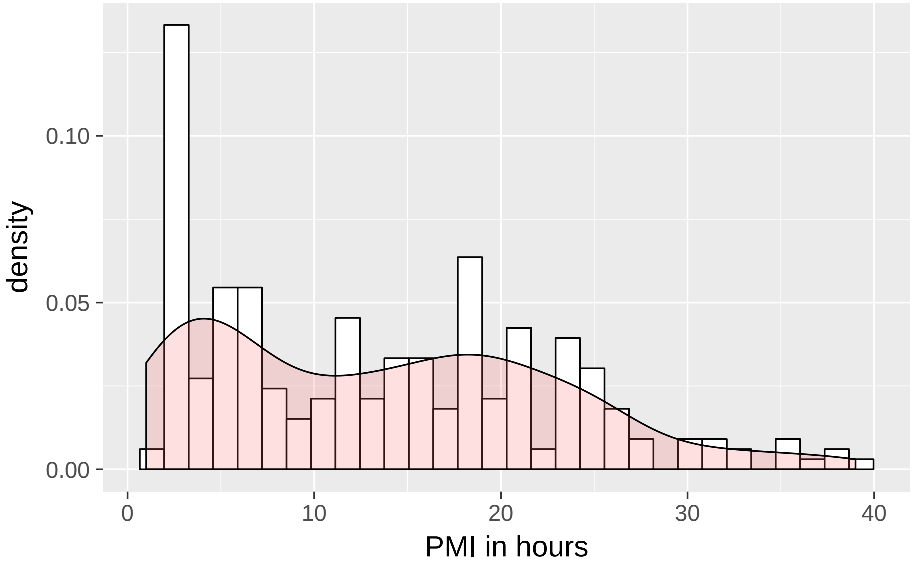

**Figure 3:** **Distribution of PMI in hours for all SRA studies with PMI information present in recount_brain and that have PMI variability**

Note that some of these studies use control samples and others focus on specific diseases. Figure 4 shows the distribution of PMI by disease status where we see no significant difference. Though it could play a role in it's relationship with expression.

##### Using recount_brain to explore relationships with post-mortem interval

```
## What is the disease status of our current samples?
table(recount_brain$disease_status, useNA = 'ifany')
##
## Control Disease
##     184      68

ggplot(recount_brain, aes(y = pmi, x = disease_status)) +
    geom_boxplot() + coord_flip() + theme_bw(base_size = 18) +
    ylab('PMI in hours') + xlab('Sample type')
```

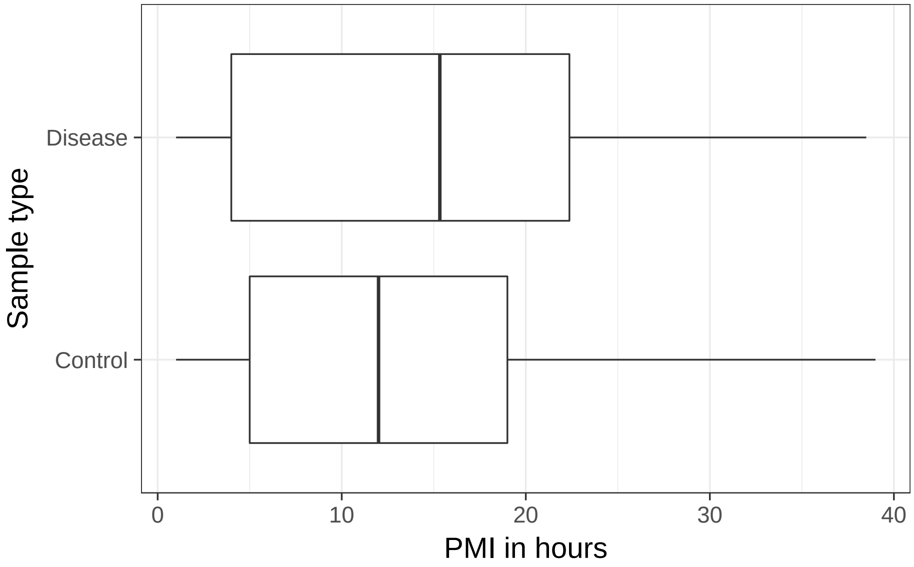

**Figure 4:** **Distribution of PMI in hours by disease status**

```
## Is there a significant difference in PMI by disease status?
with(recount_brain, t.test(pmi ~ disease_status))
##
##  Welch Two Sample t-test
##
## data:  pmi by disease_status
## t = -1.4532, df = 105.79, p-value = 0.1491
## alternative hypothesis: true difference in means is not equal to 0
## 95 percent confidence interval:
##  -4.864907  0.749606
## sample estimates:
## mean in group Control mean in group Disease
##              12.97897              15.03662
```

Awesome, we are now ready to get expression data!

### 3 Gene expression

In Figure 2d Ferreira *et al.* (Ferreira, Muñoz-Aguirre, Reverter, Godinho, et al., 2018) show the changes in gene expression for different tissues across several PMI intervals. We can download the gene count information from `recount2` using the `download_study()` function as shown next.

```
## Our studies of interest
studies <- unique(recount_brain$sra_study_s)

## Download the files if they are missing
if(any(!file.exists(file.path(studies, 'rse_gene.Rdata')))) {
    sapply(studies, download_study)
}
```

Now that we have the gene RangedSummarizedExperiment (RSE) objects in our computer, we can proceed to load them. Since each file has a `rse_gene` object we have to be careful when loading these files. One way to handle them is to load them in a loop function such as a `lapply()` and `return()` them to create a list of RSE objects. Since the sample metadata variables and gene information is the same for each of these objects, we can combine them into a single object using `cbind()`.

```
## Load each rse_gene object
rse_gene <- lapply(studies, function(s) {
    message(paste(Sys.time(), 'processing study', s))
    load(file.path(s, 'rse_gene.Rdata'), verbose = TRUE)
    return(rse_gene)
})
## 2018-03-03 11:38:22 processing study ERP001304
## Loading objects:
##   rse_gene
## 2018-03-03 11:38:22 processing study SRP032539
## Loading objects:
##   rse_gene
## 2018-03-03 11:38:23 processing study SRP032540
## Loading objects:
##   rse_gene
## 2018-03-03 11:38:23 processing study SRP048683
## Loading objects:
##   rse_gene
## 2018-03-03 11:38:24 processing study SRP051844
## Loading objects:
##   rse_gene
## 2018-03-03 11:38:24 processing study SRP007483
## Loading objects:
##   rse_gene
## 2018-03-03 11:38:25 processing study SRP056604
## Loading objects:
##   rse_gene
## 2018-03-03 11:38:26 processing study SRP058181
## Loading objects:
```

```
##   rse_gene
## 2018-03-03 11:38:26 processing study SRP017933
## Loading objects:
##   rse_gene
## 2018-03-03 11:38:27 processing study SRP019762
## Loading objects:
##   rse_gene
## Combine them into a single one
rse_gene <- do.call(cbind, rse_gene)
dim(rse_gene)
## [1] 58037   253
```

Looks like we have one extra sample than what we were working with, so lets drop it. Once that's done we have our merged RSE object with the data from all 10 studies.

```
## Keep only the samples we were working with
rse_gene <- rse_gene[, colnames(rse_gene) %in% recount_brain$run_s]

## Check that we are all ok
stopifnot(ncol(rse_gene) == nrow(recount_brain))

## Merge rse_gene object
rse_gene
## class: RangedSummarizedExperiment
## dim: 58037 252
## metadata(0):
## assays(1): counts
## rownames(58037): ENSG00000000003.14 ENSG00000000005.5 ...
##   ENSG00000283698.1 ENSG00000283699.1
## rowData names(3): gene_id bp_length symbol
## colnames(252): ERR103423 ERR103435 ... SRR786626 SRR786627
## colData names(21): project sample ... title characteristics
```

#### 3.1 Add `recount_brain` sample metadata

Now that we have the necessary gene information we can combine it with the sample metadata from `recount_brain`. We could either do this manually or simply use the `recount::add_metadata()` function.

```
## Add recount_brain metadata
dim(colData(rse_gene))
## [1] 252  21
rse_gene <- add_metadata(rse = rse_gene, source = 'recount_brain_v1')
## 2018-03-03 11:38:30 downloading the recount_brain metadata to /var/folders/cx/n9s558kx6fb7jf5z_pgszgb8000
## Loading objects:
##   recount_brain
## 2018-03-03 11:38:31 found 252 out of 252 samples in the recount_brain metadata
dim(colData(rse_gene))
## [1] 252  68
```

#### 3.2 PMI intervals

In Figure 2d Ferreira *et al.* (Ferreira, Muñoz-Aguirre, Reverter, Godinho, et al., 2018) use 5 sets of PMI intervals. We can reconstruct them with the `cut()` function and use the `right = FALSE` argument to make sure that the intervals match theirs: [left inclusive, right exclusive). In our case we don't have any PMIs in the [0, 1) hour interval so we won't be using it.

```
## Compute PMI intervals
colData(rse_gene)$pmilvl <- cut(rse_gene$pmi,
    c(0, 1, 4, 6, 15, max(rse_gene$pmi) + 1), right = FALSE)

## Check that the intervals match those from Figure 2d
table(colData(rse_gene)$pmilvl)
##
##   [0,1)   [1,4)   [4,6)  [6,15) [15,40)
##       0      47      26      63     116

## Rename to I1, I2, ..., I5
levels(colData(rse_gene)$pmilvl) <- paste0('I', 1:5)

## Drop empty interval
colData(rse_gene)$pmilvl <- droplevels(colData(rse_gene)$pmilvl)
```

#### 3.3 Normalize expression

In Figure 2d Ferreira *et al.* (Ferreira, Muñoz-Aguirre, Reverter, Godinho, et al., 2018) use $\log_2($ RPKM $+ 0.5$ ). We can compute RPKM values using the `getRPKM()` function from *recount*.

```
rpkm <- getRPKM(scale_counts(rse_gene))
```

In their case, all the expression data came from controls and from the same project. As a simple check, we can adjust the normalized expression for the disease status and project. We can do this with the `cleaningY()` function from *jaffelab* just like we did in the example with data from SRP027373. The disease status and study might have stronger effects than what this approach can remove, but it can help us explore the data.

```
library('jaffelab')
## You can install it with
# devtools::install_github('LieberInstitute/jaffelab')

mod <- with(colData(rse_gene),
    model.matrix(~ pmi + disease_status + project))
cleaned <- cleaningY(log2(rpkm + 0.5), mod, P = 2)
```

#### 3.4 Visualize example genes

In order to visualize the same genes as in Figure 2d (Ferreira, Muñoz-Aguirre, Reverter, Godinho, et al., 2018) we need to first find these genes in the `recount2` data. This gets a little bit tricky because the symbol information is stored in a `CharacterList` object.

```
## Short function for finding the genes
find_gene <- function(symb) {
    ## Note that rowRanges(rse_gene)$symbol is a CharacterList object
    finding_gene <- rowRanges(rse_gene)$symbol == symb
    i <- which(any(finding_gene))
    return(i)
}

## Find them! ^_^
gene_i <- sapply(c('RNASE2', 'EGR3', 'HBA1', 'CXCL2'), find_gene)
gene_i
## RNASE2.ENSG00000169385   EGR3.ENSG00000179388   HBA1.ENSG00000206172
##                  12598                  14687                  20421
##   CXCL2.ENSG00000081041
##                   1522
```

We finally have all the information we need:

- the gene expression data for the selected studies,
- the PMI interval in hours data,
- the gene IDs that match those used by Ferreira *et al.*

Lets make a function that can plot either the normalized gene expression as in the original work, or the normalize gene expression after removing the effects of disease status and study. We might also be interested in sub setting by study to ask if the global trend could have been seen before and whether it would have been consistent. Let's also add the p-value for the linear regression between the PMI intervals and gene expression.

```
## Our flexible plotting function.
## Mainly requires a gene symbol.
plot_gene <- function(symb, type = 'cleaned', study) {
    ## Keep the jitter reproducible
    set.seed(20180303)

    ## Find the gene based on the symbol
    i <- gene_i[grep(symb, names(gene_i))]

    ## Define what type of expression we are looking at
    if(type == 'cleaned') {
        y <- cleaned[i, ]
        ylab <- 'Norm Expr: disease status and project removed'
    } else if (type == 'norm') {
        y <- log2(rpkm[i, ] + 0.5)
        ylab <- 'Normalized Expression'
    }
    x <- colData(rse_gene)$pmilvl
    ylim <- abs(range(y)) * c(0.95, 1.05) * sign(range(y))
```

```
    ## Subset to a given study
    if(!missing(study)) {
        y <- y[rse_gene$project == study]
        x <- x[rse_gene$project == study]
    }

    ## Plot components
    f <- lm(y ~ as.integer(x))
    title <- paste(symb, 'p-value', signif(summary(f)$coef[2, 4], 3))
    if(!missing(study)) title <- paste(title, 'study', study)

    ## Make the plot ^_^
    plot(y ~ x, xlab = 'PMI interval (h)', ylab = ylab, outline = FALSE,
        ylim = ylim, main = title)
    ## Colors from color blind friendly palette at
    ## http://www.cookbook-r.com/Graphs/Colors_(ggplot2)/
    points(y ~ jitter(as.integer(x), 0.6),
        bg = c("#009E73", "#0072B2", "#E69F00", "#D55E00")[
            as.integer(x)], pch = 21)
    abline(f, lwd = 3, col = "#CC79A7")
}
```

In Figures 5, 6, 7 and 8 we check if there's a relationship in the gene expression of *RNASE2*, *EGR3*, *HBA1* and *CXCL2* with the 4 PMI intervals (in hours) that are present in our data. The directions of the trends match those in Figure 2d of the original study across the same 4 PMI intervals. In particular, *CXCL2* initially has a higher expression in artery (tibial) for I1, but it tends to increase across intervals I2 to I4 similar to what we observe in Figure 8. In Figure 5 we see a significant decrease ($p < 0.05$) in *RNASE2* expression as the PMI interval increases.

```
## Now make the plots
plot_gene('RNASE2')
```

```
plot_gene('EGR3')
```

```
plot_gene('HBA1')
```

```
plot_gene('CXCL2')
```

Ferreira *et al.* (Ferreira, Muñoz-Aguirre, Reverter, Godinho, et al., 2018) found the same negative trend in *RNASE2* expression through PMI interval progression for 13 tissues that they show in their Figure 2d, but not in the brain. From Figure 2 we know that not all the studies we are using span all PMI intervals. Using those studies that span at least 3 PMI intervals we explore the relationship between *RNASE2* expression and PMI interval progression as shown in Figures 9, 10, 11 and 12. Three out of the four studies show a negative trend although it's not significant in any of them. SRP019762 shows an increase in *RNASE2* expression but it also has no PMIs in the second interval and only two in the fifth interval.

```
## Plot RNASE2 by study
study_nint <- with(colData(rse_gene),
    tapply(pmilvl, project, function(x) length(unique(x))))
```

##### Using recount_brain to explore relationships with post-mortem interval

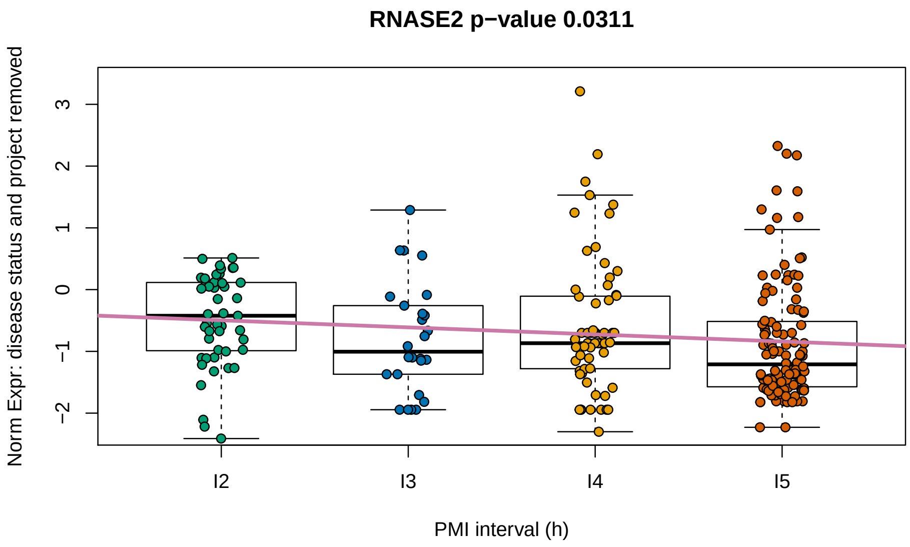

**Figure 5:** **Normalized expression adjusted for disease status and project across PMI intervals for gene RNASE2**

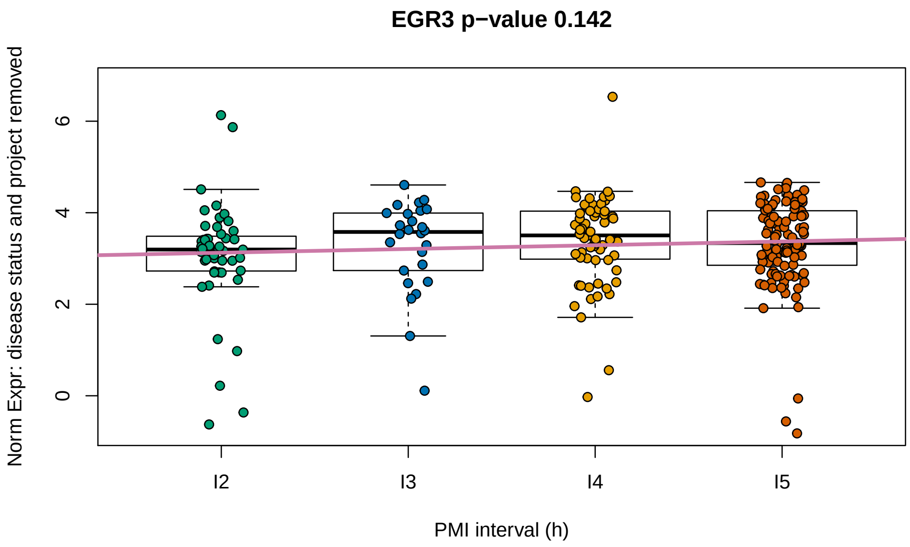

**Figure 6:** **Normalized expression adjusted for disease status and project across PMI intervals for gene EGR3**

```
study_nint <- names(study_nint)[study_nint >= 3]
```

```
for(s in study_nint) plot_gene('RNASE2', type = 'norm', study = s)
```

The variability across studies shown in Figures 9, 10, 11 and 12 as well as the variability in brain regions these studies represent could explain why Ferreira *et al.* didn't see a significant relationship between *RNASE2* expression and PMI interval in their study.

```
with(colData(rse_gene[, rse_gene$project %in% study_nint]),
    table(tissue_site_1, project, useNA = 'ifany'))
```

#### Using recount_brain to explore relationships with post-mortem interval

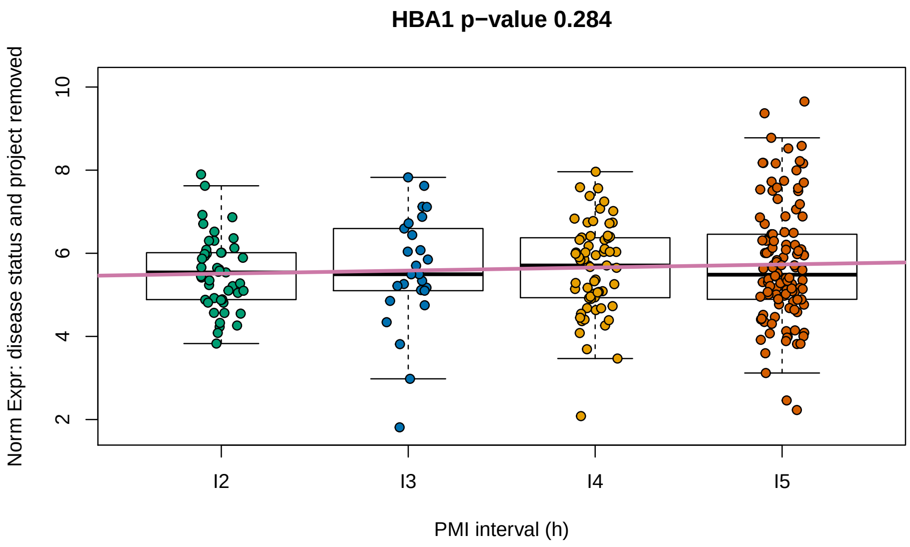

**Figure 7:** **Normalized expression adjusted for disease status and project across PMI intervals for gene HBA1**

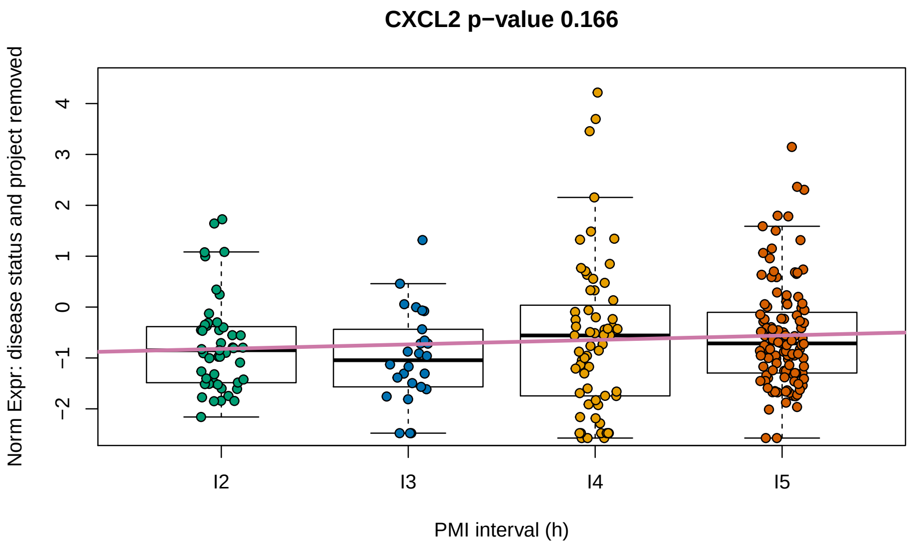

**Figure 8:** **Normalized expression adjusted for disease status and project across PMI intervals for gene CXCL2**

```
##                    project
## tissue_site_1       SRP019762 SRP048683 SRP051844 SRP058181
##   Cerebral cortex          14         0        68        73
##   Corpus callosum           0        12         0         0
with(colData(rse_gene[, rse_gene$project %in% study_nint]),
    table(tissue_site_3, project, useNA = 'ifany'))
##                                   project
## tissue_site_3                       SRP019762 SRP048683 SRP051844 SRP058181
##   Anterior prefrontal cortex               14         0         0         0
##   Dorsolateral prefrontal cortex            0         0        68         0
```

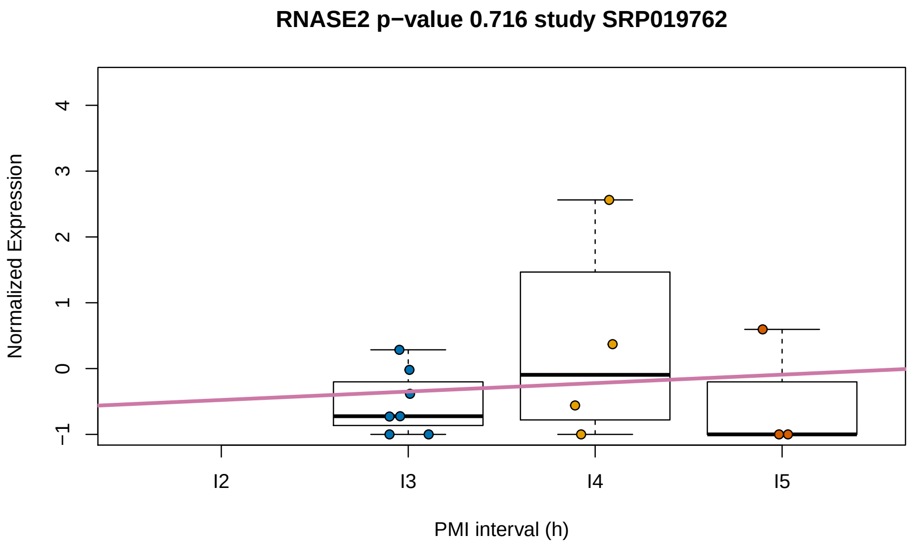

**Figure 9:** Normalized RNASE2 expression for study SRP019762

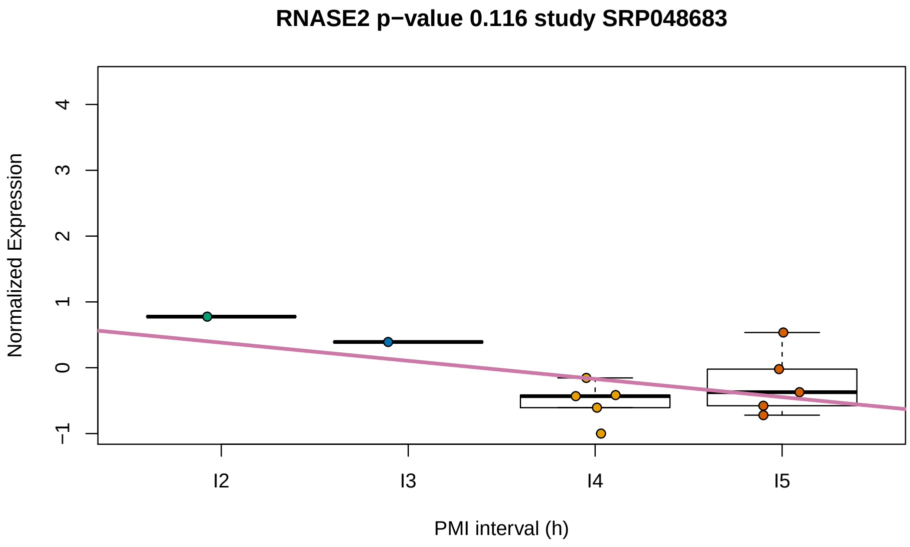

**Figure 10:** Normalized RNASE2 expression for study SRP048683

```
##    Prefrontal cortex                    0         0         0        73
##    <NA>                                 0        12         0         0
```

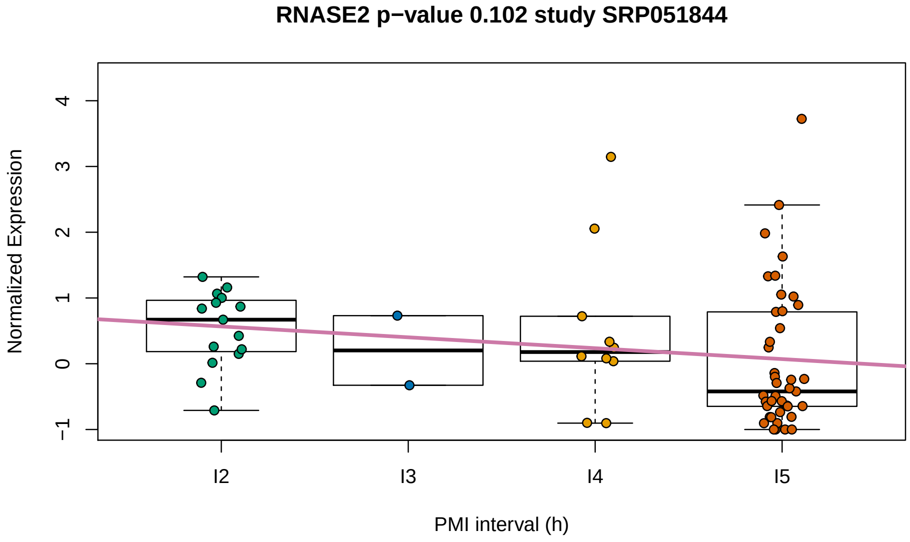

**Figure 11:** Normalized RNASE2 expression for study SRP051844

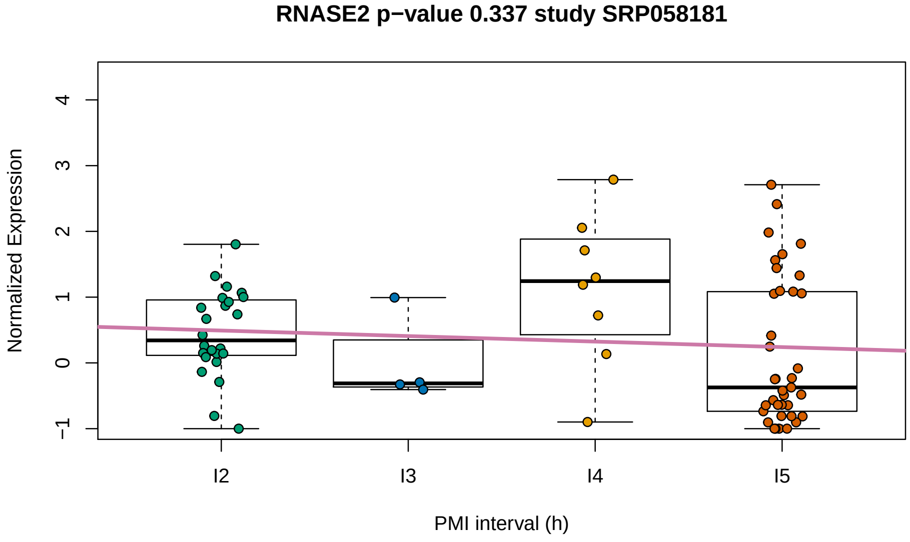

**Figure 12:** Normalized RNASE2 expression for study SRP058181

### 4 Mitochondrial expression and PMI

In Figure 4b and 4c Ferreira *et al.* (Ferreira, Muñoz-Aguirre, Reverter, Godinho, et al., 2018) show how the percent of mitochondrial transcription increases as PMI increases for liver, but decreases for the ovary tissue. In their Supplementary Figure 17 they show that in the brain (cerebellum and cortex) mitochondrial expression and PMI as positively correlated.

One of the advantages of `recount2` is that it provides expression data summarized at different feature types (genes, exons, junctions) and that it provides base-pair resolution coverage BigWig files (Collado-Torres, Nellore, Kammers, Ellis, et al., 2017; Collado-Torres, Nellore, and Jaffe, 2017). Meaning that we can quantify the levels of mitochondrial expression in the same samples and studies we have been working on.

#### 4.1 Quantify chrM expression

As explained in detail in the recount workflow paper (Collado-Torres, Nellore, and Jaffe, 2017), one of the variables included in `recount2` is the *AUC* (area under coverage) which is the sum of the base-pair coverage across all the genome. With the `coverage_matrix()` function we can compute the *AUC* for a given set of region coordinates. So using the chromosome size data for hg38 we can define a single region spanning the whole mitochondrial chromosome. Then we can quantify the `chrM` *AUC* for all of our 252 samples. This will take about 10 to 25 minutes depending on your internet connection.

```
## Got chrM size from
## https://github.com/nellore/runs/blob/master/gtex/hg38.sizes
mt <- GRanges('chrM', IRanges(1, 16569))

## Compute the chrM auc
## Turn scaling off by default so we can then use
## colData(rse_gene)$auc
rse_chrM <- lapply(studies, function(study) {
    message(paste(Sys.time(), 'processing study', study))
    coverage_matrix(project = study, chr = 'chrM', regions = mt,
        chrlen = 16569, scale = FALSE,
        verboseLoad = FALSE, verbose = FALSE)
})
## 2018-03-03 10:12:50 processing study ERP001304
## 2018-03-03 10:15:39 processing study SRP032539
## 2018-03-03 10:17:02 processing study SRP032540
## 2018-03-03 10:18:23 processing study SRP048683
## 2018-03-03 10:19:51 processing study SRP051844
## 2018-03-03 10:29:27 processing study SRP007483
## 2018-03-03 10:31:06 processing study SRP056604
## 2018-03-03 10:32:14 processing study SRP058181
## 2018-03-03 10:42:31 processing study SRP017933
## 2018-03-03 10:45:51 processing study SRP019762

## Combine the list of RSE objects into a single RSE
rse_chrM <- do.call(cbind, rse_chrM)

## Match the order of the samples we have in rse_gene
rse_chrM <- rse_chrM[, match(rse_gene$run, rse_chrM$run)]
```

Note that if we had the BigWig files in our computer, we could use the `coverage_matrix_bwtool()` function from the *recount.bwtool* package (Ellis, Collado-Torres, Jaffe, and Leek, 2017). Although 25 minutes is not a terrible time to wait. If you don't want to wait, check the example_results.Rdata file that we create later on and that contains this information.

#### 4.2 Percent of chrM expression

Now we have all the pieces we need for calculating the percent of mitochondrial expression. Lets save that information in our `rse_gene` object.

```
## chrM auc and percent saved in our sample metadata
colData(rse_gene)$chrM_auc <- assays(rse_chrM)$counts[1, ]
colData(rse_gene)$chrM_percent <- colData(rse_gene)$chrM_auc /
    colData(rse_gene)$auc * 100
```

#### 4.3 Percent of reads in genes

The gene data in `recount2` is actually the *AUC* for the exonic sequence of each gene. So we can use this information to compute the percent of expression in genes (based on Gencode v25). In order to compare it against the mitochondrial expression, we exclude the mitochondrial genes.

```
## Exclude chrM genes
colData(rse_gene)$gene_auc <- colSums(assays(rse_gene)$counts[
    as.logical(seqnames(rowRanges(rse_gene)) != 'chrM'), ])
colData(rse_gene)$gene_percent <- colData(rse_gene)$gene_auc /
    colData(rse_gene)$auc * 100

## Check that none go above 100
with(colData(rse_gene), summary(chrM_percent + gene_percent))
##    Min. 1st Qu.  Median    Mean 3rd Qu.    Max.
##   33.92   76.50   92.55   84.32   94.68   98.84
```

#### 4.4 PMI and chrM expression

Now that we wave all the information we need, we can visualize the relationship between percent of mitochondrial transcription and PMI in hours as shown in Figure 13. The trend is positive just like Ferreira *et al.* found in their Supplementary Figure 17 for brain cortex and brain cerebellum. However, compared to their Figure 4b and 4c, we observe a higher range of mitochondrial transcription. The trend we observe is significantly different from 0.

```
## For simplifying the calls later on
d <- as.data.frame(colData(rse_gene))

## Save plot object for the next plot
g <- ggplot(d, aes(y = chrM_percent, x = pmi)) + geom_point() +
    geom_smooth(method = 'lm') + xlab('PMI in hours') +
    ylab('MT%') + theme_light(base_size = 18)
g
```

```
## Is the association significant? yes
summary(with(d, lm(chrM_percent ~ pmi)))
##
## Call:
## lm(formula = chrM_percent ~ pmi)
##
## Residuals:
```

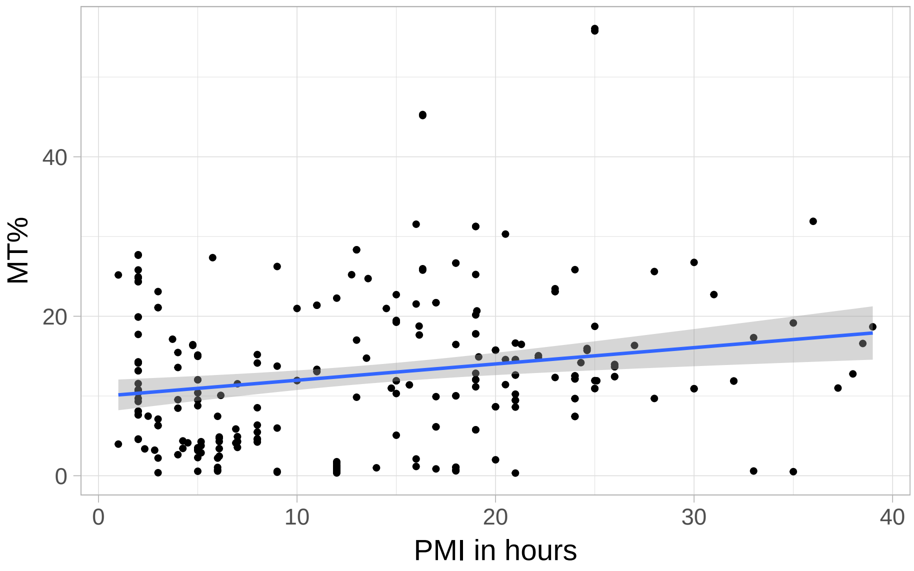

**Figure 13:** PMI in hours is positively associated with mitochondrial transcription

```
##     Min      1Q  Median      3Q     Max
## -16.574  -6.724  -1.587   4.417  41.040
##
## Coefficients:
##             Estimate Std. Error t value Pr(>|t|)
## (Intercept)  9.93532    1.02902   9.655  < 2e-16 ***
## pmi          0.20424    0.06262   3.262  0.00126 **
## ---
## Signif. codes:  0 '***' 0.001 '**' 0.01 '*' 0.05 '.' 0.1 ' ' 1
##
## Residual standard error: 9.264 on 250 degrees of freedom
## Multiple R-squared:  0.04081,    Adjusted R-squared:  0.03698
## F-statistic: 10.64 on 1 and 250 DF,  p-value: 0.001262
```

Figure 14 breaks up samples by disease status. The trend has the same direction although the intercepts do change.

```
g + facet_grid(~ disease_status)
```

In Figure 15 we observe that some studies have a positive and others a negative relationship between PMI in hours and the percent of mitochondrial transcription. Although the standard errors are wide, for study SRP058181 we see that there is an opposite relationship between cases and controls, while it remains negative by disease status for SRP051844.

```
## Make the projects into a factor so that we can use
## scale_colour_hue(drop = FALSE) when plotting a subset
## of the studies and the colors will stay consistent when
## using the full data
d$project <- as.factor(d$project)

## Use only the studies that have PMI values at least in 3 PMI intervals
```

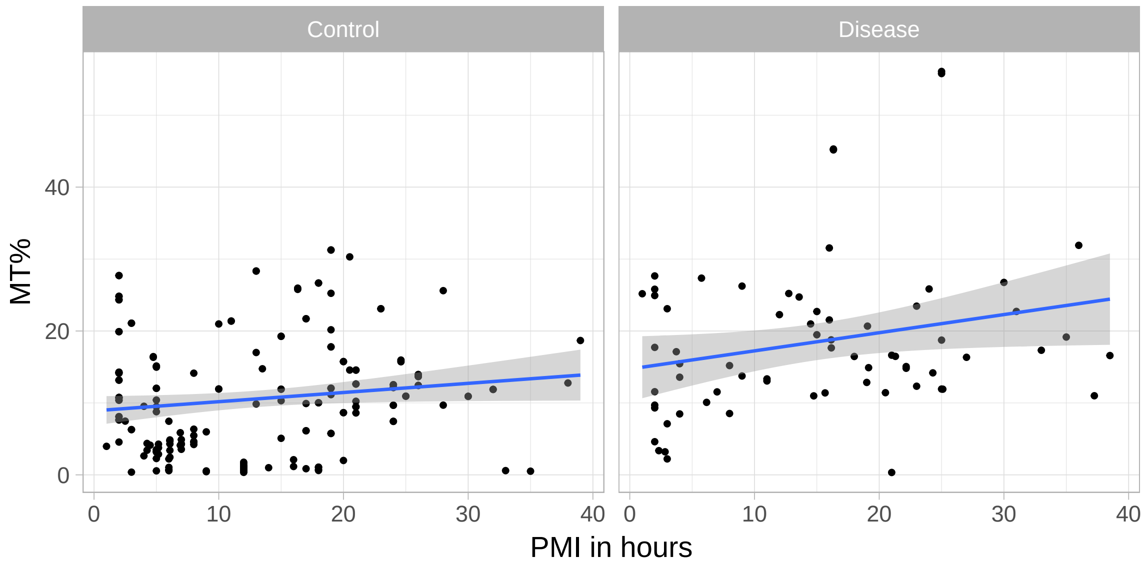

**Figure 14:** **PMI in hours is positively associated with mitochondrial transcription regardless of disease status**

```
ggplot(subset(d, project %in% study_nint),
    aes(y = chrM_percent, x = pmi, colour = project)) +
    geom_point() + geom_smooth(method = 'lm') +
    xlab('PMI in hours') + ylab('MT%') +
    theme_light(base_size = 18) +
    scale_colour_hue(l = 40, drop = FALSE, name = 'study') +
    facet_grid(~ disease_status)
```

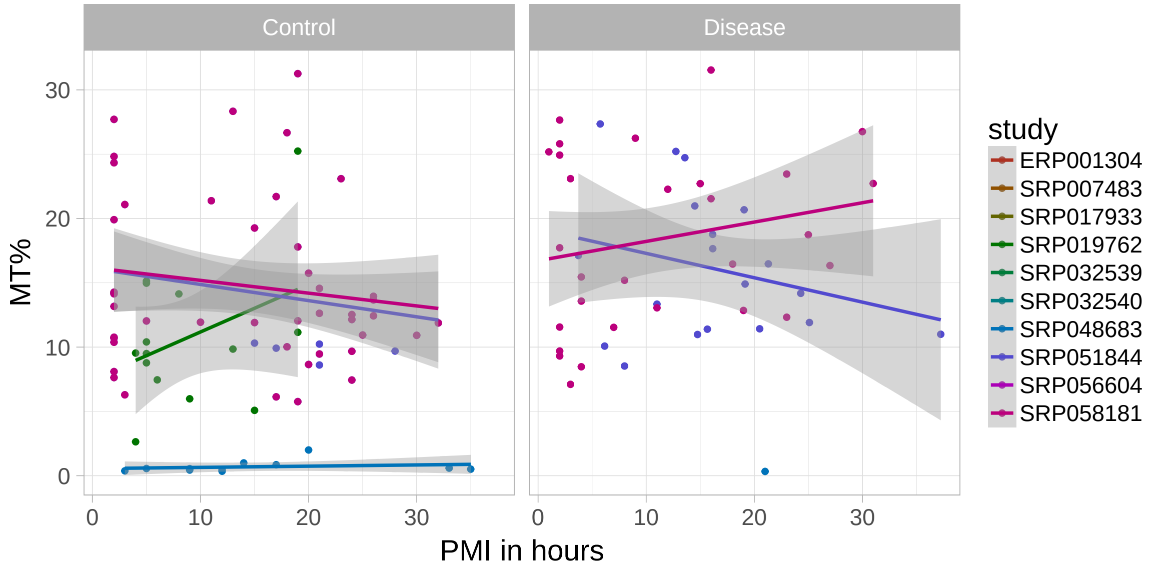

**Figure 15:** **PMI in hours and mitochondrial transcription by study for studies with PMI values across at least 3 intervals**

#### 4.5 PMI and gene expression

Just like we explored the association between PMI in hours and mitochondrial expression in Figure 13, in Figure 16 we check the relationship between PMI in hours and the percent of transcription that overlaps known genes (from Gencode v25) excluding the mitochondrial genes. The global trend is also slightly positive.

```
g2 <- ggplot(d, aes(y = gene_percent, x = pmi)) + geom_point() +
    geom_smooth(method = 'lm') + xlab('PMI in hours') +
    ylab('Gene % (excluding MT)') + theme_light(base_size = 18)
g2
```

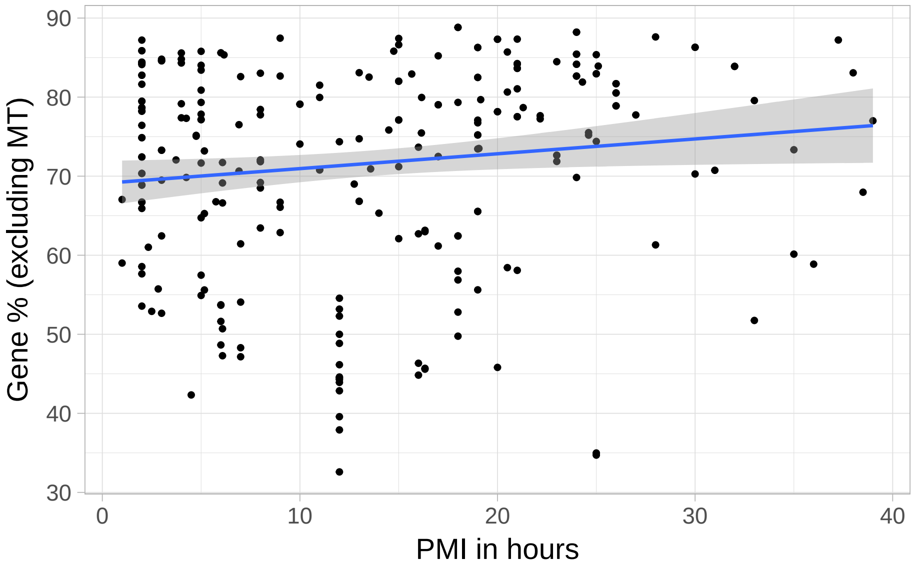

**Figure 16: PMI in hours is positively associated with global gene transcription excluding mitochondrial genes**

Unlike Figure 14, in Figure 17 we do see different trends by disease status. However, the error bands are wide and we can see in a linear regression that neither the disease status term nor the interaction between disease status and PMI in hours are significant.

```
g2 + facet_grid(~ disease_status)
```

```
## Use a linear regresion and check if the
## disease_status coefficient is significantly different from 0
summary(with(d, lm(gene_percent ~ pmi * disease_status)))
##
## Call:
## lm(formula = gene_percent ~ pmi * disease_status)
##
## Residuals:
##     Min      1Q  Median      3Q     Max
## -38.406  -7.323   3.903  10.018  17.658
##
## Coefficients:
```

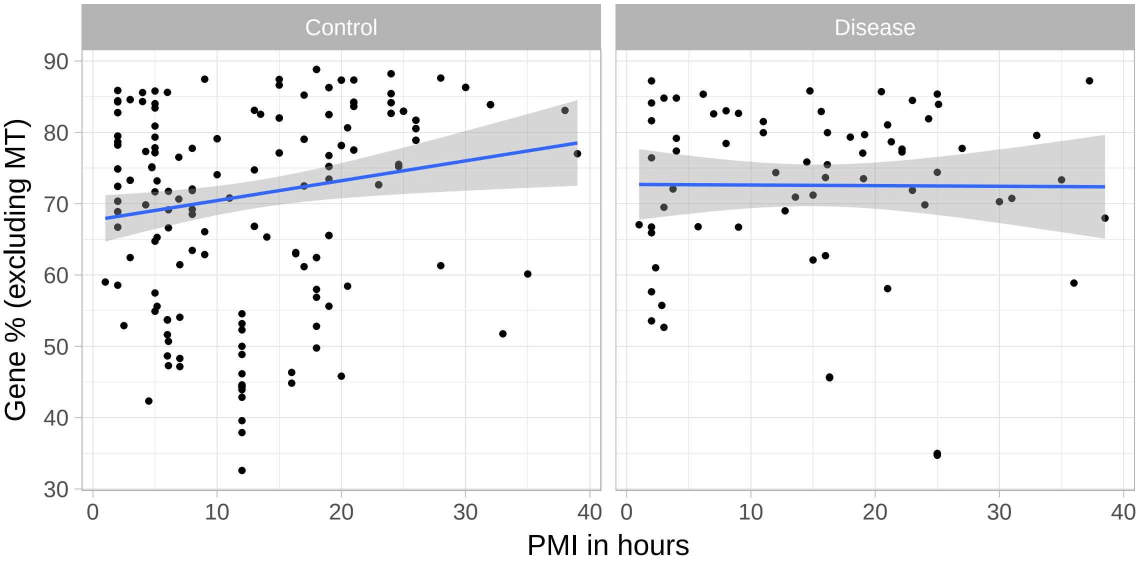

**Figure 17:** **PMI in hours is positively associated with global gene transcription excluding mitochondrial genes for control samples and has no association for the samples wiht a disease phenotype, although they are not significantly different**

```
##                            Estimate Std. Error t value Pr(>|t|)
## (Intercept)                 67.6556     1.6975  39.855   <2e-16 ***
## pmi                          0.2783     0.1079   2.578   0.0105 *
## disease_statusDisease        5.0473     3.2706   1.543   0.1240
## pmi:disease_statusDisease   -0.2871     0.1877  -1.530   0.1274
## ---
## Signif. codes:  0 '***' 0.001 '**' 0.01 '*' 0.05 '.' 0.1 ' ' 1
##
## Residual standard error: 13 on 248 degrees of freedom
## Multiple R-squared:  0.02802,    Adjusted R-squared:  0.01626
## F-statistic: 2.383 on 3 and 248 DF,  p-value: 0.06992

## Could also do the following if we assume parallel lines
## Results in p-value > 0.05 for the disease status term
# summary(with(d, lm(gene_percent ~ pmi + disease_status)))
```

Figure 18 is the equivalent of Figure 15 where we again see variability between studies with wide standard errors.

```
ggplot(subset(d, project %in% study_nint),
    aes(y = gene_percent, x = pmi, colour = project)) +
    geom_point() + geom_smooth(method = 'lm') +
    xlab('PMI in hours') + ylab('Gene % (excluding MT)') +
    theme_light(base_size = 18) +
    scale_colour_hue(l = 40, drop = FALSE, name = 'study') +
    facet_grid(~ disease_status)
```

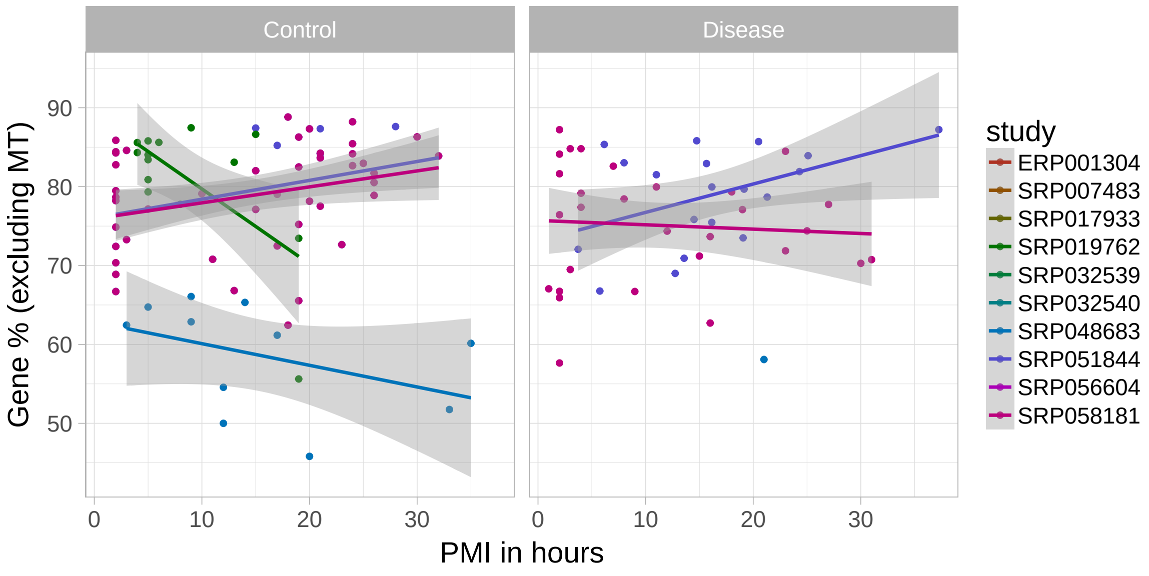

**Figure 18: PMI in hours and global gene transcription excluding mitochondrial genes by study for studies with PMI values across at least 3 intervals**

#### 4.6 chrM vs gene expression

Lets now compare the mitochondrial transcription vs the global gene transcription (excluding mitochondrial genes) percents directly. We expect that high levels of mitochondrial transcription will lead to lower levers of global gene expression. Figure 19 shows that the expected negative association is true, but not for all samples. There is either an elbow curve, or two sets of relationships: one where it's first increasing, and another one that is decreasing as we expected.

```
ggplot(d, aes(y = gene_percent, x = chrM_percent)) +
    geom_point() + ylab('Gene % (excluding MT)') +
    xlab('MT%') + theme_light(base_size = 18) +
    geom_smooth(method = 'loess')
```

Figure 20 shows that PMI interval doesn't seem to play a role in discriminating between the two groups of samples.

```
ggplot(d, aes(y = gene_percent, x = chrM_percent, colour = pmilvl)) +
    geom_point() + ylab('Gene % (excluding MT)') +
    xlab('MT%') + theme_light(base_size = 18) +
    scale_colour_brewer(palette = 'PuOr', name = 'PMI interval')
```

In Figure 21 we separate the samples by disease status and study. Most of the samples from a disease phenotype have the expected negative relationship. However, the disease status doesn't really discriminate between the samples in projects SRP056604 and SRP048683.

```
ggplot(d, aes(y = gene_percent, x = chrM_percent, colour = project)) +
    geom_point() + facet_grid(~disease_status) +
    ylab('Gene % (excluding MT)') + xlab('MT%') +
    theme_light(base_size = 18) +
    scale_colour_hue(l = 40, name = 'study')
```

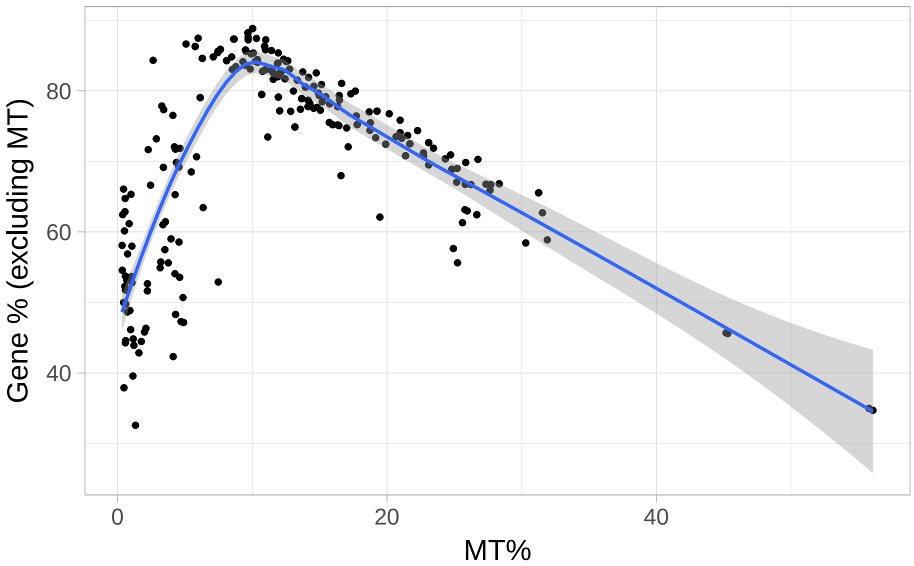

**Figure 19: Mitochondrial expression versus global gene expression excluding mitochondrial genes has an elbow shape or two types of relationships (one increasing, one decreasing)**

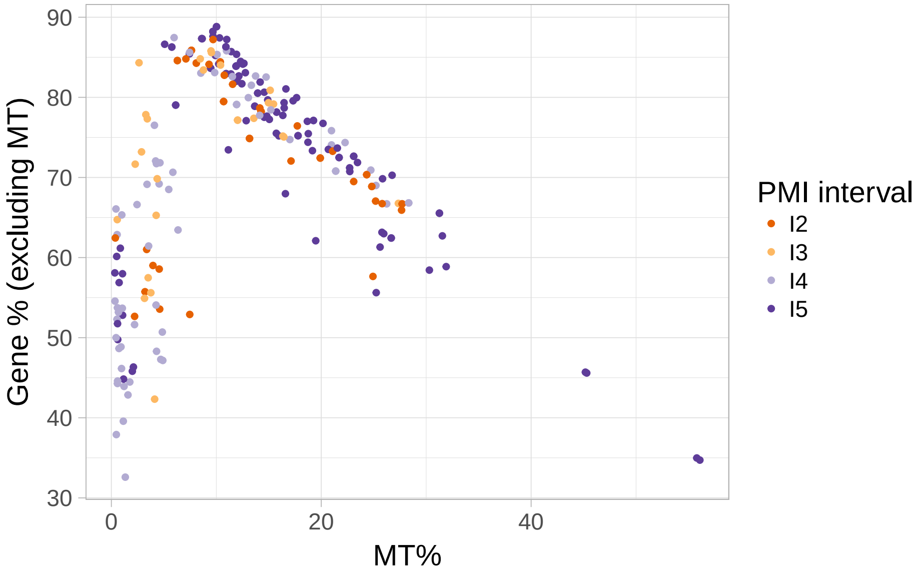

**Figure 20: Mitochondrial expression versus global gene expression excluding mitochondrial genes is not influenced by PMI interval**

In Figure 21 we observe that the studies do help differentiate the samples. If we group the studies in the following two sets:

- set1: SRP017933, SRP032539, SRP032540, SRP048683, SRP056604
- set2: ERP001304, SRP051844, SRP007483, SRP058181, SRP019762

then the samples are perfectly grouped with the two types of relationships between global gene and mitochondrial expression as shown in Figure 22.

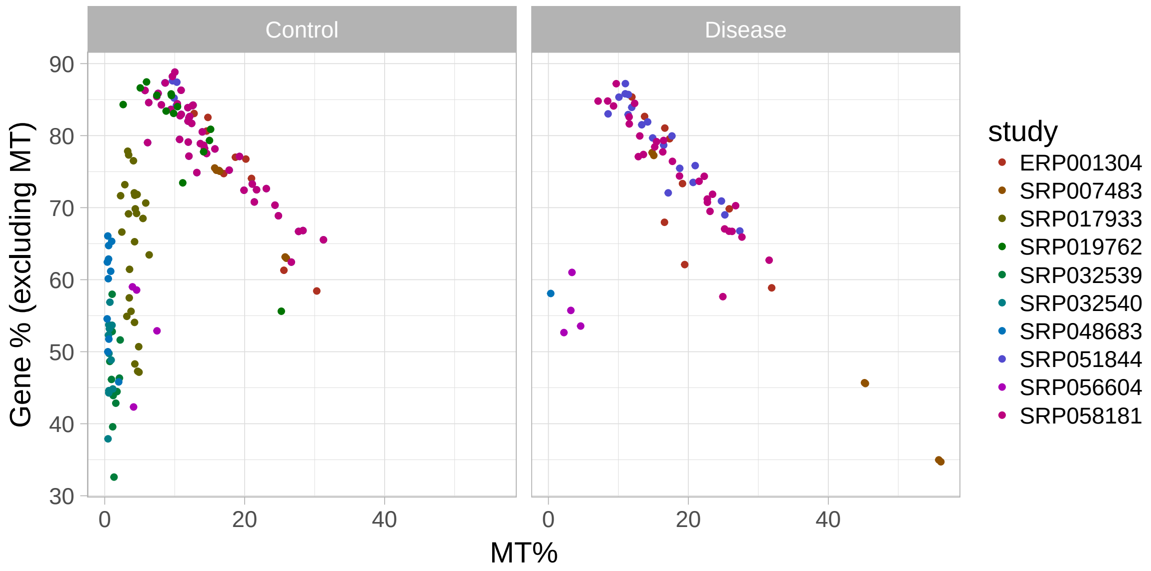

**Figure 21:** **Mitochondrial expression versus global gene expression excluding mitochondrial genes depends on study rather than disease status**

```
## Group studies in two sets
set1 <- c('SRP017933', 'SRP032539', 'SRP032540', 'SRP048683', 'SRP056604')
colData(rse_gene)$study_set <- ifelse(
    colData(rse_gene)$project %in% set1, 'set1', 'set2')

## Remake the d data.frame
d <- as.data.frame(colData(rse_gene))
d$project <- as.factor(d$project)

## Now make the plot
ggplot(d, aes(y = gene_percent, x = chrM_percent,
    colour = study_set)) + geom_point() +
    ylab('Gene % (excluding MT)') + xlab('MT%') +
    theme_light(base_size = 18) +
    geom_smooth(method = 'lm') +
    scale_colour_hue(l = 30, name = 'Study set')
```

Lets revise the relationship between PMI and mitochondrial expression from Figure 13 by separating the studies as shown in Figure 23. For the studies from the first set we see a range of mitochondrial transcription much closer than the one shown by Ferreira *et al.* (Ferreira, Muñoz-Aguirre, Reverter, Godinho, et al., 2018) for liver and ovary tissues in their Figure 4b and 4c. The relationship is negative, contrary to their findings in Supplementary Figure 17 for the brain cerebellum and cortex. Studies from the second set are more in line with our previous observations from Figure 13. Interestingly, there is no significant interaction between PMI in hours and the study sets for explaining the percent of mitochondrial transcription. That is, the main difference is in the intercepts.

```
ggplot(d, aes(y = chrM_percent, x = pmi)) + geom_point() +
    geom_smooth(method = 'lm') + xlab('PMI in hours') +
    ylab('MT%') + theme_light(base_size = 18) +
```

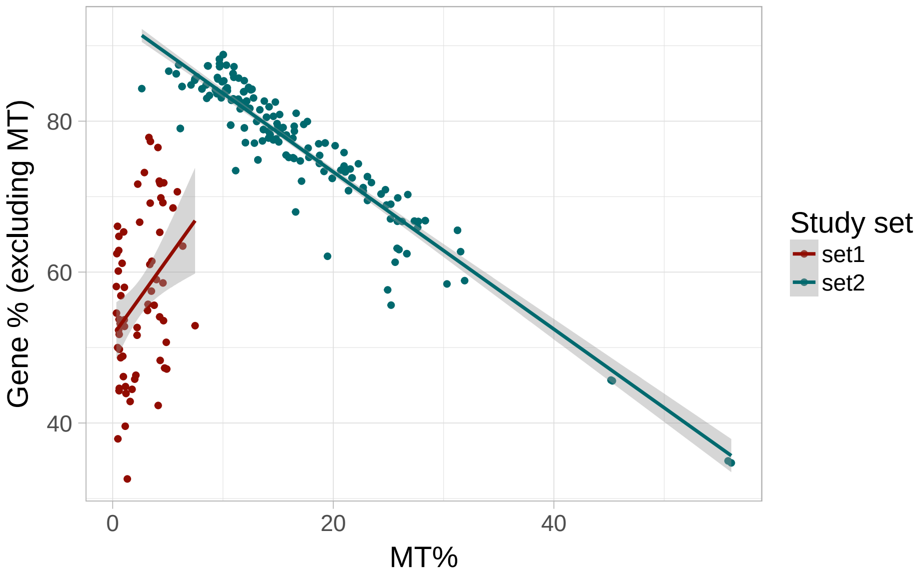

**Figure 22:** Mitochondrial expression versus global gene expression excluding mitochondrial genes depends on study

```
    facet_grid(~ study_set)
```

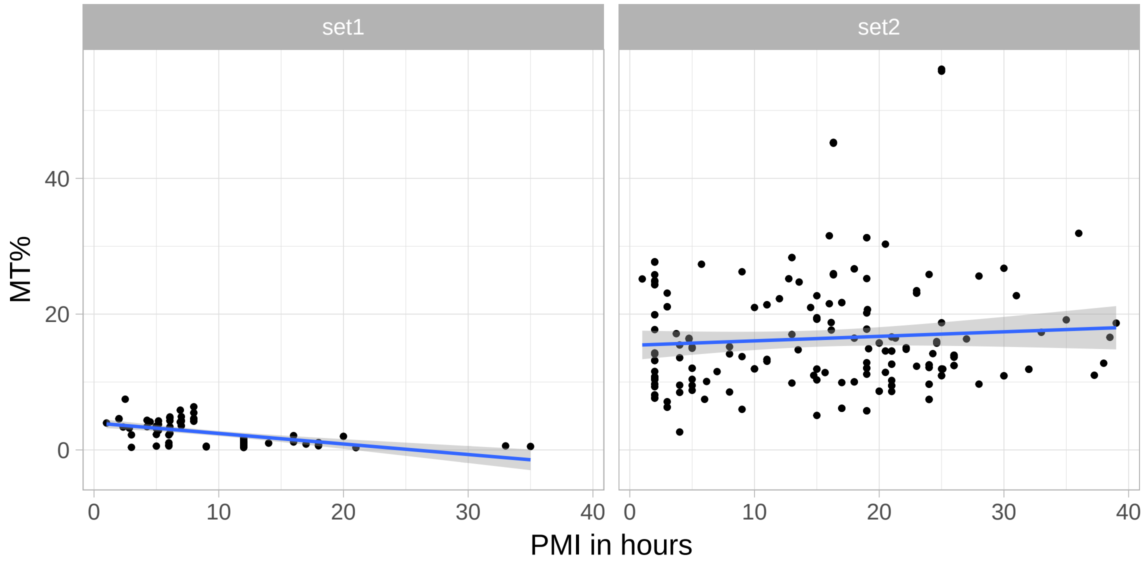

**Figure 23:** PMI in hours is positively associated with mitochondrial transcription for studies from set2

```
## There is a significant difference by study set
summary(with(d, lm(chrM_percent ~ pmi + study_set)))
##
## Call:
## lm(formula = chrM_percent ~ pmi + study_set)
##
```

```
## Residuals:
##     Min      1Q  Median      3Q     Max
## -13.358  -4.495  -1.595   2.347  39.324
##
## Coefficients:
##               Estimate Std. Error t value Pr(>|t|)
## (Intercept)    2.15351    0.99974   2.154   0.0322 *
## pmi            0.03630    0.05026   0.722   0.4709
## study_setset2 13.69631    1.06029  12.918   <2e-16 ***
## ---
## Signif. codes:  0 '***' 0.001 '**' 0.01 '*' 0.05 '.' 0.1 ' ' 1
##
## Residual standard error: 7.183 on 249 degrees of freedom
## Multiple R-squared:  0.4257, Adjusted R-squared:  0.4211
## F-statistic: 92.28 on 2 and 249 DF,  p-value: < 2.2e-16

## The intercepts are significantly different,
## but the interaction term isn't
summary(with(d, lm(chrM_percent ~ pmi * study_set)))
##
## Call:
## lm(formula = chrM_percent ~ pmi * study_set)
##
## Residuals:
##     Min      1Q  Median      3Q     Max
## -13.024  -4.698  -0.954   1.975  39.020
##
## Coefficients:
##                   Estimate Std. Error t value Pr(>|t|)
## (Intercept)         3.9784     1.5605   2.549   0.0114 *
## pmi                -0.1552     0.1356  -1.145   0.2534
## study_setset2      11.4170     1.8348   6.223 2.07e-09 ***
## pmi:study_setset2   0.2218     0.1459   1.520   0.1297
## ---
## Signif. codes:  0 '***' 0.001 '**' 0.01 '*' 0.05 '.' 0.1 ' ' 1
##
## Residual standard error: 7.164 on 248 degrees of freedom
## Multiple R-squared:  0.431,  Adjusted R-squared:  0.4241
## F-statistic: 62.61 on 3 and 248 DF,  p-value: < 2.2e-16
```

Note that the PMI term, that is the association between PMI measured in hours and mitochondrial expression, is not significant. This is in contrast to our initial observation in Figure 13.

#### 4.7 Differences between sets of studies

These two sets of studies might have some properties that differentiate them. In this section we explore in many different ways the differences between the study sets.

As we saw in Figure 21, the first set is mostly made up of control samples. Also, the PMI intervals don't seem to play a role just like we observed in Figure 20.

```
with(colData(rse_gene), table(disease_status, study_set, useNA = 'ifany'))
##                study_set
## disease_status set1 set2
##        Control   62  122
##        Disease    5   63
with(colData(rse_gene), table(pmilvl, study_set, useNA = 'ifany'))
##       study_set
## pmilvl set1 set2
##     I2    8   39
##     I3   11   15
##     I4   37   26
##     I5   11  105
```

#### 4.7.1 Biological

Next we explore some biological covariates. Samples from the second set are all from the cerebral cortex although some samples from the first set are also from the cerebral cortex. For other variables such as the Brodmann area we don't have information for all the samples.

```
with(colData(rse_gene), table(tissue_site_1, study_set, useNA = 'ifany'))
##                   study_set
## tissue_site_1     set1 set2
##   Caudate            5    0
##   Cerebral cortex   32  185
##   Corpus callosum   12    0
##   Hippocampus       13    0
##   Putamen            5    0
with(colData(rse_gene), table(tissue_site_2, study_set, useNA = 'ifany'))
##                   study_set
## tissue_site_2     set1 set2
##   Cingulate gyrus   12    0
##   Frontal lobe      15  159
##   Temporal lobe      5   26
##   <NA>              35    0
with(colData(rse_gene), table(tissue_site_3, study_set, useNA = 'ifany'))
##                                  study_set
## tissue_site_3                    set1 set2
##   Anterior prefrontal cortex        0   14
##   Dorsolateral prefrontal cortex    0   72
##   Motor cortex                     10    0
##   Prefrontal cortex                 0   73
##   Superior temporal gyrus           0   26
##   <NA>                             57    0
with(colData(rse_gene), table(brodmann_area, study_set, useNA = 'ifany'))
##              study_set
## brodmann_area set1 set2
##             4   10    0
##             9    0  145
```

```
##          10      0   14
##          22      0   20
##          41      0    6
##          <NA>   57    0
with(colData(rse_gene), table(hemisphere, study_set, useNA = 'ifany'))
##           study_set
## hemisphere set1 set2
##       Left    0   18
##       <NA>   67  167
```

##### 4.7.2 Demographics

By demographics, we don't see a difference by sex, race (mostly missing) and age for the samples in the two study sets.

```
with(colData(rse_gene), table(sex, study_set, useNA = 'ifany'))
##          study_set
## sex       set1 set2
##   female     5    2
##   male      37  169
##   <NA>      25   14
with(colData(rse_gene), table(race, study_set, useNA = 'ifany'))
##        study_set
## race    set1 set2
##   Asian    0    5
##   Black    0    4
##   White   12   23
##   <NA>    55  153
with(colData(rse_gene), tapply(age, study_set, summary))
## $set1
##    Min. 1st Qu.  Median    Mean 3rd Qu.    Max.
##    4.00   54.00   68.00   64.66   87.00   97.00
##
## $set2
##    Min. 1st Qu.  Median    Mean 3rd Qu.    Max.
##    8.00   52.00   64.00   62.76   76.00  106.00
with(colData(rse_gene), t.test(age ~ study_set))
##
##  Welch Two Sample t-test
##
## data:  age by study_set
## t = 0.51215, df = 88.236, p-value = 0.6098
## alternative hypothesis: true difference in means is not equal to 0
## 95 percent confidence interval:
##  -5.456603  9.245712
## sample estimates:
## mean in group set1 mean in group set2
##           64.65672           62.76216
```

##### 4.7.3 Quality

32 out of the 67 samples from the first set have RIN values with most of the samples in the second set being observed (167 / 185). There is a significant difference in mean RIN between the sets, with the first set having a lower mean RIN. When we use RIN in the graph comparing the percents of transcription in Figure 24, RIN doesn't seem to be as good at differentiating the samples.

```
with(colData(rse_gene), tapply(rin, study_set, summary))
## $set1
##    Min. 1st Qu.  Median    Mean 3rd Qu.    Max.    NA's
##   5.900   6.500   6.750   6.910   7.125   8.800      35
##
## $set2
##    Min. 1st Qu.  Median    Mean 3rd Qu.    Max.    NA's
##   5.200   7.000   7.600   7.545   8.300   9.700      18
with(colData(rse_gene), t.test(rin ~ study_set))
##
##  Welch Two Sample t-test
##
## data:  rin by study_set
## t = -4.449, df = 50.179, p-value = 4.804e-05
## alternative hypothesis: true difference in means is not equal to 0
## 95 percent confidence interval:
##  -0.9215250 -0.3482954
## sample estimates:
## mean in group set1 mean in group set2
##            6.91000            7.54491

ggplot(d, aes(y = gene_percent, x = chrM_percent,
    colour = rin)) + geom_point() +
    ylab('Gene % (excluding MT)') + xlab('MT%') +
    theme_light(base_size = 18) +
    scale_colour_gradientn(colours = c('darkred', 'orange'), name = 'RIN')
```

##### 4.7.4 Technical

We next compare technical covariates between the studies. Samples from the first and second set are mostly single end and paired end, respectively. However, there's some overlap.

```
with(colData(rse_gene), table(paired_end, study_set, useNA = 'ifany'))
##           study_set
## paired_end set1 set2
##      FALSE   55   44
##      TRUE    12  141
```

Looking at library selection, brain bank (assuming that "NICHD Brain and Tissue Bank" and "NICHD Brain and Tissue Bank for Developmental Disorders" are the same ones), instrument, and date of submission we don't see any striking differences between the sets of studies.

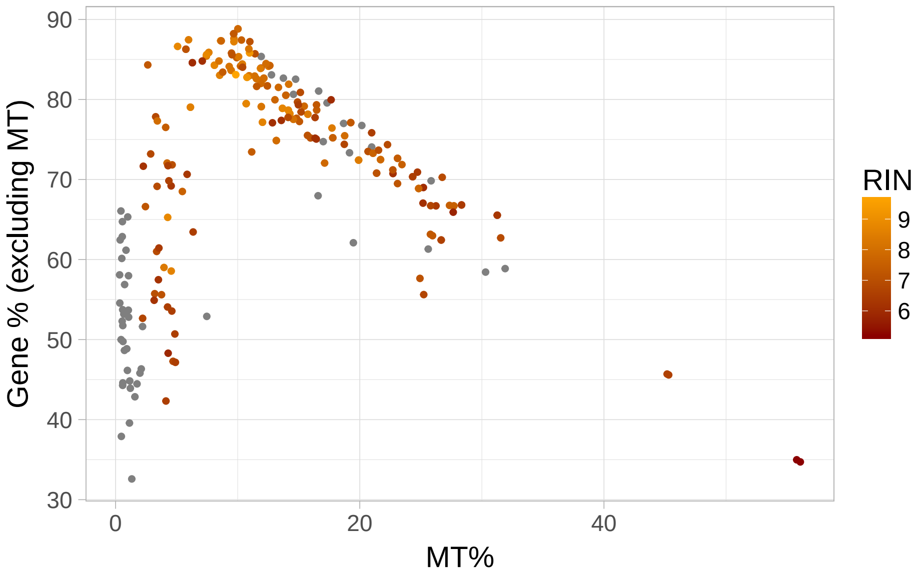

**Figure 24:** Mitochondrial expression versus global gene expression excluding mitochondrial genes with RIN levels

```
## Check technical variables
with(colData(rse_gene), table(libraryselection_s, study_set, useNA = 'ifany'))
##                       study_set
## libraryselection_s   set1 set2
##   CAGE                 25    0
##   cDNA                 31  167
##   RANDOM                0   18
##   size fractionation   11    0
with(colData(rse_gene), table(brain_bank, study_set, useNA = 'ifany'))
##                                                          study_set
## brain_bank                                                set1 set2
##   Autism Tissue Project and Harvard Brain Bank               0   12
##   Brain Resource Center at Johns Hopkins                    22    0
##   Netherlands Brain Bank                                    25    0
##   NICHD Brain and Tissue Bank                               12    0
##   NICHD Brain and Tissue Bank for Developmental Disorders    0    9
##   NSW Tissue Resource Centre                                 0   18
##   Wuhan Xiehe Hospital                                       0    5
##   <NA>                                                       8  141
with(colData(rse_gene), table(instrument_s, study_set, useNA = 'ifany'))
##                                study_set
## instrument_s                    set1 set2
##   Illumina Genome Analyzer II      0   30
##   Illumina Genome Analyzer IIx    25    0
##   Illumina HiSeq 2000             42  155
with(colData(rse_gene),
     table(ss(biosample_submission_date, '-'), study_set, useNA = 'ifany'))
##       study_set
##        set1 set2
```

```
##   2011    0   12
##   2012    0   18
##   2013   47   14
##   2014   12    0
##   2015    8  141
```

Studies from set 1 do have lower median number of reads and mapped reads, however there is no significant difference in the means of these two covariates by study set.

```
with(colData(rse_gene), tapply(reads_downloaded, study_set, summary))
## $set1
##      Min.   1st Qu.    Median      Mean   3rd Qu.      Max.
##    971586   2135760   5065724  88257750 175501444 338362548
##
## $set2
##      Min.   1st Qu.    Median      Mean   3rd Qu.      Max.
##  13172066  39185846  71712108  70365162  86840836 167044880
with(colData(rse_gene), tapply(mapped_read_count, study_set, summary))
## $set1
##      Min.   1st Qu.    Median      Mean   3rd Qu.      Max.
##    385202   1585520   4506226  84196851 164614984 338306029
##
## $set2
##      Min.   1st Qu.    Median      Mean   3rd Qu.      Max.
##  12893433  39029374  71117968  69462444  86133806 163991896

with(colData(rse_gene), t.test(reads_downloaded ~ study_set))
##
##  Welch Two Sample t-test
##
## data:  reads_downloaded by study_set
## t = 1.1947, df = 70.006, p-value = 0.2363
## alternative hypothesis: true difference in means is not equal to 0
## 95 percent confidence interval:
##  -11978565  47763742
## sample estimates:
## mean in group set1 mean in group set2
##           88257750           70365162
with(colData(rse_gene), t.test(mapped_read_count ~ study_set))
##
##  Welch Two Sample t-test
##
## data:  mapped_read_count by study_set
## t = 0.98195, df = 69.85, p-value = 0.3295
## alternative hypothesis: true difference in means is not equal to 0
## 95 percent confidence interval:
##  -15193638  44662452
## sample estimates:
## mean in group set1 mean in group set2
##           84196851           69462444
```

If we look at the overall amount of transcription using the *AUC*, we do see a difference between study sets in both the paired end and the single end samples as shown in Figure 25. However, the study group with the highest *AUC* is not consistent. Note that the *AUC* is roughly the number of mapped reads multiplied by the average read length.

```
d$paired_end_text <- ifelse(d$paired_end, 'Paired end', 'Single end')
ggplot(d, aes(y = auc, x = study_set)) + geom_boxplot() +
    scale_y_log10() +
    facet_grid( ~ paired_end_text) +
    theme_light(base_size = 18) + ylab('AUC') +
    xlab('Study set')
```

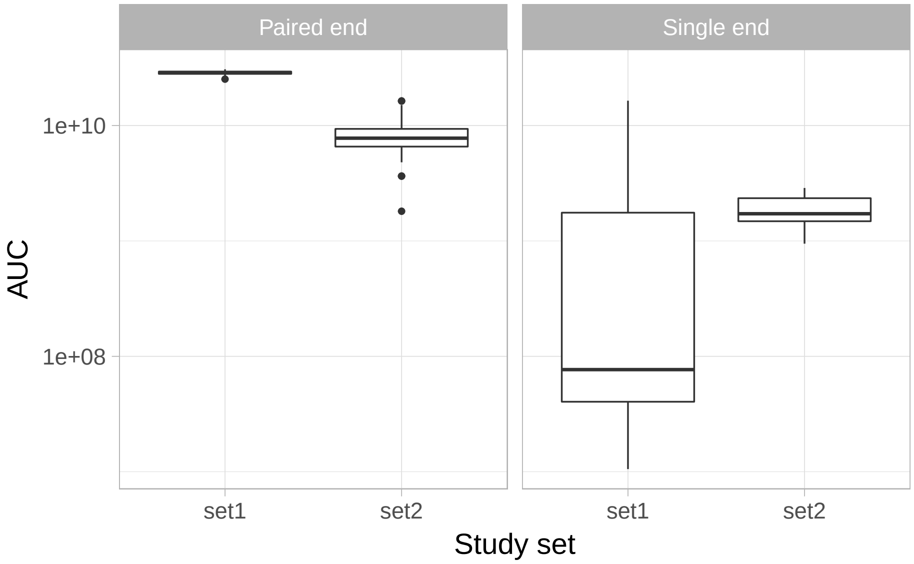

**Figure 25:** **Global expression by AUC between study sets and library protocols**

#### 4.7.5 Read length

In Figure 26 there is a difference in average read length between the studies[2]. But in a closer inspection of the outliers in the second set, we see that there are several samples with shorter read lengths.

```
ggplot(d, aes(y = avgspotlen_l, x = study_set)) + geom_boxplot() +
    theme_light(base_size = 18) + ylab('Average read length') +
    xlab('Study set')
```

```
with(colData(rse_gene), table(avgspotlen_l[study_set == 'set2']))
##
##  73  74  75  76 100 202
##   2   4   5  19  14 141
```

Within the samples from the second set, read length doesn't seem to inform the relationship between mitochondrial and global gene expression as shown in Figure 27.

[2] For paired-end studies, this is the sum of the length of both paired reads.

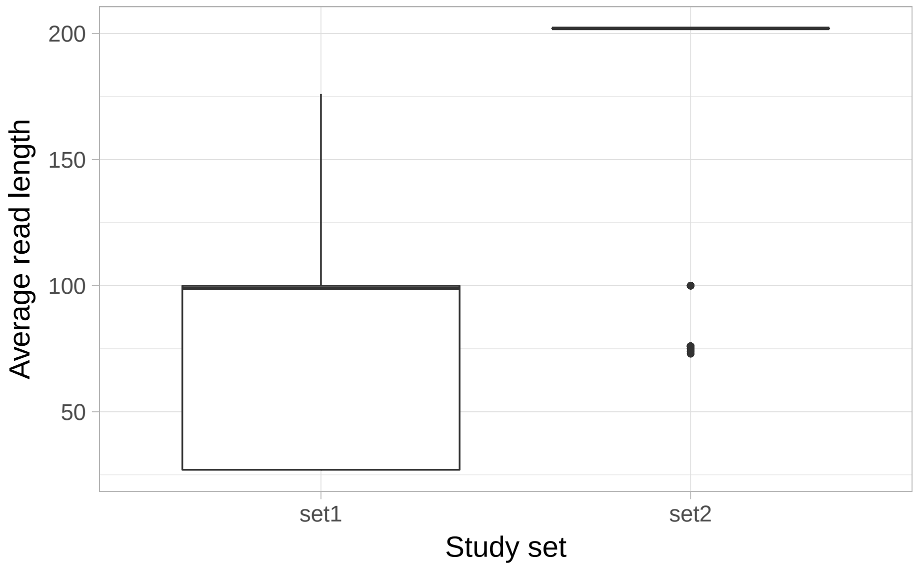

**Figure 26:** Average read length by study group

```
ggplot(d[d$study_set == 'set2', ], aes(y = gene_percent, x = chrM_percent,
    colour = avgspotlen_l < 200)) + geom_point() +
    facet_grid(~study_set) + ylab('Gene % (excluding MT)') +
    xlab('MT%') + theme_light(base_size = 18) +
    scale_colour_brewer(palette = 'Set1', name = 'Avg Read\nLength < 200')
```

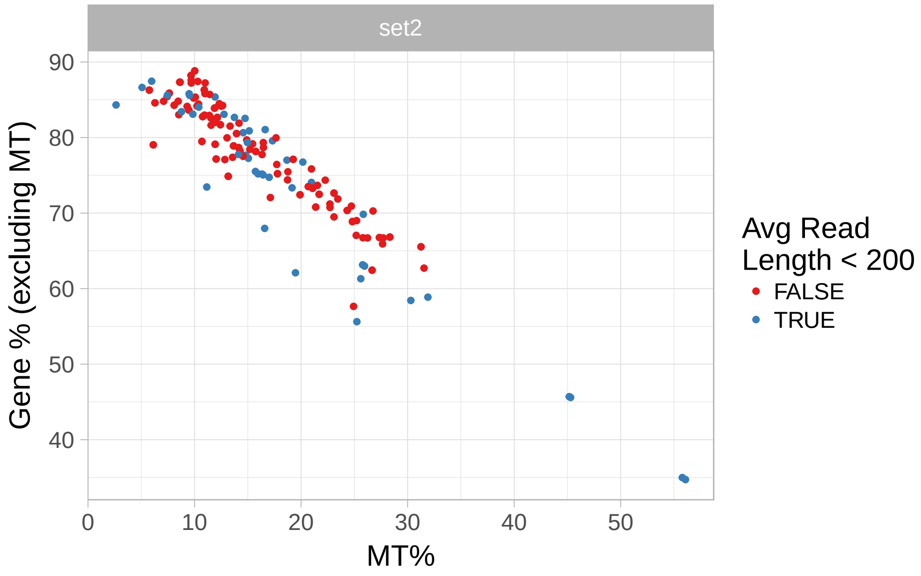

**Figure 27:** Mitochondrial and gene expression for samples from the second set of studies by average read length cut at 200 base-pairs

##### 4.7.6 From predictions

Using the SHARQ beta sample metadata we see that there are cortex samples in both sets, so again, no clear cut separation.

```
## Metadata from SHARQ beta
with(colData(rse_gene), table(sharq_beta_cell_type, study_set, useNA = 'ifany'))
##                       study_set
## sharq_beta_cell_type set1 set2
##               bj        0   12
##               cortex   20   14
##               esc      27    0
##               <NA>     20  159
```

We also have predicted phenotype data (Ellis, Collado-Torres, Jaffe, and Leek, 2017) which we can get using the `add_predictions()` function from *recount*. None of the samples are predicted to be cell lines, and we again see that most samples are from males with 38 / 42 samples matching the reported and the predicted sex for set 1 and all of them matching for set 2.

```
## Get the predictions from phenopredict
rse_gene <- add_predictions(rse_gene)
## 2018-03-03 11:39:18 downloading the predictions to /var/folders/cx/n9s558kx6fb7jf5z_pgszgb80000gn/T//Rtmp(
## Loading objects:
##   PredictedPhenotypes

## Check sample source, and sex
with(colData(rse_gene),
    table(predicted_samplesource, study_set, useNA = 'ifany'))
##                         study_set
## predicted_samplesource set1 set2
##             cell_line     0    0
##             tissue       67  185
##             Unassigned    0    0
with(colData(rse_gene),
    table(predicted_sex, study_set, useNA = 'ifany'))
##                study_set
## predicted_sex set1 set2
##    female       18    7
##    male         49  178
##    Unassigned    0    0
with(colData(rse_gene),
    addmargins(table(predicted_sex, sex, study_set, useNA = 'ifany')))
## , , study_set = set1
##
##                sex
## predicted_sex female male <NA> Sum
##    female          3    2   13  18
##    male            2   35   12  49
##    Unassigned      0    0    0   0
##    Sum             5   37   25  67
##
```

```
## , , study_set = set2
##
##               sex
## predicted_sex female male <NA> Sum
##    female          2    0    5   7
##    male            0  169    9 178
##    Unassigned      0    0    0   0
##    Sum             2  169   14 185
##
## , , study_set = Sum
##
##               sex
## predicted_sex female male <NA> Sum
##    female          5    2   18  25
##    male            2  204   21 227
##    Unassigned      0    0    0   0
##    Sum             7  206   39 252
```

#### 4.8 Save results

We finally save our exploratory results in case we want to carry out more analyses with them later on.

```
## Save results
save(rse_gene, rse_chrM, file = 'example_results.Rdata')
```

### 5 Conclusions

In this document we showed how you can identify samples and studies of interest using `recount_brain` to check if some results from another study are replicated in brain samples from the SRA present in `recount2`. We found that *RNASE2* expression is negatively associated with PMI intervals and show that the variability across studies and brain regions could be why Ferreira *et al.* (Ferreira, Muñoz-Aguirre, Reverter, Godinho, et al., 2018) didn't observe a significant association in their study. We observed a significant increase in mitochondrial transcription with increasing PMI measured in hours that matches the observations by Ferreira *et al.* for brain cortex and brain cerebellum. However, we identified two sets of studies that present different relationships between mitochondrial and global gene transcription (excluding mitochondrial genes). Once we considered the two of studies we found that there is no significant association between mitochondrial transcription and PMI measured in hours. We also showed how you can compare two sets of studies using sample metadata from `recount_brain`, SHARQ beta information present in `recount2` and predicted sample metadata (Ellis, Collado-Torres, Jaffe, and Leek, 2017).

### Reproducibility

```
## Reproducibility information
Sys.time()
## [1] "2018-03-03 11:39:30 EST"
proc.time()
##    user  system elapsed
##  73.341   4.434  91.684
options(width = 120)
devtools::session_info()
## Session info -------------------------------------------------------------------------------------------------
##  setting  value
##  version  R Under development (unstable) (2017-11-29 r73789)
##  system   x86_64, darwin15.6.0
##  ui       X11
##  language (EN)
##  collate  en_US.UTF-8
##  tz       America/New_York
##  date     2018-03-03
## Packages -----------------------------------------------------------------------------------------------------
##  package              * version   date       source
##  acepack                1.4.1     2016-10-29 CRAN (R 3.5.0)
##  AnnotationDbi          1.41.4    2017-12-11 Bioconductor
##  assertthat             0.2.0     2017-04-11 CRAN (R 3.5.0)
##  backports              1.1.2     2017-12-13 CRAN (R 3.5.0)
##  base                 * 3.5.0     2017-11-29 local
##  base64enc              0.1-3     2015-07-28 CRAN (R 3.5.0)
##  bibtex                 0.4.2     2017-06-30 CRAN (R 3.5.0)
##  bindr                  0.1       2016-11-13 CRAN (R 3.5.0)
##  bindrcpp               0.2       2017-06-17 CRAN (R 3.5.0)
##  Biobase              * 2.39.2    2018-01-25 Bioconductor
##  BiocGenerics         * 0.25.3    2018-02-09 Bioconductor
##  BiocParallel         * 1.13.1    2017-12-31 Bioconductor
##  BiocStyle            * 2.7.8     2018-01-20 Bioconductor
##  biomaRt                2.35.12   2018-03-03 Bioconductor
##  Biostrings             2.47.9    2018-02-10 Bioconductor
##  bit                    1.1-12    2014-04-09 CRAN (R 3.5.0)
##  bit64                  0.9-7     2017-05-08 CRAN (R 3.5.0)
##  bitops                 1.0-6     2013-08-17 CRAN (R 3.5.0)
##  blob                   1.1.0     2017-06-17 CRAN (R 3.5.0)
##  bookdown               0.7       2018-02-18 CRAN (R 3.5.0)
##  BSgenome               1.47.5    2018-02-13 Bioconductor
##  bumphunter             1.21.0    2017-10-31 Bioconductor
##  checkmate              1.8.5     2017-10-24 CRAN (R 3.5.0)
##  cluster                2.0.6     2017-03-10 CRAN (R 3.5.0)
##  codetools              0.2-15    2016-10-05 CRAN (R 3.5.0)
##  colorout             * 1.1-3     2017-11-29 Github (jalvesaq/colorout@e2a175c)
##  colorspace             1.3-2     2016-12-14 CRAN (R 3.5.0)
##  compiler               3.5.0     2017-11-29 local
##  curl                   3.1       2017-12-12 CRAN (R 3.5.0)
```

```
## data.table           1.10.4-3 2017-10-27 CRAN (R 3.5.0)
## datasets           * 3.5.0    2017-11-29 local
## DBI                  0.8      2018-03-02 CRAN (R 3.5.0)
## DelayedArray       * 0.5.22   2018-03-02 Bioconductor
## derfinder            1.13.8   2018-02-24 cran (@1.13.8)
## derfinderHelper      1.13.0   2017-10-31 Bioconductor
## devtools             1.13.5   2018-02-18 CRAN (R 3.5.0)
## digest               0.6.15   2018-01-28 CRAN (R 3.5.0)
## doRNG                1.6.6    2017-04-10 CRAN (R 3.5.0)
## downloader           0.4      2015-07-09 CRAN (R 3.5.0)
## dplyr                0.7.4    2017-09-28 CRAN (R 3.5.0)
## evaluate             0.10.1   2017-06-24 CRAN (R 3.5.0)
## foreach              1.4.4    2017-12-12 CRAN (R 3.5.0)
## foreign              0.8-70   2017-11-28 CRAN (R 3.5.0)
## Formula              1.2-2    2017-07-10 CRAN (R 3.5.0)
## GenomeInfoDb       * 1.15.5   2018-02-04 Bioconductor
## GenomeInfoDbData     1.1.0    2017-12-15 Bioconductor
## GenomicAlignments    1.15.12  2018-02-11 Bioconductor
## GenomicFeatures      1.31.10  2018-02-10 Bioconductor
## GenomicFiles         1.15.2   2018-02-09 Bioconductor
## GenomicRanges      * 1.31.22  2018-02-16 Bioconductor
## GEOquery             2.47.18  2018-03-02 Bioconductor
## ggplot2            * 2.2.1    2016-12-30 CRAN (R 3.5.0)
## glue                 1.2.0    2017-10-29 CRAN (R 3.5.0)
## graphics           * 3.5.0    2017-11-29 local
## grDevices          * 3.5.0    2017-11-29 local
## grid                 3.5.0    2017-11-29 local
## gridExtra            2.3      2017-09-09 CRAN (R 3.5.0)
## gtable               0.2.0    2016-02-26 CRAN (R 3.5.0)
## Hmisc                4.1-1    2018-01-03 CRAN (R 3.5.0)
## hms                  0.4.1    2018-01-24 CRAN (R 3.5.0)
## htmlTable            1.11.2   2018-01-20 CRAN (R 3.5.0)
## htmltools            0.3.6    2017-04-28 CRAN (R 3.5.0)
## htmlwidgets          1.0      2018-01-20 CRAN (R 3.5.0)
## httr                 1.3.1    2017-08-20 CRAN (R 3.5.0)
## IRanges            * 2.13.28  2018-02-24 cran (@2.13.28)
## iterators            1.0.9    2017-12-12 CRAN (R 3.5.0)
## jaffelab           * 0.99.18  2018-02-27 Github (LieberInstitute/jaffelab@a8e6430)
## jsonlite             1.5      2017-06-01 CRAN (R 3.5.0)
## knitcitations      * 1.0.8    2017-07-04 CRAN (R 3.5.0)
## knitr                1.20     2018-02-20 CRAN (R 3.5.0)
## labeling             0.3      2014-08-23 CRAN (R 3.5.0)
## lattice              0.20-35  2017-03-25 CRAN (R 3.5.0)
## latticeExtra         0.6-28   2016-02-09 CRAN (R 3.5.0)
## lazyeval             0.2.1    2017-10-29 CRAN (R 3.5.0)
## limma                3.35.12  2018-02-22 Bioconductor
## locfit               1.5-9.1  2013-04-20 CRAN (R 3.5.0)
## lubridate            1.7.3    2018-02-27 CRAN (R 3.5.0)
## magrittr             1.5      2014-11-22 CRAN (R 3.5.0)
## Matrix               1.2-12   2017-11-20 CRAN (R 3.5.0)
## matrixStats        * 0.53.1   2018-02-11 CRAN (R 3.5.0)
```

```
##  memoise                1.1.0     2017-04-21 CRAN (R 3.5.0)
##  methods              * 3.5.0     2017-11-29 local
##  munsell                0.4.3     2016-02-13 CRAN (R 3.5.0)
##  nnet                   7.3-12    2016-02-02 CRAN (R 3.5.0)
##  parallel             * 3.5.0     2017-11-29 local
##  pillar                 1.2.1     2018-02-27 CRAN (R 3.5.0)
##  pkgconfig              2.0.1     2017-03-21 CRAN (R 3.5.0)
##  pkgmaker               0.22      2014-05-14 CRAN (R 3.5.0)
##  plyr                   1.8.4     2016-06-08 CRAN (R 3.5.0)
##  prettyunits            1.0.2     2015-07-13 CRAN (R 3.5.0)
##  progress               1.1.2     2016-12-14 CRAN (R 3.5.0)
##  purrr                  0.2.4     2017-10-18 CRAN (R 3.5.0)
##  qvalue                 2.11.0    2017-10-31 Bioconductor
##  R6                     2.2.2     2017-06-17 CRAN (R 3.5.0)
##  rafalib              * 1.0.0     2015-08-09 CRAN (R 3.5.0)
##  RColorBrewer           1.1-2     2014-12-07 CRAN (R 3.5.0)
##  Rcpp                   0.12.15   2018-01-20 CRAN (R 3.5.0)
##  RCurl                  1.95-4.10 2018-01-04 CRAN (R 3.5.0)
##  readr                  1.1.1     2017-05-16 CRAN (R 3.5.0)
##  recount              * 1.5.9     2018-03-01 Github (leekgroup/recount@458d4f2)
##  RefManageR             0.14.20   2017-08-17 CRAN (R 3.5.0)
##  registry               0.5       2017-12-03 CRAN (R 3.5.0)
##  rentrez                1.2.0     2018-02-12 CRAN (R 3.5.0)
##  reshape2               1.4.3     2017-12-11 CRAN (R 3.5.0)
##  rlang                  0.2.0     2018-02-20 CRAN (R 3.5.0)
##  rmarkdown              1.9       2018-03-01 CRAN (R 3.5.0)
##  RMySQL                 0.10.14   2018-02-26 CRAN (R 3.5.0)
##  rngtools               1.2.4     2014-03-06 CRAN (R 3.5.0)
##  rpart                  4.1-13    2018-02-23 CRAN (R 3.5.0)
##  rprojroot              1.3-2     2018-01-03 CRAN (R 3.5.0)
##  Rsamtools              1.31.3    2018-02-02 Bioconductor
##  RSQLite                2.0       2017-06-19 CRAN (R 3.5.0)
##  rstudioapi             0.7       2017-09-07 CRAN (R 3.5.0)
##  rtracklayer            1.39.9    2018-02-11 Bioconductor
##  S4Vectors            * 0.17.36   2018-03-03 Bioconductor
##  scales                 0.5.0     2017-08-24 CRAN (R 3.5.0)
##  segmented              0.5-3.0   2017-11-30 CRAN (R 3.5.0)
##  splines                3.5.0     2017-11-29 local
##  stats                * 3.5.0     2017-11-29 local
##  stats4               * 3.5.0     2017-11-29 local
##  stringi                1.1.6     2017-11-17 CRAN (R 3.5.0)
##  stringr                1.3.0     2018-02-19 CRAN (R 3.5.0)
##  SummarizedExperiment * 1.9.15    2018-02-24 cran (@1.9.15)
##  survival               2.41-3    2017-04-04 CRAN (R 3.5.0)
##  tibble                 1.4.2     2018-01-22 CRAN (R 3.5.0)
##  tidyr                  0.8.0     2018-01-29 CRAN (R 3.5.0)
##  tools                  3.5.0     2017-11-29 local
##  utils                * 3.5.0     2017-11-29 local
##  VariantAnnotation      1.25.12   2018-01-25 Bioconductor
##  withr                  2.1.1     2017-12-19 CRAN (R 3.5.0)
##  xfun                   0.1       2018-01-22 CRAN (R 3.5.0)
```

```
## XML                     3.98-1.10 2018-02-19 CRAN (R 3.5.0)
## xml2                    1.2.0     2018-01-24 CRAN (R 3.5.0)
## xtable                  1.8-2     2016-02-05 CRAN (R 3.5.0)
## XVector                 0.19.9    2018-02-28 Bioconductor
## yaml                    2.1.17    2018-02-27 CRAN (R 3.5.0)
## zlibbioc                1.25.0    2017-10-31 Bioconductor
```

### References

The analyses were made possible thanks to:

- R (R Core Team, 2017)
- *BiocStyle* (Oleś, Morgan, and Huber, 2018)
- *devtools* (Wickham, Hester, and Chang, 2018)
- *ggplot2* (Wickham, 2009)
- *jaffelab* (Collado-Torres and Jaffe, 2018)
- *knitcitations* (Boettiger, 2017)
- *knitr* (Xie, 2014)
- *recount* (Collado-Torres, Nellore, Kammers, Ellis, et al., 2017; Collado-Torres, Nellore, and Jaffe, 2017; Ellis, Collado-Torres, Jaffe, and Leek, 2017)
- *rmarkdown* (Allaire, Xie, McPherson, Luraschi, et al., 2018)

Full bibliography file.

[1] J. Allaire, Y. Xie, J. McPherson, J. Luraschi, et al. rmarkdown: Dynamic Documents for R. R package version 1.9. 2018. URL: https://CRAN.R-project.org/package=rmarkdown.

[2] C. Boettiger. knitcitations: Citations for 'Knitr' Markdown Files. R package version 1.0.8. 2017. URL: https://CRAN.R-project.org/package=knitcitations.

[3] L. Collado-Torres and A. E. Jaffe. jaffelab: Commonly used functions by the Jaffe lab. R package version 0.99.18. 2018. URL: https://github.com/LieberInstitute/jaffelab.

[4] L. Collado-Torres, A. Nellore and A. E. Jaffe. "recount workflow: Accessing over 70,000 human RNA-seq samples with Bioconductor [version 1; referees: 1 approved, 2 approved with reservations]". In: F1000Research (2017). DOI: 10.12688/f1000research.12223.1. URL: https://f1000research.com/articles/6-1558/v1.

[5] L. Collado-Torres, A. Nellore, K. Kammers, S. E. Ellis, et al. "Reproducible RNA-seq analysis using recount2". In: Nature Biotechnology (2017). DOI: 10.1038/nbt.3838. URL: http://www.nature.com/nbt/journal/v35/n4/full/nbt.3838.html.

[6] S. E. Ellis, L. Collado-Torres, A. E. Jaffe and J. T. Leek. "Improving the value of public RNA-seq expression data by phenotype prediction". In: bioRxiv (2017). DOI: 10.1101/145656. URL: https://www.biorxiv.org/content/early/2017/06/03/145656.

[7] P. G. Ferreira, M. Muñoz-Aguirre, F. Reverter, C. P. S. Godinho, et al. "The effects of death and post-mortem cold ischemia on human tissue transcriptomes". In: Nature Communications 9.1 (Feb. 2018). DOI: 10.1038/s41467-017-02772-x. URL: https://doi.org/10.1038/s41467-017-02772-x.

[8] A. Oleś, M. Morgan and W. Huber. BiocStyle: Standard styles for vignettes and other Bioconductor documents. R package version 2.7.8. 2018. URL: https://github.com/Bioconductor/BiocStyle.

[9] R Core Team. R: A Language and Environment for Statistical Computing. R Foundation for Statistical Computing. Vienna, Austria, 2017. URL: https://www.R-project.org/.

[10] H. Wickham. ggplot2: Elegant Graphics for Data Analysis. Springer-Verlag New York, 2009. ISBN: 978-0-387-98140-6. URL: http://ggplot2.org.

[11] H. Wickham, J. Hester and W. Chang. devtools: Tools to Make Developing R Packages Easier. R package version 1.13.5. 2018. URL: https://CRAN.R-project.org/package=devtools.

[12] Y. Xie. "knitr: A Comprehensive Tool for Reproducible Research in R". In: Implementing Reproducible Computational Research. Ed. by V. Stodden, F. Leisch and R. D. Peng. ISBN 978-1466561595. Chapman and Hall/CRC, 2014. URL: http://www.crcpress.com/product/isbn/9781466561595.
