## Supplementary File 3 for "*recount-brain*: a curated repository of human brain RNA-seq datasets metadata"

### recount_brain cross-study example

**Shannon E. Ellis** *[1]

[1] Johns Hopkins Bloomberg School of Public Health

*

**19 June 2018**

##### Abstract

Example on how to use recount_brain metadata across multiple studies. We show how to download data from recount2, add the sample metadata from recount_brain, explore the sample metadata and the gene expression data, and perform a cross-study analysis.

### 1 Introduction

This document is an example of how you can use `recount_brain`. We will first explore what samples are available in `recount_brain`. After determining what samples are of most interest to us, we will download the expression data for the studies from which these samples were generated. We will remove batch effects across studies. Finally, we will show how to add the `recount_brain` metadata and perform a gene differential expression analysis using this information to assess replicability across studies. Specifically, we'll be looking to see if the same genes are most variable across different glioblastoma datasets. We'll use a kidney cancer data set as our negative control data set. We'll use concordance across variable genes between datasets as our measure of interest.

#### 1.1 Load libraries

Since we will be using many functions from the `recount` package, let's load it first[1]. We'll also download all of the other

[1] If you are a first time `recount` user, we recommend first reading the package vignette at bioconductor.org/packages/recount.

```
library(recount) # version >1.5.9
library(dplyr)
library(downloader)
library(janitor)
# install_github('LieberInstitute/jaffelab')
library(jaffelab)
```

### 2 The Data

#### 2.1 Sample metadata

To explore the samples available, we'll first search among TCGA samples (McLendon, Friedman, Bigner, Meir, et al., 2008) and available samples in `recount-brain` to determine which samples and which studies we'll use for our analysis. We'll similarly load the TCGA meta data. We will use samples from TCGA and from `recount-brain` in our analysis.

```
## get recount_brain metadata
recount_brain_v1 <- recount::add_metadata(source = 'recount_brain_v1')
## Loading objects:
##   recount_brain
brain = recount_brain_v1

## get TCGA metadata
tcga <- as.data.frame(recount::all_metadata('TCGA'))
```

#### 2.2 Identifying studies

Among the TCGA samples, there are 707 samples with either Glioma or Glioblastoma Multiforme (GBM) (Brennan, Verhaak, McKenna, Campos, et al., 2013). There are four `recount_brain` studies with at least 20 cancer samples. Of these four studies, three studies have Glioblastoma samples. Two of these three studies have more than 20 samples. We'll use these two studies – `SRP027383` ((Bao, Chen, Yang, Zhang, et al., 2014) and `SRP044668` (Gill, Pisapia, Malone, Goldstein, et al., 2014) – to look for variability in expression across studies.

#### 2.3 TCGA brain cancer samples

```
# Find TCGA samples that are brain cancer samples
tcga %>%
  filter(gdc_cases.project.primary_site == "Brain") %>%
  group_by(gdc_cases.project.name) %>%
  summarise (n = n())
## # A tibble: 2 x 2
##   gdc_cases.project.name       n
##   <chr>                    <int>
## 1 Brain Lower Grade Glioma   532
## 2 Glioblastoma Multiforme    175
```

#### 2.4 recount-brain cancer studies

```
# Find recount-brain studies that are cancer studies
(studies <- brain %>% filter(!is.na(tumor_type)) %>%
  group_by(sra_study_s) %>%
  summarise (n = n()) %>%
  filter(n>=20))
## # A tibble: 4 x 2
##   sra_study_s     n
##   <chr>       <int>
## 1 ERP010930      49
## 2 SRP027383     274
## 3 SRP044668      75
## 4 SRP055730      24
```

#### 2.5 Cancer type overlap (TCGA, recount-brain)

```
# Find recount-brain studies with cancer type that overlaps with TCGA
(GBM_samples <- brain %>%
  filter(sra_study_s %in% studies$sra_study_s) %>%
  group_by(sra_study_s, tumor_type) %>%
  summarise (n = n()) %>%
  filter(tumor_type == "Glioblastoma") %>%
```

```
  filter(n>=20))
## # A tibble: 2 x 3
## # Groups:   sra_study_s [2]
##   sra_study_s tumor_type       n
##   <chr>       <chr>        <int>
## 1 SRP027383   Glioblastoma   100
## 2 SRP044668   Glioblastoma    75
```

#### 2.6 Download expression data

Just like any study in `recount2` (Collado-Torres, Nellore, Kammers, Ellis, et al., 2017), we first need to download the gene count data using `recount::download_study()`. We'll additionally use the `add_metadata()` function to include the `recount-brain` metadata information for these two studies. Additionally, below, we read in the TCGA gene expression data from brain samples.

```
# SRA (recount-brain) expression data
todownload <- GBM_samples$sra_study_s

for(i in 1:length(todownload)){
  if(!file.exists(file.path(todownload[i], 'rse_gene.Rdata'))) {
      download_study(todownload[i])
  } else {
      load(file.path(todownload[i], 'rse_gene.Rdata'), verbose = TRUE)
  }
  assign(paste0("rse_gene_", todownload[i]), rse_gene)
}
## Loading objects:
##   rse_gene
## Loading objects:
##   rse_gene

# add recount-brain metadata
rse_gene_SRP027383 <- recount::add_metadata(rse_gene_SRP027383)
## Loading objects:
##   recount_brain
rse_gene_SRP044668 <- recount::add_metadata(rse_gene_SRP044668)
## Loading objects:
##   recount_brain

# TCGA brain expression data
if(!file.exists(file.path('TCGA', 'rse_gene_brain.Rdata'))) {

 dir.create('TCGA', showWarnings = FALSE)
 downloader::download('http://duffel.rail.bio/recount/v2/TCGA/rse_gene_brain.Rdata', destfile =
               'TCGA/rse_gene_brain.Rdata', mode = 'wb')
}

load(file.path('TCGA','rse_gene_brain.Rdata'))
```

```
assign('rse_gene_TCGA', rse_gene)
```

#### 2.7 Expresion Counts

After loading the expression data from our studies of interest, we filter to only include TCGA disease samples, requiring samples to have Glioblastoma (GBM) and to be samples from the primary tumor, rather than nearby healthy tissue or more advanced tissue sample (i.e. a metastasis). We combine all our data into a single object, so that when we scale the data in the next steps, it can all be completed together.

```
# get expression counts
## combine rses to scale counts all together
rse_gene_SRA <- cbind(rse_gene_SRP027383,rse_gene_SRP044668)

## just 15 Primary Tumor GBM samples in TCGA
tokeep <- (colData(rse_gene_TCGA)$gdc_cases.project.name=="Glioblastoma Multiforme" &
             colData(rse_gene_TCGA)$cgc_sample_sample_type=="Primary Tumor")
rse_gene_TCGA<-rse_gene_TCGA[,tokeep]

# get metadata
# add dataset column for easy tracking later
tcga_md <-  as.data.frame(colData(rse_gene_TCGA)) %>%
  mutate(dataset='TCGA', disease_status='Disease', tumor_type="Glioblastoma") %>%
  filter(cgc_sample_sample_type=="Primary Tumor")
SRP027383_md <- as.data.frame(colData(rse_gene_SRP027383)) %>%
  mutate(dataset='SRP027383')
SRP044668_md <- as.data.frame(colData(rse_gene_SRP044668)) %>%
  mutate(dataset='SRP044668')

## make sure that same metadata columns are present so that RSEs can be merged
cols_to_bind <- colnames(colData(rse_gene_TCGA))[colnames(colData(rse_gene_TCGA)) %in%
                                                  colnames(colData(rse_gene_SRA))]
colData(rse_gene_SRA)  <- colData(rse_gene_SRA)[,cols_to_bind]
colData(rse_gene_TCGA)  <- colData(rse_gene_TCGA)[,cols_to_bind]

## merge data so that it can all be scaled together
rse_gene_total <- cbind(rse_gene_TCGA, rse_gene_SRA)

# Expression and metadata combined across data sets
# 538 samples
md <- bind_rows(tcga_md, SRP027383_md) %>%
        bind_rows(., SRP044668_md)
```

### 3 Quality Control

There are a number of critical quality control steps that are necessary before we can make any cross-study comparisons. We'll (1) filter out lowly-expressed genes, (2) remove outlier samples, and (3) normalize the data to remove unwanted sources of variation from the data.

#### 3.1 Filter genes (low expression)

```
## remove lowly expressed genes
rse_rpkm <- getRPKM(scale_counts(rse_gene_total)) # 707 samples

## Compute RPKM and mean RPKM
rpkm_mean <- rowMeans(rse_rpkm)
## Esmate a mean RPKM cutoff
expr_cuts <- expression_cutoff(rse_rpkm)
```

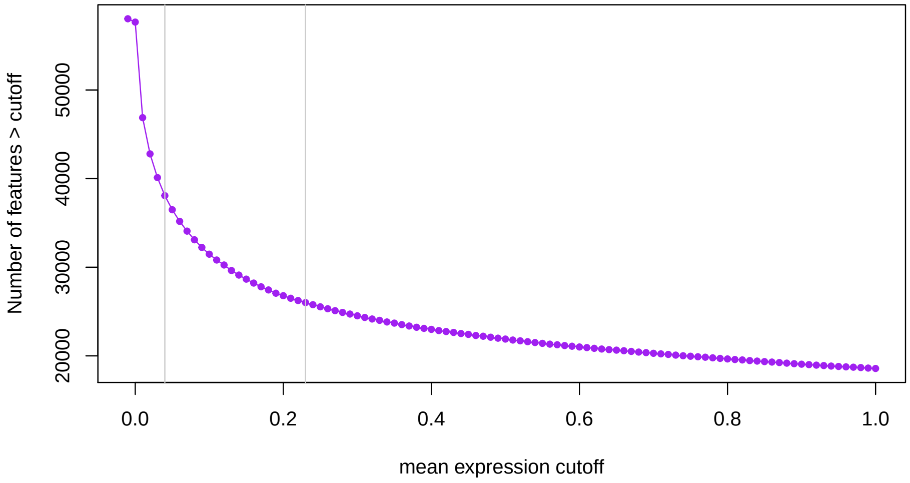

**Figure 1: Number of genes expressed at given mean RPKM cutoff**

```
## 2018-06-19 10:37:38 the suggested expression cutoff is 0.21
```

```
#round(mean(expr_cuts), 2)

## Filter genes with low levels of expression
rpkm <- rse_rpkm[rpkm_mean > round(mean(expr_cuts), 2),]
rpkm_log2 <- log2(rpkm+0.5)
```

Here, we have scale the data and calculate RPKM for all genes in the data set (N = 58037. We then remove lowly expressed genes, defining lowly expressed genes is a mean expression across samples $< 0.21$. This leaves 26499 genes for downstream analysis.

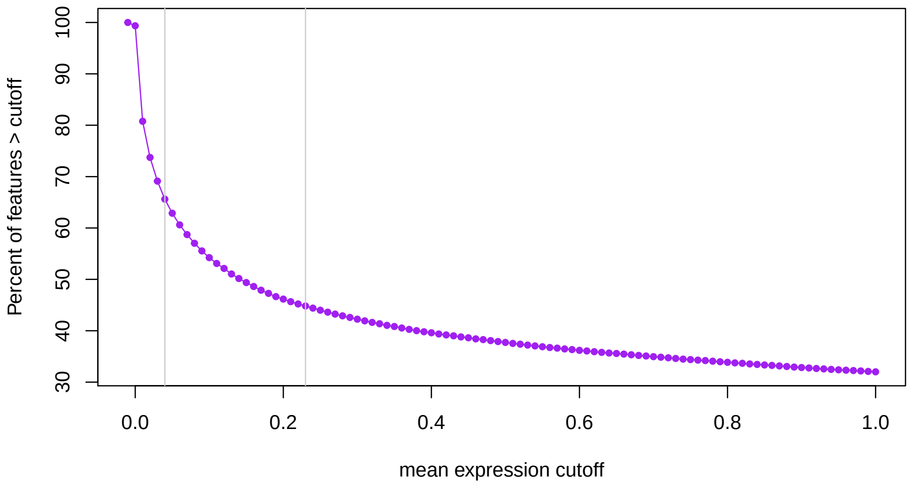

**Figure 2:** Percent of genes epxressed at a given mean RPKM cutoff

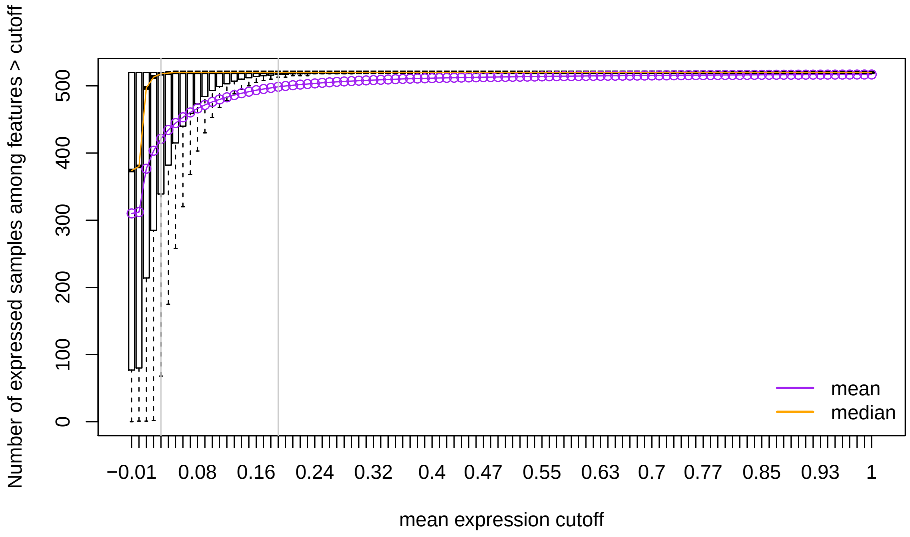

**Figure 3:** Distribution of number of expressed samples across all genes at a given mean RPKM cut-off

#### 3.2 Run PCA

Fist, we'll write a few functions to run PCA, plot the results, and assess the percent of variance explained. These functions will be used throughout.

```
## Run PCA
pc_function <- function(exp){
  svd(exp - rowMeans(exp))
}

## Plot PCA
pc_plot <- function(pca,legend=NULL, color=NULL, main=NULL,
                    type=NULL,
```

```
                    ptsize=1.5,position="bottomright"){
  par(mfrow=c(2,2))

  if(type=='character'){
    color = colors
    c2 = names(table(color))
  }else{
    color = as.factor(color)
    c2 = 1:length(unique(as.factor(color)))
  }

  par(font.lab = 2, cex.lab = 1.2, font.axis = 2, cex.axis = 1.2)
  plot(pca$v[, 1], pca$v[, 2], col= color,
       pch = 19, cex = ptsize,
       xlab = 'PC1', ylab = 'PC2',
       main = main)

  par(font.lab = 2, cex.lab = 1.2, font.axis = 2, cex.axis = 1.2)
  plot(pca$v[, 3], pca$v[, 4], col= color,
       pch = 19, cex = ptsize,
       xlab = 'PC3', ylab = 'PC4',
       main = main)

  par(font.lab = 2, cex.lab = 1.2, font.axis = 2, cex.axis = 1.2)
  plot(pca$v[, 5], pca$v[, 6], col= color ,
       pch = 19, cex = ptsize,
       xlab = 'PC5', ylab = 'PC6',
       main = main)

  par(font.lab = 2, cex.lab = 1.2, font.axis = 2, cex.axis = 1.2)
  plot(pca$v[, 7], pca$v[, 8], col= color,
       pch = 19, cex = ptsize,
       xlab = 'PC7', ylab = 'PC8',
       main = main)
  legend(position, pch = 19, col= c2,
         names(summary(as.factor(legend))),bty="n")
}

## Plot Variance Explained
var_plot <- function(pca){
  par(mfrow=c(1,1))
  plot((pca$d^2/sum(pca$d^2))*100, xlim = c(0, 15), type = "b",
      pch = 16, xlab = "principal components",
      ylab = "variance explained (%)")
}
```

Having filtered out lowly-expressed genes, we now run a principal component analysis (PCA) to look at global expression patterns across this data set. Below we see that, as expected, samples are clustering by study. This is something we will have to correct for before carrying out our analysis.

##### recount_brain cross-study example

```
## Look at mean variance relationship
## Calculate PCs with svd function
expr.pca.rpkm.log2 <- pc_function(rpkm_log2)

## plot by dataset
pc_plot(pca=expr.pca.rpkm.log2,legend=md$dataset, color=md$dataset,
        main="PC (gene level) : log2(RPKM + 0.5)",
        position="bottomleft", type='variable')
```

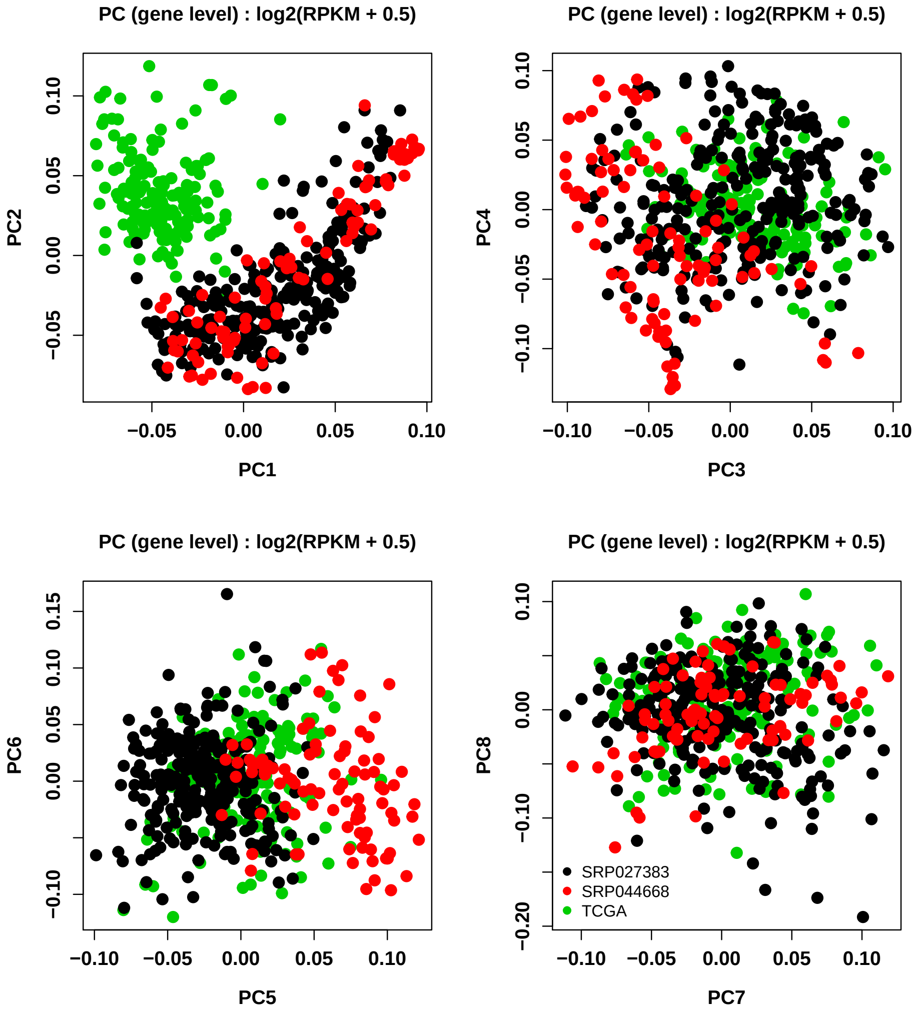

```
## check to see if healthy samples are clustering
pc_plot(pca=expr.pca.rpkm.log2,legend=md$disease_status, color=md$disease_status,
        main="PC (gene level) : log2(RPKM + 0.5)",
        position="bottomleft", type='variable')
```

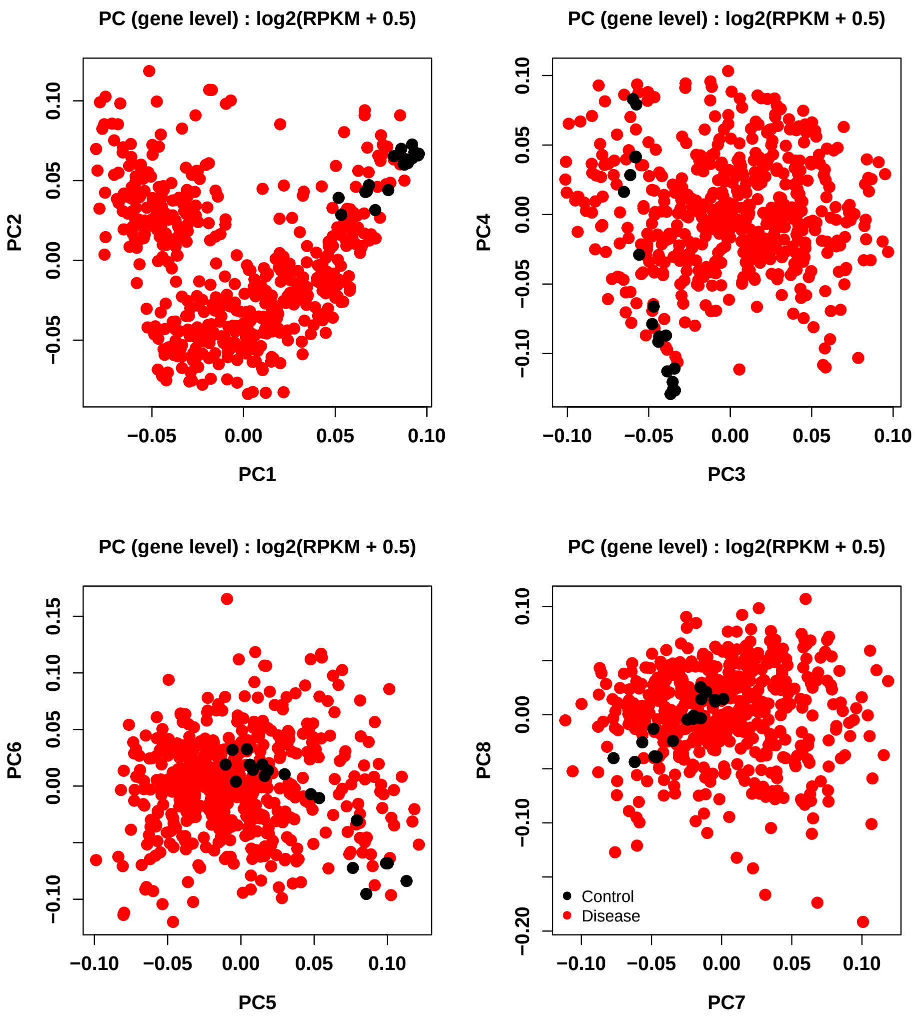

#### 3.3 Remove controls

Given the fact that control samples are clustering in the PCA plot above, we'll remove these from the analysis prior to normalizing the data to remove dataset specific effects. After removing these control samples from analysis, we see that the data still cluster by data set.

```
## remove 19 control samples and just include GBM samples
rpkm_log2 <- rpkm_log2[,(md$tumor_type=="Glioblastoma" & !is.na(md$tumor_type))]
md <- md[(md$tumor_type=="Glioblastoma" & !is.na(md$tumor_type)),]

tabyl(md$dataset)
##  md$dataset   n   percent
##   SRP027383  99 0.3000000
##   SRP044668  74 0.2242424
##        TCGA 157 0.4757576

## Run pca
expr.pca.counts.log2 <- pc_function(rpkm_log2)
```

#### recount_brain cross-study example

```
## visualize
pc_plot(pca=expr.pca.counts.log2, legend=md$dataset,
        color=md$dataset,
        main="PC (gene level) : log2(RPKM + 0.5)",
        position="topleft", type='variable')
```

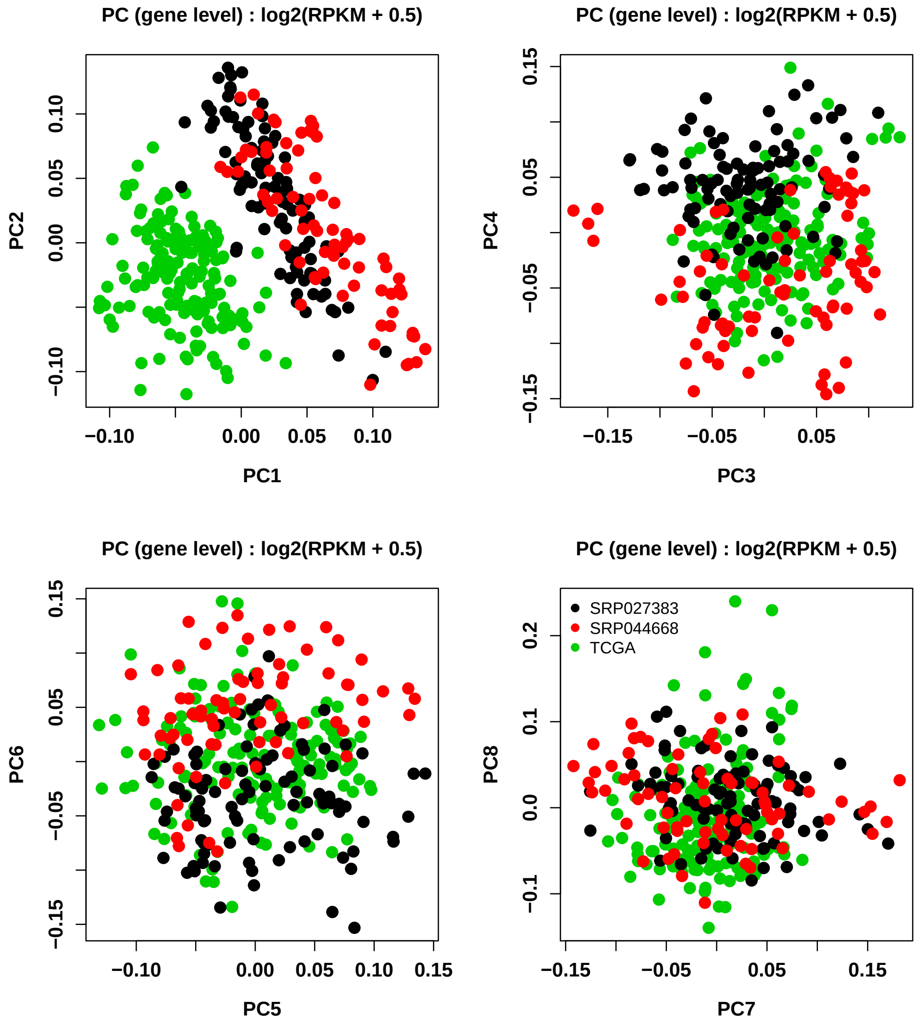

```
#Plot Variance Explained
var_plot(pca=expr.pca.counts.log2)
```

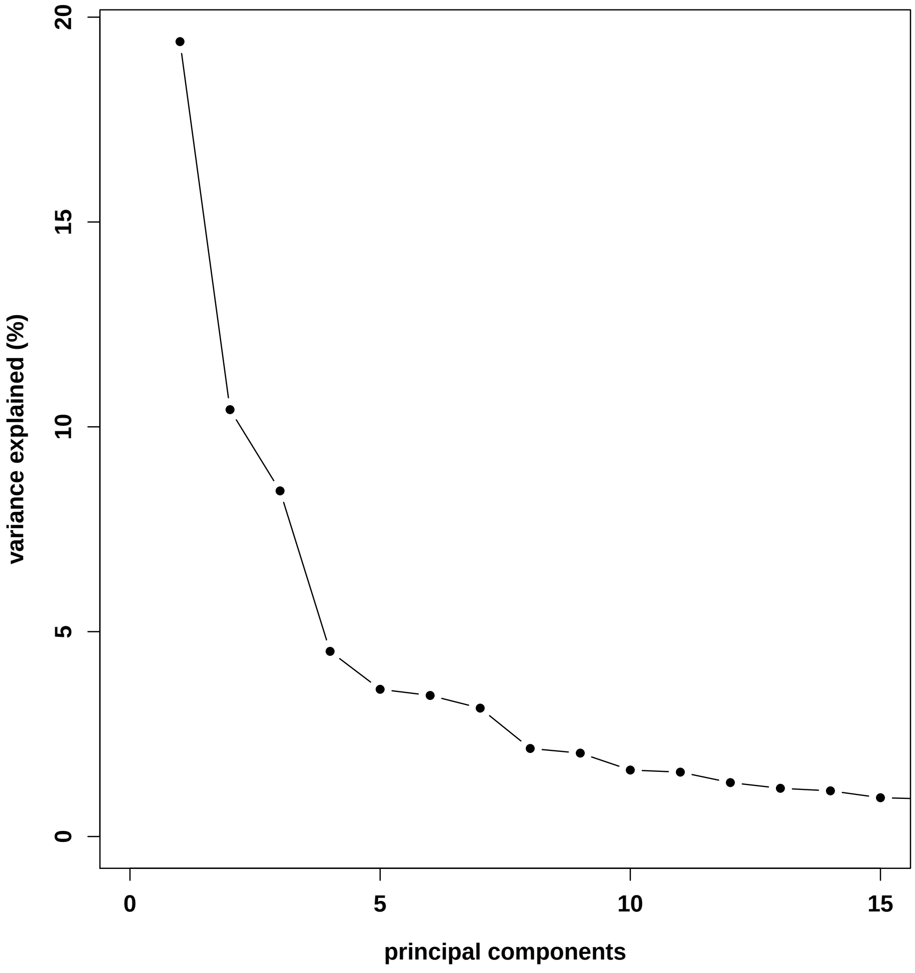

#### 3.4 Normalize for Dataset

To remove effects of TCGA vs SRA data sets (as they separate out in initial PCA), we first adjust the expression data and remove the effects of dataset. As TCGA is separating out in PC1 above while the two SRA samples are not, we code dataset as 1 if TGCA and 0 otherwise.

```
## code dataset where SRA dataset is 0 and TCGA is 1
dataset <- md$dataset
dataset[md$dataset=="SRP027383"] <- 0
dataset[md$dataset=="SRP044668"] <- 0
dataset[md$dataset=="TCGA"] <- 1
dataset <- as.numeric(dataset)

## function to normalize expression removing effects of dataset
normalize <- function(x){
  #run model
  lmfit <- lm(x ~ dataset)
  #adjust data
  out <- x -  (coef(lmfit)["dataset"]*(dataset))
```

```
    return(out)
  }

  df <- apply(rpkm_log2,1,normalize)
  rpkm_normalized <- t(df)

  expr.pca.normalized <- pc_function(rpkm_normalized)

  pc_plot(pca=expr.pca.normalized,legend=md$dataset, color=md$dataset,
          main="PC (gene level) : log2(normalized(RPKM) + 0.5)",
          type='variable')
```

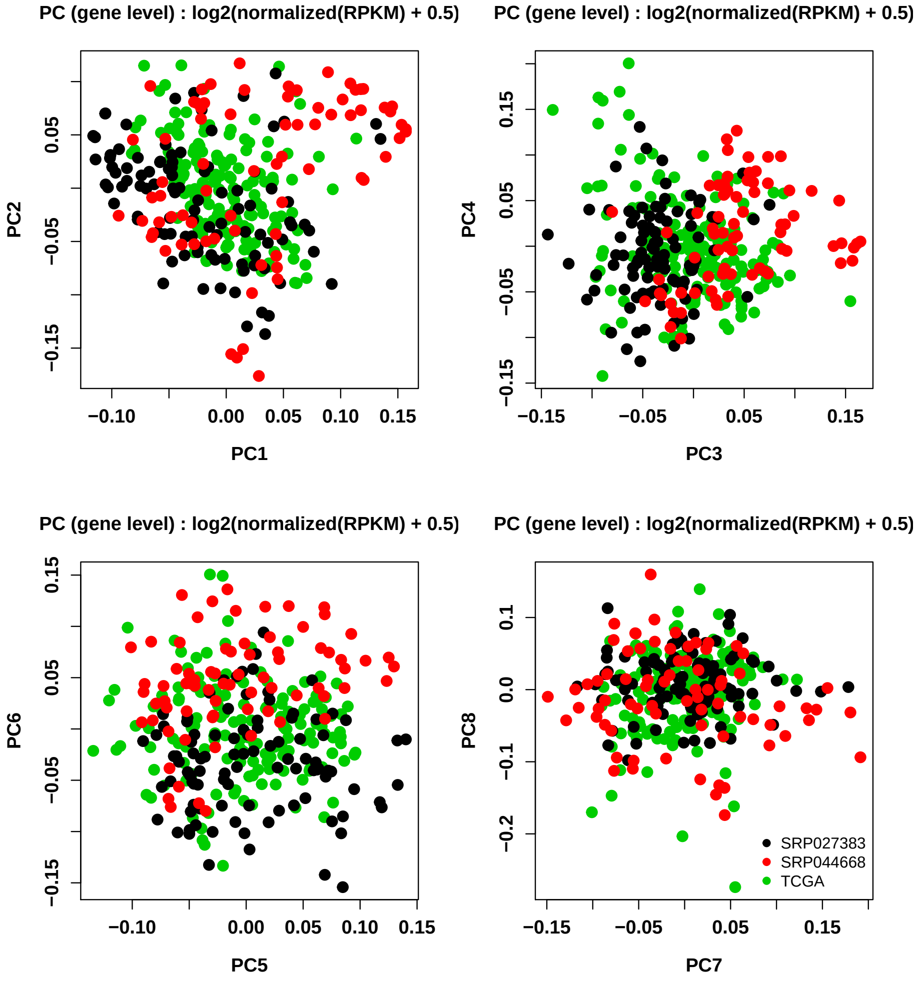

```
  ## Plot Variance Explained
  var_plot(pca=expr.pca.normalized)
```

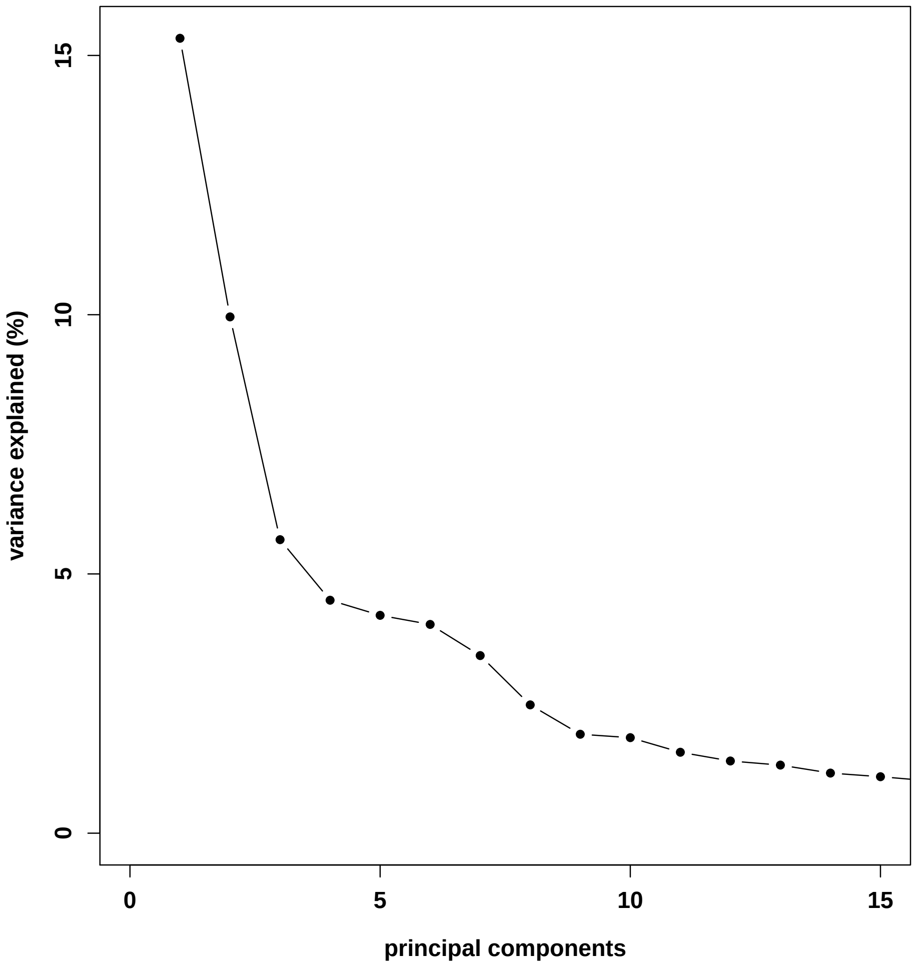

#### 3.5 Adjust for PCs

Above we see that removing the effects of dataset is helpful, however, the data do not completely overlap. To address this, we then remove the effects of the first 6 PCs from the expression data. After running PCA on these normalized expression data, we see that the effects of dataset have been removed and cross-study comparisons are now possible.

```
PC1 <- expr.pca.normalized$v[, 1]
PC2 <- expr.pca.normalized$v[, 2]
PC3 <- expr.pca.normalized$v[, 3]
PC4 <- expr.pca.normalized$v[, 4]
PC5 <- expr.pca.normalized$v[, 5]
PC6 <- expr.pca.normalized$v[, 6]

normalize_PC <- function(x){
  #run model
  lmfit <- lm(x ~  + PC1 + PC2 + PC3 + PC4 + PC5 + PC6 )
  #adjust data
  out <- x - (coef(lmfit)["PC1"]*(PC1-mean(PC1))) - (coef(lmfit)["PC2"]*(PC2-mean(PC2))) -
```

#### recount_brain cross-study example

```
    (coef(lmfit)["PC3"]*(PC3-mean(PC3))) - (coef(lmfit)["PC4"]*(PC4-mean(PC4))) -
    (coef(lmfit)["PC5"]*(PC5-mean(PC5))) - (coef(lmfit)["PC6"]*(PC6-mean(PC6)))
  return(out)
}

df <- apply(rpkm_normalized,1,normalize_PC)
rpkm_normalized_PC <- t(df)

## Run PCA
expr.pca.normalized.pc <- pc_function(rpkm_normalized_PC)

## Plot PCA
pc_plot(pca=expr.pca.normalized.pc,legend=md$dataset, color=md$dataset,
        main="PC (gene level) : log2(normalized(RPKM) + 0.5)",
        type='variable')
```

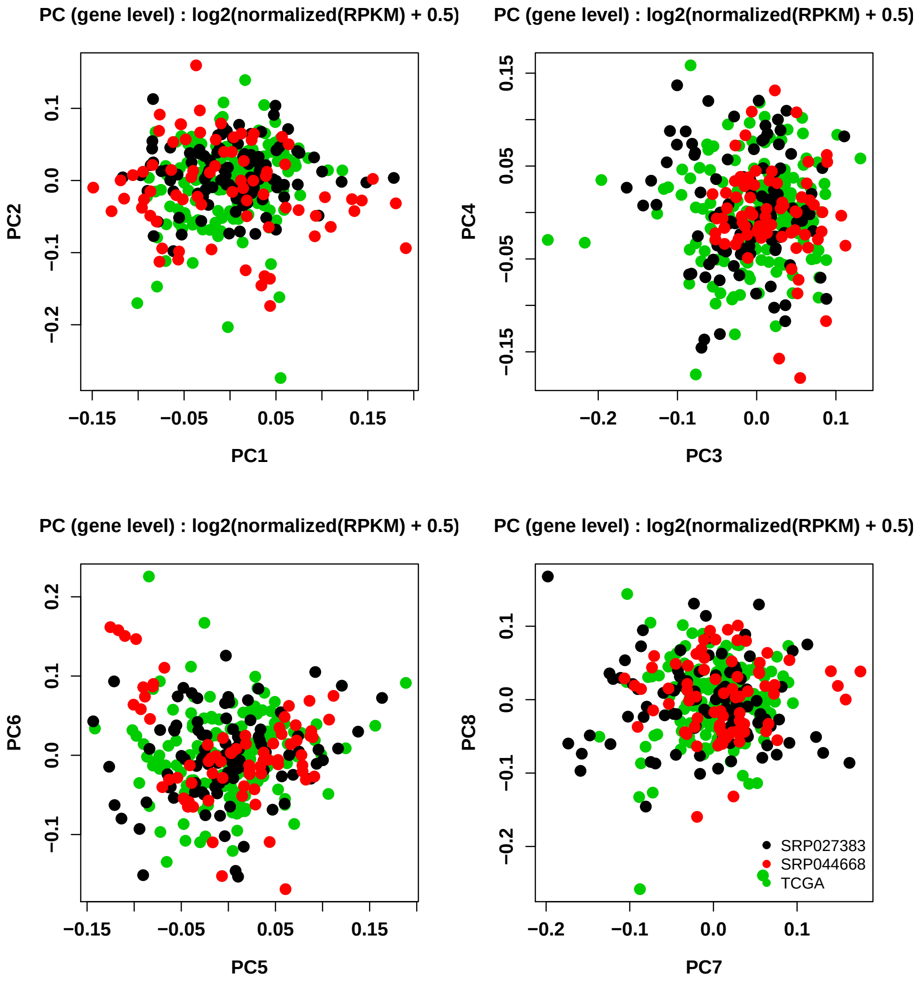

```
## Plot Variance Explained
var_plot(pca=expr.pca.normalized.pc)
```

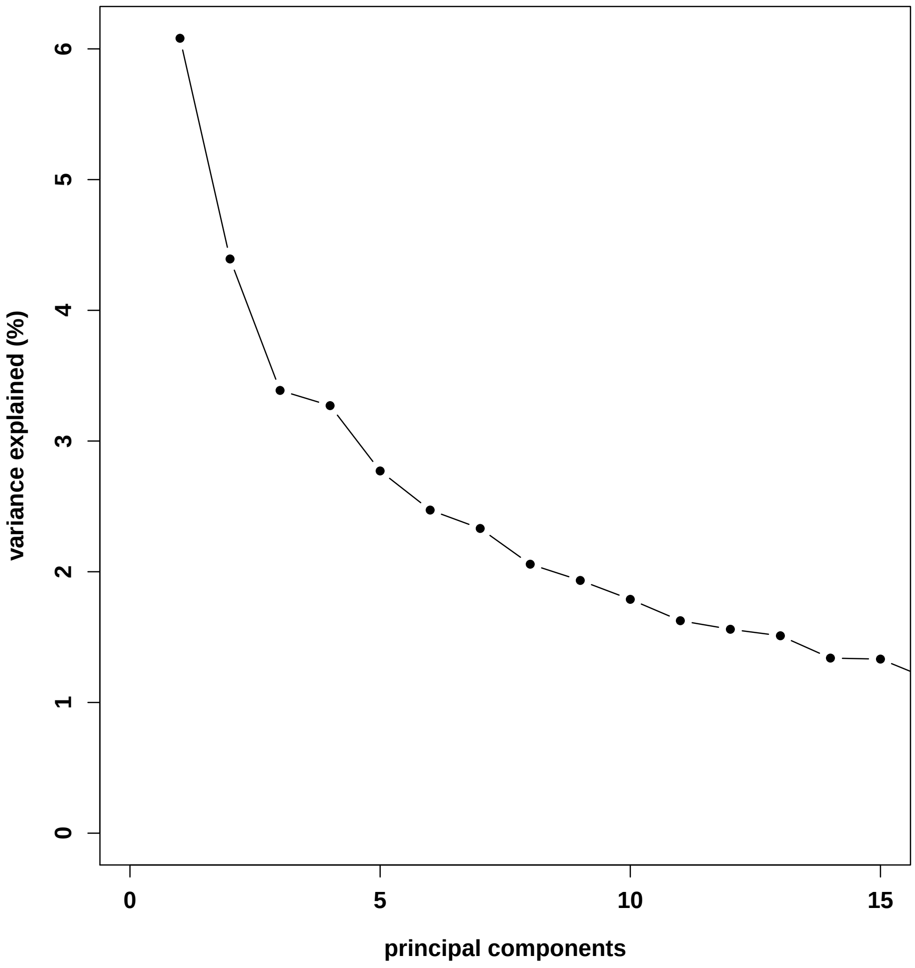

### 4 Glioblastoma Data Analysis

Having cleaned the data and made cross-study comparisons possible, we're now interested in answering our question of interest. Are the same genes most variable across different GBM studies?

#### 4.1 Variable Expression Analysis

To first answer this question, we calculate variance for each gene within each study.

```
df = as.data.frame(t(rpkm_normalized_PC))

## expression from each dataset
## N=74
SRP044668_df <- df %>%
  filter(md$dataset == 'SRP044668')
```

```
## N=175
SRP027383_df <- df %>%
  filter(md$dataset == 'SRP027383')

## N=270
TCGA_df <- df %>%
  filter(md$dataset == 'TCGA')

## is measure variance across each dataset?
SRP044668_vars <- colVars(as.matrix(SRP044668_df))
SRP027383_vars <- colVars(as.matrix(SRP027383_df))
TCGA_vars <- colVars(as.matrix(TCGA_df))

## add genenames back in
names(SRP044668_vars) <- names(SRP027383_vars) <- names(TCGA_vars) <- colnames(df)
```

#### 4.2 Concordance across studies

Having calculated within-study variance, we can then look at concordance at the top (CAT) plots to assess the results. To generate a CAT plot, we sort each study's genes by variance. Then, we compare the genes found in cross-study comparison to one another. If the analyses find the same genes, the line in the CAT plot will fall along the 45 degree (grey) line. The less concordant the results are, the further from this 45 degree line, the results will fall.

```
## sort by variance
p.mod1.sort <- SRP044668_vars[order(SRP044668_vars,decreasing=TRUE)]
p.mod2.sort <- SRP027383_vars[order(SRP027383_vars,decreasing=TRUE)]
p.mod3.sort <- TCGA_vars[order(TCGA_vars,decreasing=TRUE)]

conc <- NULL
conc_TCGA <- NULL
conc_TCGA2 <- NULL

for(i in 1:length(p.mod2.sort)){
    conc[i] <- sum(names(p.mod2.sort)[1:i] %in% names(p.mod1.sort)[1:i])
    conc_TCGA[i] <- sum(names(p.mod2.sort)[1:i] %in% names(p.mod3.sort)[1:i])
    conc_TCGA2[i] <- sum(names(p.mod1.sort)[1:i] %in% names(p.mod3.sort)[1:i])
}

## all genes
par(mfrow = c(1, 1), font.lab = 1.5, cex.lab = 1.2, font.axis = 1.5, cex.axis = 1.2)
plot(seq(1:length(p.mod2.sort)), conc,
     type = 'l', las = 0,
     xlim = c(0, length(conc)),
     ylim = c(0, length(conc)),
     xlab = 'ordered genes in reference study',
     ylab = 'ordered genes in new study',
     main = 'Concordance')
for(k in 1:30){
```

```r
    abline(v = k * 5000, cex = 0.5, col = 'lightgrey')
    abline(h = k * 5000, cex = 0.5, col = 'lightgrey')
}
abline(coef=c(0,1),col="grey48")
lines(seq(1:length(p.mod2.sort)), conc, col = bright[2], lwd = 3)
lines(seq(1:length(p.mod3.sort)), conc_TCGA, col = bright[5], lwd = 3)
lines(seq(1:length(p.mod3.sort)), conc_TCGA2, col = bright[8], lwd = 3)
legend('topleft', col = bright[c(2,5,8)],
       c("SRP044668_SRP027383",  "SRP027383_TCGA","SRP044668_TCGA"),
       lty=1,lwd=5, bg="white",bty="n")
```

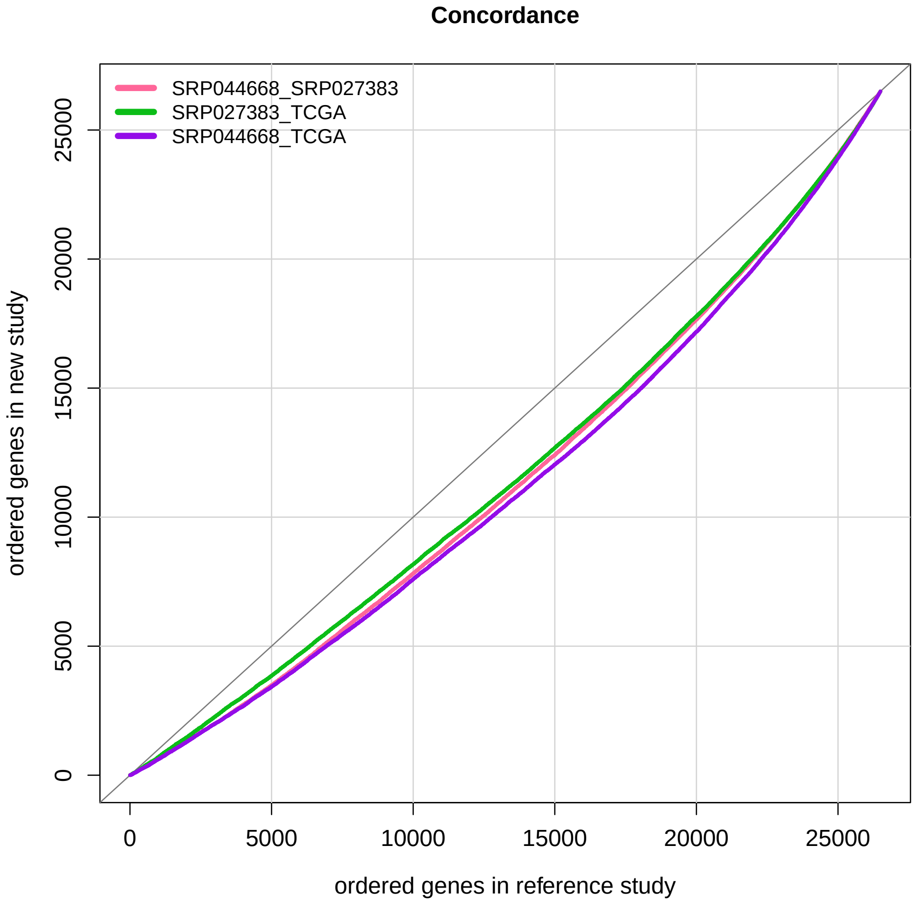

```r
## top 1000 genes
par(mfrow = c(1, 1), font.lab = 1.5, cex.lab = 1.2, font.axis = 1.5, cex.axis = 1.2)
plot(seq(1:1000), conc[1:1000],
     type = 'l', las = 0,
     xlim = c(0, 1000),
     ylim = c(0, 1000),
     xlab = 'ordered genes in reference study',
     ylab = 'ordered genes in new study',
     main = 'Concordance')
for(k in 1:10){
```

```
    abline(v = k * 200, cex = 0.5, col = 'lightgrey')
    abline(h = k * 200, cex = 0.5, col = 'lightgrey')
}
abline(coef=c(0,1),col="grey48")
lines(seq(1:1000), conc[1:1000], col = bright[2], lwd = 3)
lines(seq(1:1000), conc_TCGA[1:1000], col = bright[5], lwd = 3)
lines(seq(1:1000), conc_TCGA2[1:1000], col = bright[8], lwd = 3)
legend('topleft', col = bright[c(2,5,8)],
       c("SRP027383_SRP044668", "SRP027383_TCGA","SRP044668_TCGA"),
       lty=1,lwd=5, bg="white",bty="n")
```

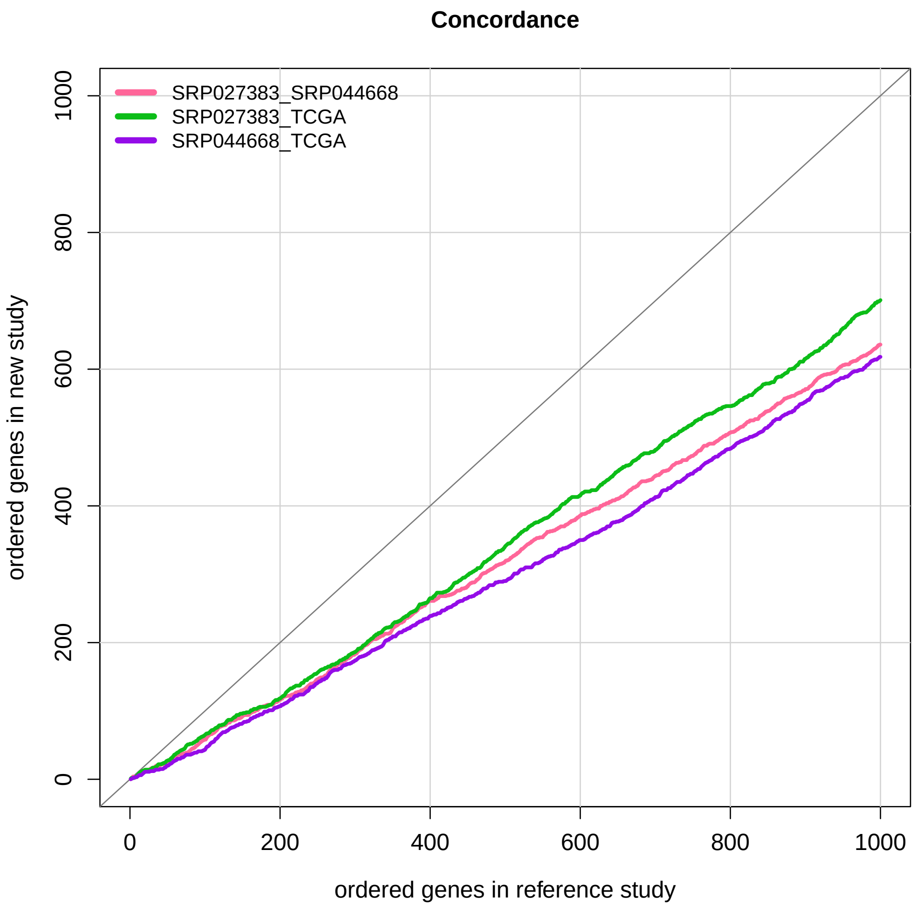

```
## most variable genes overlap across datasets?
```

Here, wee see that each cross-study comparison shows a similar level of concordance between studies.

### 5 Overlap with non-GBM samples

To compare these results to a non-GBM cancer study, we download TCGA kidney cancer data from `recount`.

#### 5.1 Data

The data here are kidney primary tumor samples. We chose kidney because there are a relatively large number of available samples and because this tissue is biologically dissimilar from brain.

```
# take a look at what samples we have normal tissue for
tcga %>% group_by(gdc_cases.project.primary_site,cgc_sample_sample_type) %>%
  summarise(n=n()) %>%
  filter(cgc_sample_sample_type=="Solid Tissue Normal") %>%
  arrange(-n)
## # A tibble: 20 x 3
## # Groups:   gdc_cases.project.primary_site [20]
##    gdc_cases.project.primary_site cgc_sample_sample_type     n
##    <chr>                          <chr>                  <int>
##  1 Kidney                         Solid Tissue Normal      129
##  2 Breast                         Solid Tissue Normal      112
##  3 Lung                           Solid Tissue Normal      110
##  4 Thyroid                        Solid Tissue Normal       59
##  5 Prostate                       Solid Tissue Normal       52
##  6 Colorectal                     Solid Tissue Normal       51
##  7 Liver                          Solid Tissue Normal       50
##  8 Head and Neck                  Solid Tissue Normal       44
##  9 Stomach                        Solid Tissue Normal       37
## 10 Uterus                         Solid Tissue Normal       35
## 11 Bladder                        Solid Tissue Normal       19
## 12 Esophagus                      Solid Tissue Normal       13
## 13 Bile Duct                      Solid Tissue Normal        9
## 14 Brain                          Solid Tissue Normal        5
## 15 Pancreas                       Solid Tissue Normal        4
## 16 Adrenal Gland                  Solid Tissue Normal        3
## 17 Cervix                         Solid Tissue Normal        3
## 18 Soft Tissue                    Solid Tissue Normal        2
## 19 Thymus                         Solid Tissue Normal        2
## 20 Skin                           Solid Tissue Normal        1

# can see that there are a lot of kidney healthy and kidney tumor.
# Will compare these to GBM with hypothesis that GBM most similar
# to kidney and less similar to healthy but overall less similar to GBM comparison

if(!file.exists(file.path('TCGA', 'rse_gene_kidney.Rdata'))) {

dir.create('TCGA', showWarnings = FALSE)
  downloader::download('http://duffel.rail.bio/recount/v2/TCGA/rse_gene_kidney.Rdata', destfile =
                'TCGA/rse_gene_kidney.Rdata', mode = 'wb')
}
load(file.path('TCGA','rse_gene_kidney.Rdata'))
assign('rse_gene_TCGA_kidney', rse_gene)

use <- colData(rse_gene_TCGA_kidney)$cgc_sample_sample_type=="Primary Tumor"
rse_gene_TCGA_kidney <- rse_gene_TCGA_kidney[,use]
```

```
load(file.path('TCGA','rse_gene_brain.Rdata'))
md_t <- colData(rse_gene)

rse_combine_TCGA <- rse_gene_TCGA
tokeep2 <- (md_t$gdc_cases.project.name=="Glioblastoma Multiforme" &
              md_t$cgc_sample_sample_type=="Primary Tumor")
md_t<-md_t[tokeep,]
colData(rse_combine_TCGA) <- md_t

rse_TCGA <- cbind(rse_combine_TCGA, rse_gene_TCGA_kidney)
md_TCGA <- colData(rse_TCGA)
```

#### 5.2 Filter genes (low expression)

We again filter out lowly-expressed genes. Here we include the same genes for analysis as were used in the analysis above.

```
## remove lowly expressed genes
rse_rpkm_tcga <- getRPKM(scale_counts(rse_TCGA))

## Filter genes with low levels of expression (same genes as initial analysis)
rpkm_tcga <- rse_rpkm_tcga[rpkm_mean > round(mean(expr_cuts), 2),]
rpkm_log2_tcga <- log2(rpkm_tcga+0.5)
```

#### 5.3 Run PCA

We again assess global gene expression patterns using PCA. We see that samples are clustering by tissue, as expected.

```
## Run PCA
expr.pca.rpkm.log2.tcga <- pc_function(rpkm_log2_tcga)

## Plot PCA

colors <- rep("green" , length(md_TCGA$gdc_cases.project.primary_site))
colors[md_TCGA$gdc_cases.project.primary_site=="Brain"] <- bright["purple"]

pc_plot(pca=expr.pca.rpkm.log2.tcga,legend=md_TCGA$gdc_cases.project.primary_site,
        color=colors,
        main="PC (gene level) : log2(RPKM + 0.5)",
        ptsize=0.9, position="topright",
        type='character')
```

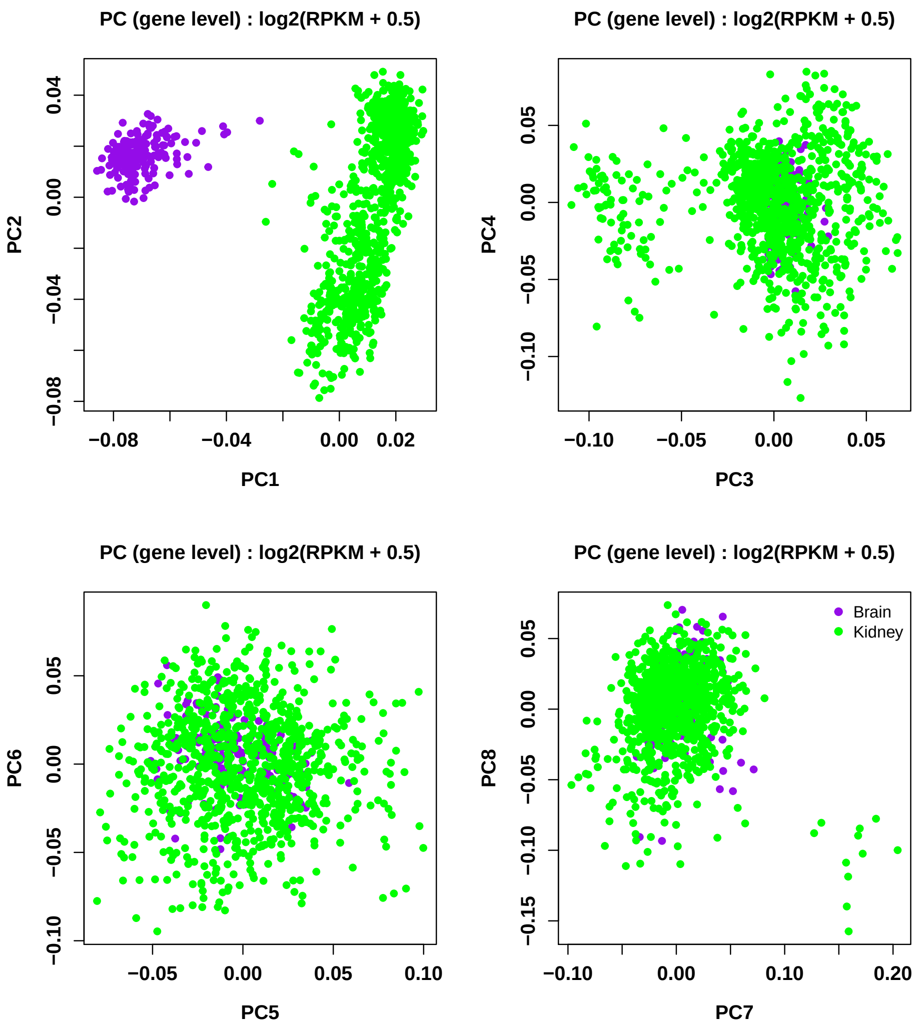

```
## Plot Variance Explained
var_plot(pca=expr.pca.rpkm.log2.tcga)
```

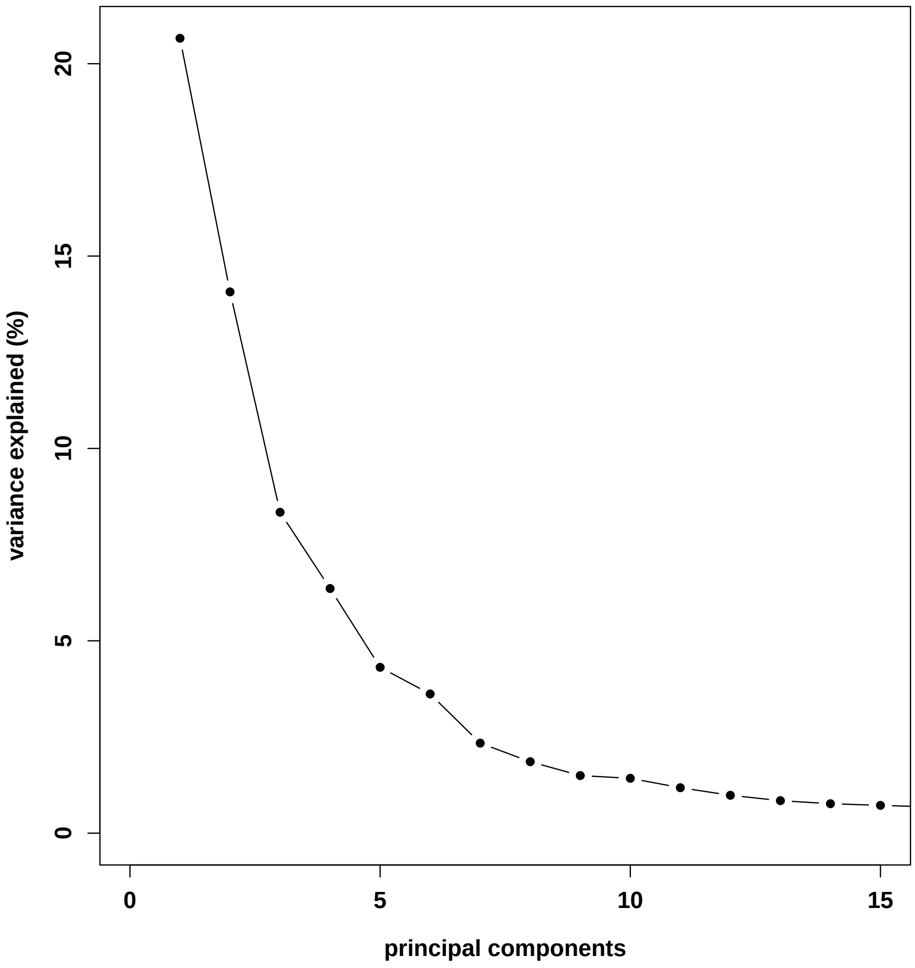

#### 5.4 Adjust for tissue

Looking at the separation across PCs1-4 above, we remove the effects of these PCs from the expression data before proceeding.

```
PC1 <- expr.pca.rpkm.log2.tcga$v[, 1]
PC2 <- expr.pca.rpkm.log2.tcga$v[, 2]
PC3 <- expr.pca.rpkm.log2.tcga$v[, 3]
PC4 <- expr.pca.rpkm.log2.tcga$v[, 4]

## function to normalize expression removing effects of dataset
normalize_kidney <- function(x){
  #run model
  lmfit <- lm(x ~  PC1 + PC2+ PC3 + PC4)
  #adjust data
  out <- x - (coef(lmfit)["PC1"]*(PC1-mean(PC1))) -
    (coef(lmfit)["PC2"]*(PC2-mean(PC2))) - (coef(lmfit)["PC3"]*(PC3-mean(PC3))) -
    (coef(lmfit)["PC4"]*(PC4-mean(PC4)))
  return(out)
```

```
}
df <- apply(rpkm_log2_tcga,1,normalize_kidney)
rpkm_normalized_tcga <- t(df)

## Run PCA
expr.pca.normalized.tcga <- pc_function(rpkm_normalized_tcga)

## Plot PCA
pc_plot(pca=expr.pca.normalized.tcga,legend=md_TCGA$gdc_cases.project.primary_site,
        color=colors,
        main="PC (gene level) : log2(RPKM + 0.5)",
        ptsize=0.9, position="topright",
        type='character')
```

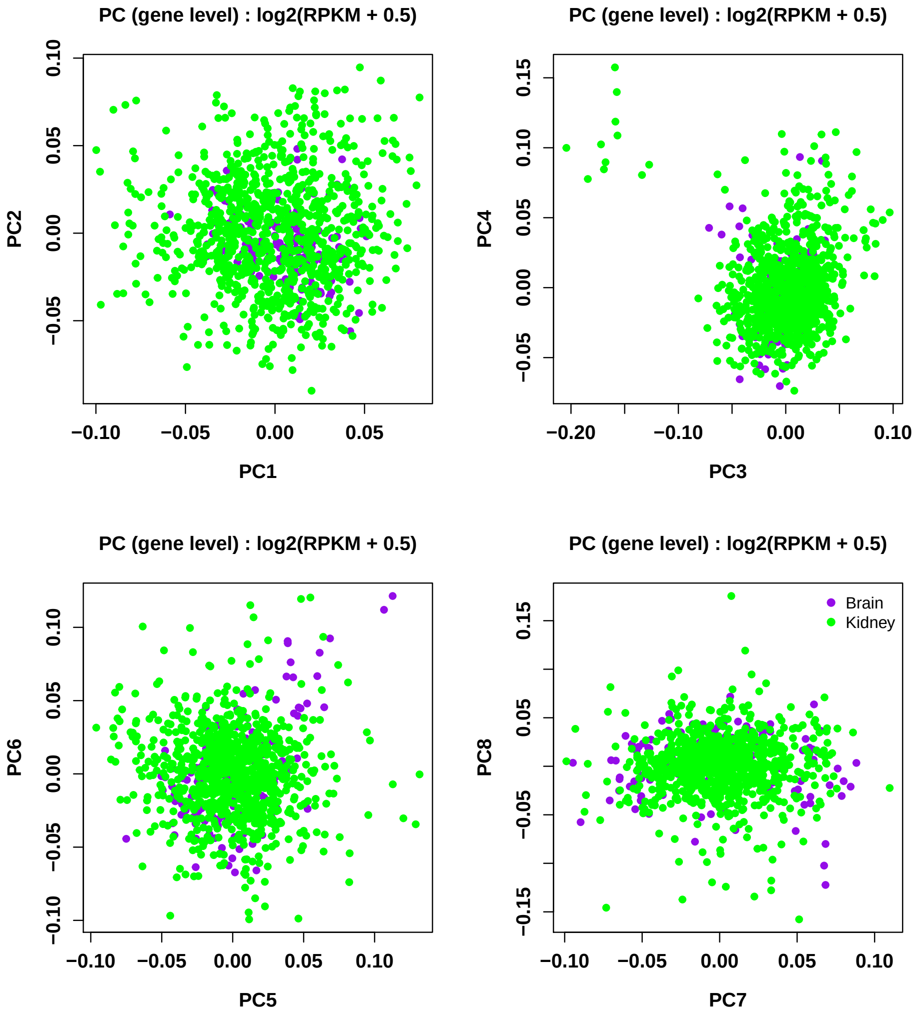

```
## Plot Variance Explained
var_plot(pca=expr.pca.normalized.tcga)
```

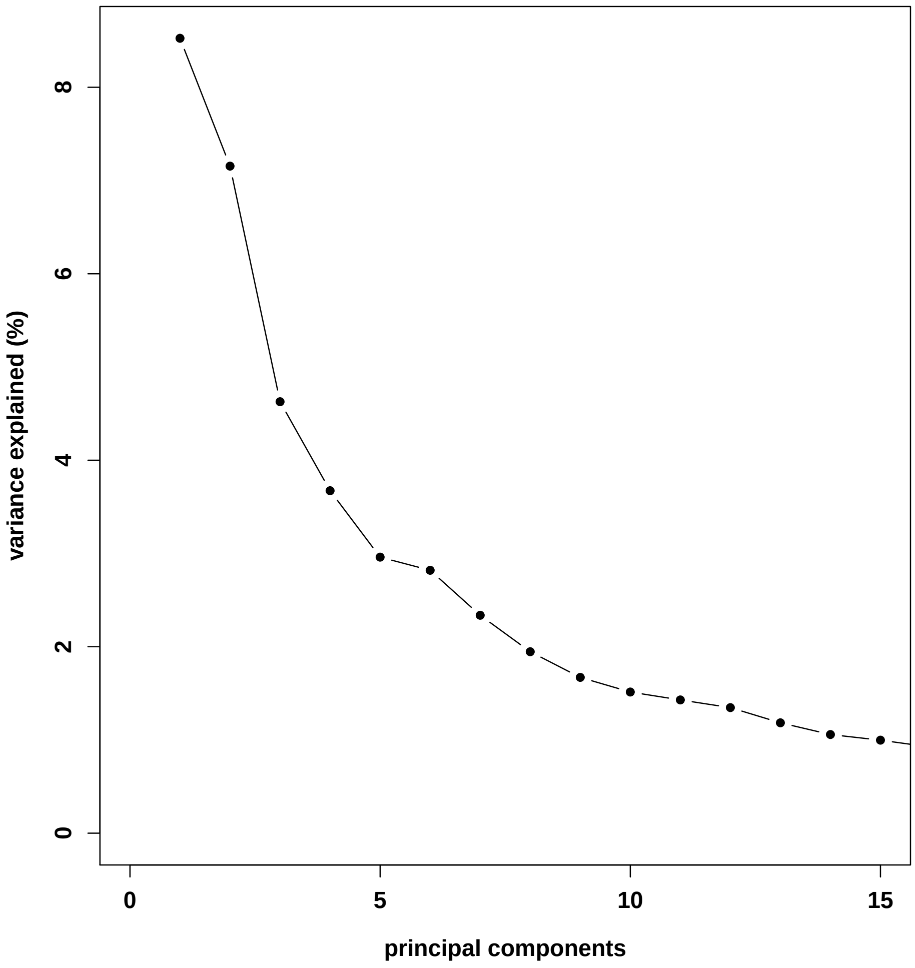

#### 5.5 Variable Expression Analysis

With expression data that can be compared across studies, we can then calculate variance across genes within each dataset.

```
df = as.data.frame(t(rpkm_normalized_tcga))

## expression from each dataset

# N=899
kidney_df <- df %>%
  filter(md_TCGA$gdc_cases.project.primary_site == 'Kidney')

brain_df <- df %>%
  filter(md_TCGA$gdc_cases.project.primary_site == 'Brain')

## is measure variance across each dataset?
kidney_vars <- colVars(as.matrix(kidney_df))
brain_vars <- colVars(as.matrix(brain_df))
```

```
## add genenames back in
names(kidney_vars)  <- names(brain_vars) <- colnames(df)
```

#### 5.6 Concordance across studies

Again, we look at concordance between the studies. Here, we're comparing TCGA's GBM data to TCGA's kidney tumor data. Concordance between these studies is included along with the previous concordance estimates.

```
## sort by variance
p.mod4.sort <- kidney_vars[order(kidney_vars,decreasing=TRUE)]
p.mod5.sort <- brain_vars[order(brain_vars,decreasing=TRUE)]

conc_kidney <- NULL
for(i in 1:length(p.mod2.sort)){
    conc_kidney[i] <- sum(names(p.mod5.sort)[1:i] %in% names(p.mod4.sort)[1:i])
}

## top 1000 genes
par(mfrow = c(1, 1), font.lab = 1.5, cex.lab = 1.2,
    font.axis = 1.5, cex.axis = 1.2)
plot(seq(1:1000), conc[1:1000],
     type = 'l', las = 0,
     xlim = c(0, 1000),
     ylim = c(0, 1000),
     xlab = 'ordered genes in reference study',
     ylab = 'ordered genes in new study',
     main = 'Concordance')
for(k in 1:10){
    abline(v = k * 200, cex = 0.5, col = 'lightgrey')
    abline(h = k * 200, cex = 0.5, col = 'lightgrey')
}
abline(coef=c(0,1),col="grey48")
lines(seq(1:1000), conc[1:1000], col = bright[2], lwd = 3)
lines(seq(1:1000), conc_TCGA[1:1000], col = bright[5], lwd = 3)
lines(seq(1:1000), conc_TCGA2[1:1000], col = bright[8], lwd = 3)
lines(seq(1:1000), conc_kidney[1:1000], col = bright[6], lwd = 3)
legend('topleft', col = bright[c(2,5,8,6)],
       c("SRP027383_SRP044668", "SRP027383_TCGA","SRP044668_TCGA", "brain_kidney"),
       lty=1,lwd=5, bg="white",bty="n")
```

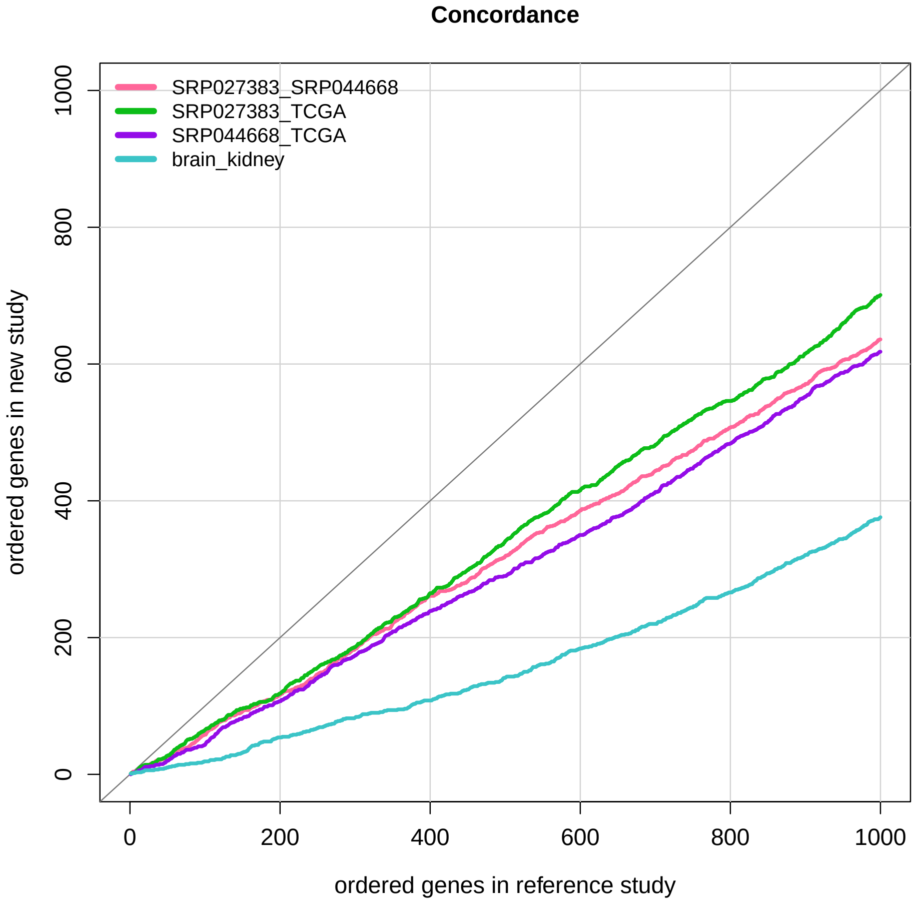

As expected, lower concordance is found between kidney and GBM data sets, suggesting that genes variable across GBM datasets are likely specific to GBM pathology.

### 6 Conclusions

Here, we demonstrate the utility of having well-curated metadata available from the `recount-brain` project. With confidence, we are able to identify three studies with relatively large sample sizes (N>=20) with samples whose disease pathology is overlapping. With three independent studies of samples from individuals with glioblastoma, we are able to normalize the data and then look to see if the same genes are found to be variable across the different studies. Additionally, we are able to assess whether overlap is lower among a different cancer type (kidney). Using this as a negative control, we see that, in fact, concordance across variable genes in glioblastoma appears to be consistent across studies. Following up on these concordantly variable genes could provide insight into glioblastoma pathology.

### Reproducibility

```
## Reproducibility information
Sys.time()
## [1] "2018-06-19 11:00:58 EDT"
proc.time()
##     user   system  elapsed
## 1096.177   97.148 1552.342
options(width = 120)
devtools::session_info()
## Session info ------------------------------------------------------------------------------------------------------------
##  setting  value
##  version  R version 3.5.0 (2018-04-23)
##  system   x86_64, darwin15.6.0
##  ui       X11
##  language (EN)
##  collate  en_US.UTF-8
##  tz       America/New_York
##  date     2018-06-19
## Packages ----------------------------------------------------------------------------------------------------------------
##  package                * version   date       source
##  acepack                  1.4.1     2016-10-29 CRAN (R 3.5.0)
##  AnnotationDbi            1.42.1    2018-05-08 Bioconductor
##  assertthat               0.2.0     2017-04-11 cran (@0.2.0)
##  backports                1.1.2     2017-12-13 cran (@1.1.2)
##  base                   * 3.5.0     2018-04-24 local
##  base64enc                0.1-3     2015-07-28 cran (@0.1-3)
##  bibtex                   0.4.2     2017-06-30 CRAN (R 3.5.0)
##  bindr                    0.1.1     2018-03-13 cran (@0.1.1)
##  bindrcpp               * 0.2.2     2018-03-29 cran (@0.2.2)
##  Biobase                * 2.40.0    2018-05-01 Bioconductor
##  BiocGenerics           * 0.26.0    2018-05-01 Bioconductor
##  BiocParallel           * 1.14.1    2018-05-06 Bioconductor
##  BiocStyle              * 2.8.2     2018-05-30 Bioconductor
##  biomaRt                  2.36.1    2018-05-24 Bioconductor
##  Biostrings               2.48.0    2018-05-01 Bioconductor
##  bit                      1.1-14    2018-05-29 CRAN (R 3.5.0)
##  bit64                    0.9-7     2017-05-08 CRAN (R 3.5.0)
##  bitops                   1.0-6     2013-08-17 cran (@1.0-6)
##  blob                     1.1.1     2018-03-25 CRAN (R 3.5.0)
##  bookdown                 0.7       2018-02-18 CRAN (R 3.5.0)
##  BSgenome                 1.48.0    2018-05-01 Bioconductor
##  bumphunter               1.22.0    2018-05-01 Bioconductor
##  checkmate                1.8.5     2017-10-24 CRAN (R 3.5.0)
##  cli                      1.0.0     2017-11-05 cran (@1.0.0)
##  cluster                  2.0.7-1   2018-04-13 CRAN (R 3.5.0)
##  codetools                0.2-15    2016-10-05 CRAN (R 3.5.0)
##  colorout               * 1.2-0     2018-05-03 Github (jalvesaq/colorout@c42088d)
##  colorspace               1.3-2     2016-12-14 cran (@1.3-2)
##  compiler                 3.5.0     2018-04-24 local
```

```
##  crayon                1.3.4     2017-09-16 cran (@1.3.4)
##  curl                  3.2       2018-03-28 CRAN (R 3.5.0)
##  data.table            1.11.4    2018-05-27 CRAN (R 3.5.0)
##  datasets            * 3.5.0     2018-04-24 local
##  DBI                   1.0.0     2018-05-02 CRAN (R 3.5.0)
##  DelayedArray        * 0.6.1     2018-06-15 Bioconductor
##  derfinder             1.14.0    2018-05-01 Bioconductor
##  derfinderHelper       1.14.0    2018-05-01 Bioconductor
##  devtools              1.13.5    2018-02-18 CRAN (R 3.5.0)
##  digest                0.6.15    2018-01-28 CRAN (R 3.5.0)
##  doRNG                 1.6.6     2017-04-10 CRAN (R 3.5.0)
##  downloader          * 0.4       2015-07-09 CRAN (R 3.5.0)
##  dplyr               * 0.7.5     2018-05-19 cran (@0.7.5)
##  evaluate              0.10.1    2017-06-24 cran (@0.10.1)
##  foreach               1.4.4     2017-12-12 CRAN (R 3.5.0)
##  foreign               0.8-70    2017-11-28 CRAN (R 3.5.0)
##  Formula               1.2-3     2018-05-03 CRAN (R 3.5.0)
##  GenomeInfoDb        * 1.16.0    2018-05-01 Bioconductor
##  GenomeInfoDbData      1.1.0     2018-05-03 Bioconductor
##  GenomicAlignments     1.16.0    2018-05-01 Bioconductor
##  GenomicFeatures       1.32.0    2018-05-01 Bioconductor
##  GenomicFiles          1.16.0    2018-05-01 Bioconductor
##  GenomicRanges       * 1.32.3    2018-05-16 Bioconductor
##  GEOquery              2.48.0    2018-05-01 Bioconductor
##  ggplot2               2.2.1     2016-12-30 CRAN (R 3.5.0)
##  glue                  1.2.0     2017-10-29 cran (@1.2.0)
##  graphics            * 3.5.0     2018-04-24 local
##  grDevices           * 3.5.0     2018-04-24 local
##  grid                  3.5.0     2018-04-24 local
##  gridExtra             2.3       2017-09-09 CRAN (R 3.5.0)
##  gtable                0.2.0     2016-02-26 CRAN (R 3.5.0)
##  Hmisc                 4.1-1     2018-01-03 CRAN (R 3.5.0)
##  hms                   0.4.2     2018-03-10 CRAN (R 3.5.0)
##  htmlTable             1.12      2018-05-26 CRAN (R 3.5.0)
##  htmltools             0.3.6     2017-04-28 cran (@0.3.6)
##  htmlwidgets           1.2       2018-04-19 CRAN (R 3.5.0)
##  httr                  1.3.1     2017-08-20 CRAN (R 3.5.0)
##  IRanges             * 2.14.10   2018-05-16 Bioconductor
##  iterators             1.0.9     2017-12-12 CRAN (R 3.5.0)
##  jaffelab            * 0.99.21   2018-05-03 Github (LieberInstitute/jaffelab@7ed0ab7)
##  janitor             * 1.0.0     2018-03-22 CRAN (R 3.5.0)
##  jsonlite              1.5       2017-06-01 CRAN (R 3.5.0)
##  knitcitations       * 1.0.8     2017-07-04 CRAN (R 3.5.0)
##  knitr                 1.20      2018-02-20 cran (@1.20)
##  lattice               0.20-35   2017-03-25 CRAN (R 3.5.0)
##  latticeExtra          0.6-28    2016-02-09 CRAN (R 3.5.0)
##  lazyeval              0.2.1     2017-10-29 CRAN (R 3.5.0)
##  limma                 3.36.1    2018-05-05 Bioconductor
##  locfit                1.5-9.1   2013-04-20 CRAN (R 3.5.0)
##  lubridate             1.7.4     2018-04-11 CRAN (R 3.5.0)
##  magrittr              1.5       2014-11-22 cran (@1.5)
```

```
##  Matrix                  1.2-14    2018-04-13 CRAN (R 3.5.0)
##  matrixStats           * 0.53.1    2018-02-11 CRAN (R 3.5.0)
##  memoise                 1.1.0     2017-04-21 CRAN (R 3.5.0)
##  methods               * 3.5.0     2018-04-24 local
##  munsell                 0.5.0     2018-06-12 CRAN (R 3.5.0)
##  nnet                    7.3-12    2016-02-02 CRAN (R 3.5.0)
##  parallel              * 3.5.0     2018-04-24 local
##  pillar                  1.2.3     2018-05-25 CRAN (R 3.5.0)
##  pkgconfig               2.0.1     2017-03-21 cran (@2.0.1)
##  pkgmaker                0.27      2018-05-25 CRAN (R 3.5.0)
##  plyr                    1.8.4     2016-06-08 cran (@1.8.4)
##  prettyunits             1.0.2     2015-07-13 CRAN (R 3.5.0)
##  progress                1.2.0     2018-06-14 CRAN (R 3.5.0)
##  purrr                   0.2.5     2018-05-29 cran (@0.2.5)
##  qvalue                  2.12.0    2018-05-01 Bioconductor
##  R6                      2.2.2     2017-06-17 CRAN (R 3.5.0)
##  rafalib               * 1.0.0     2015-08-09 CRAN (R 3.5.0)
##  RColorBrewer            1.1-2     2014-12-07 cran (@1.1-2)
##  Rcpp                    0.12.17   2018-05-18 cran (@0.12.17)
##  RCurl                   1.95-4.10 2018-01-04 CRAN (R 3.5.0)
##  readr                   1.1.1     2017-05-16 CRAN (R 3.5.0)
##  recount               * 1.6.2     2018-05-15 Bioconductor
##  RefManageR              1.2.0     2018-04-25 CRAN (R 3.5.0)
##  registry                0.5       2017-12-03 CRAN (R 3.5.0)
##  rentrez                 1.2.1     2018-03-05 CRAN (R 3.5.0)
##  reshape2                1.4.3     2017-12-11 cran (@1.4.3)
##  rlang                   0.2.1     2018-05-30 cran (@0.2.1)
##  rmarkdown               1.10      2018-06-11 CRAN (R 3.5.0)
##  rngtools                1.3.1     2018-05-15 CRAN (R 3.5.0)
##  rpart                   4.1-13    2018-02-23 CRAN (R 3.5.0)
##  rprojroot               1.3-2     2018-01-03 cran (@1.3-2)
##  Rsamtools               1.32.0    2018-05-01 Bioconductor
##  RSQLite                 2.1.1     2018-05-06 CRAN (R 3.5.0)
##  rstudioapi              0.7       2017-09-07 CRAN (R 3.5.0)
##  rtracklayer             1.40.3    2018-06-02 Bioconductor
##  S4Vectors             * 0.18.3    2018-06-08 Bioconductor
##  scales                  0.5.0     2017-08-24 cran (@0.5.0)
##  segmented               0.5-3.0   2017-11-30 CRAN (R 3.5.0)
##  splines                 3.5.0     2018-04-24 local
##  stats                 * 3.5.0     2018-04-24 local
##  stats4                * 3.5.0     2018-04-24 local
##  stringi                 1.2.3     2018-06-12 CRAN (R 3.5.0)
##  stringr                 1.3.1     2018-05-10 CRAN (R 3.5.0)
##  SummarizedExperiment * 1.10.1    2018-05-11 Bioconductor
##  survival                2.42-3    2018-04-16 CRAN (R 3.5.0)
##  tibble                  1.4.2     2018-01-22 cran (@1.4.2)
##  tidyr                   0.8.1     2018-05-18 cran (@0.8.1)
##  tidyselect              0.2.4     2018-02-26 cran (@0.2.4)
##  tools                   3.5.0     2018-04-24 local
##  utf8                    1.1.4     2018-05-24 CRAN (R 3.5.0)
##  utils                 * 3.5.0     2018-04-24 local
```

```
##  VariantAnnotation      1.26.0    2018-05-01 Bioconductor
##  withr                  2.1.2     2018-03-15 CRAN (R 3.5.0)
##  xfun                   0.2       2018-06-16 CRAN (R 3.5.0)
##  XML                    3.98-1.11 2018-04-16 CRAN (R 3.5.0)
##  xml2                   1.2.0     2018-01-24 CRAN (R 3.5.0)
##  xtable                 1.8-2     2016-02-05 CRAN (R 3.5.0)
##  XVector                0.20.0    2018-05-01 Bioconductor
##  yaml                   2.1.19    2018-05-01 CRAN (R 3.5.0)
##  zlibbioc               1.26.0    2018-05-01 Bioconductor
```

### References

The analyses were made possible thanks to:

- R (R Core Team, 2018)
- *BiocStyle*
- *clusterProfiler*
- *devtools* (Wickham, Hester, and Chang, 2018)
- *knitcitations* (Boettiger, 2017)
- *knitr* (Xie, 2014)
- *recount* (Collado-Torres, Nellore, Kammers, Ellis, et al., 2017; Collado-Torres, Nellore, and Jaffe, 2017)
- *rmarkdown* (Allaire, Xie, McPherson, Luraschi, et al., 2018)

Full bibliography file.

[1] J. Allaire, Y. Xie, J. McPherson, J. Luraschi, et al. rmarkdown: Dynamic Documents for R. R package version 1.10. 2018. URL: https://CRAN.R-project.org/package=rmarkdown.

[1] C. Boettiger. knitcitations: Citations for 'Knitr' Markdown Files. R package version 1.0.8. 2017. URL: https://CRAN.R-project.org/package=knitcitations.

[1] R Core Team. R: A Language and Environment for Statistical Computing. R Foundation for Statistical Computing. Vienna, Austria, 2018. URL: https://www.R-project.org/.

[1] H. Wickham, J. Hester and W. Chang. devtools: Tools to Make Developing R Packages Easier. R package version 1.13.5. 2018. URL: https://CRAN.R-project.org/package=devtools.

[1] Y. Xie. "knitr: A Comprehensive Tool for Reproducible Research in R". In: Implementing Reproducible Computational Research. Ed. by V. Stodden, F. Leisch and R. D. Peng. ISBN 978-1466561595. Chapman and Hall/CRC, 2014. URL: http://www.crcpress.com/product/isbn/9781466561595.
